# The logic of native enhancer-promoter compatibility and cell-type-specific gene expression variation

**DOI:** 10.1101/2022.07.18.500456

**Authors:** Takeo Narita, Yoshiki Higashijima, Sinan Kilic, Elina Maskey, Katrin Neumann, Chunaram Choudhary

## Abstract

Cis-regulatory enhancers are essential for differential expression of developmental and housekeeping genes. However, the specificity of native mammalian enhancers and how it shapes cell-type-specific gene expression landscapes remain largely unknown. We show that endogenous enhancers are broadly compatible with the promoters of developmental and housekeeping genes. Broad enhancer compatibility affords ‘retrofitting’ new regulatory capabilities to housekeeping genes that evolved before the advent of enhancers. This enables cell-type-specific tuning of ubiquitously expressed genes. Segregation between enhancer-dependent and –independent type regulation is blurred. Within the same cell type, a single promoter can be activated by enhancers and non-enhancer promoter-regulatory elements (PREs). It is the tunable and integrated strengths of enhancers and PREs that quantitatively shape gene expression landscapes, within and across cell types. Our findings have broad implications for understanding cell-type-specific quantitative gene expression variation, as well as the emergence and rewiring of gene regulatory networks in disease and organismal evolution.

## Introduction

Gene transcription in metazoans is controlled by promoters and enhancers^1^. An autonomous promoter consists of a core promoter (CP) and proximally occurring regulatory sequences (hereafter non-enhancer promoter regulatory elements; PREs). CPs define the transcription start site (TSS) and recruit the basal transcription machinery. PREs recruit transcription factors (TFs) and cofactors (COFs) and provide an activation signal to the CP. Many cell-type-specific (CTS) gene promoters lack potent PREs, and instead, are activated by enhancers, which act orientation and distance independently and can be located hundreds of kilobases away from the target CP^2, 3^. In mammals, enhancers outnumber genes, many enhancers skip their nearest genes and contact distally located promoters, multiple promoters interact with one enhancer, multiple enhancers interact with one promoter, and enhancers can coalesce to form phase-separated condensates^4–10^. A key question is, in such environments, how enhancers specifically activate some genes without affecting others.

In different analyses, enhancer-promoter (E-P) compatibility shows considerable differences. An early study reported that enhancers lack specificity and similarly activate diverse promoters^11^. Some of the recent analyses confirm this idea on a larger scale^12, 13^. However, in other analyses, enhancers show remarkable specificity and activate some but not other promoters^9, 14–17^. In massively parallel reporter assays (MPRAs), a majority of *Drosophila* and mouse enhancers show promoter specificity^18, 19^, and the compatibility information appears to be sequence-encoded in the enhancers and promoters^18, 20^. Supporting E-P specificity, different groups of human enhancers use different cofactors (COFs)^21^, and different COFs activate different promoters in *Drosophila*^22^. These findings present two seemingly diverging models of enhancer specificity. In one model, enhancers are broadly compatible with diverse promoters^11–,13^, while in another model, E-P compatibility is restricted such that enhancers activate selected promoters with high specificity^14–19^. An attractive aspect of the sequence-encoded E-P compatibility model is that it can rationalize why enhancers frequently skip their nearest genes and connect with distal genes^4–10^, how specificity can be achieved in phase-separated enhancer condensate hubs^23–25^, and how enhancers differentially activate alternative promoters within the same genes^22^.

One of the limitations is that the current understanding of E-P specificity is largely derived using plasmid-based ectopic reporters. Configurations of native cis-regulatory elements are far more complex, and arguably, endogenous enhancer specificity could be affected by many additional factors, such as 3D genome organization, chromatin modifications, competition among promoters, and binding of proteins that can facilitate or impede specific E-P interactions^2, 26–30^. Supporting this notion, a recent study identified thousands of candidate silencer elements in mammalian cells^31^, and 3D contact frequency suggests that the closest gene is often not the likely target of native enhancers^4, 32, 33^. Because of these complexities in the native environment, two fundamental questions remain unresolved. (1) Do enhancers display restricted or broad promoter specificity in the endogenous context? (2) What is the basis of cell-type-specific gene expression variability, and how does E-P compatibility impact quantitative gene expression variation?

Here, we use a reporter-free approach to infer native E-P specificity on a genome-wide scale. Importantly, we use native E-P compatibility principles to explain how E-P compatibility shapes endogenous transcription landscapes.

## Results

### Considerations for studying native enhancer specificity

A major unresolved challenge in the field is to find generalizable principles of enhancer-dependent gene regulation in mammalian cells, in the native context, on a global scale, across diverse cell systems. We reasoned that it should be possible to infer native E-P compatibility if we could: (1) define the cell-type-specificity of the expressed genes, (2) map enhancers and define the distances between enhancers and their nearest active promoters, (3) estimate the strength of enhancers, and (4) comprehensively quantify the enhancer-dependence of gene expression. We further posited that, by performing these analyses across diverse cell types, it should be possible to quantitatively assess enhancer and PRE function in cell-type-specific gene expression variation.

CBP/p300-dependent enhancers constitute the largest and best-characterized group of mammalian enhancers. To our knowledge, it is the only known group of enhancers that can be mapped using chromatin marks, whose strength can be estimated (see below), and whose activity can be rapidly ablated on a global scale^34^. Even in plasmid-based reporter assay, CBP/p300 is uniquely required for the activity of natively accessible enhancers (**Extended Data Fig. 1; Supplemental note 1**). For these reasons, we used CBP/p300-dependent enhancers as a model for understanding the rules of enhancer-dependent transcription.

### Determination of cell-type-specificity of genes

To investigate enhancer specificity and cell-type-specific gene regulation, we generated three isogenic cell types by differentiating mouse embryonic stem cells (ESCs) into epiblast stem cells (EpiSCs) and neural progenitor cells (NPCs), and confirmed their identity by cell-type-specific gene expression (**Extended Data Fig. 2a**). To confirm the generality of the rules, we used six human cell lines: HCT116, SH-SY5Y, Jurkat, KG-1, HepG2, and K562. The latter two are also used as reference cell lines by the ENCODE consortium^35^. In each cell line, we performed RNA-seq to define the cell-type-specificity of the expressed genes. Similar to a previous study ^36^, the expressed genes were categorized as cell-type-specific (CTS), cell-type-enriched (CTE), housekeeping (HK), and not classified (NC) (**Fig. 1a**; see Methods). Within each category, genes from all cell lines were summed and shown in a separate category (Combined). In all cell lines, HK and CTE genes constituted a roughly equal portion (∼40% each), and CTS genes constituted 7-14% (**Fig. 1a**). A slightly lower fraction of CTS genes in human cancer cell lines possibly reflects a lack of specific cytokines in the culture media and downscaling of CTS genes in long-term cultured cancer cells^37^.

**Fig. 1.**
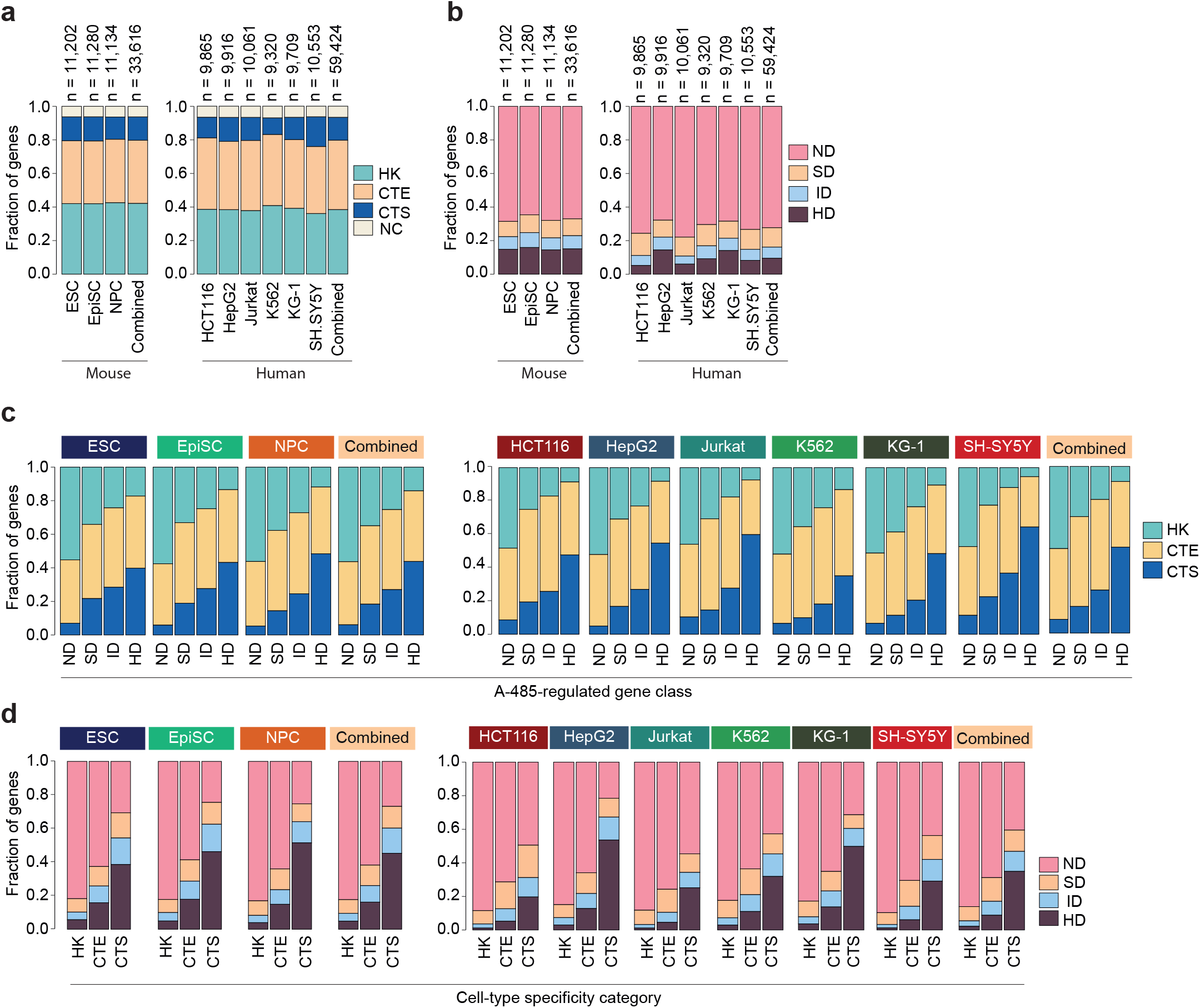
Enhancers prominently regulate cell-type-specific genes. **a,** Cell-type specificity of genes expressed in the indicat-use and human cell lines. HK, housekeeping; CTS, cell-type-specific; CTE, cell-type-enriched; NC, not classified into the three categories. In each cell line, the fraction of cell-type-specificity categories and the number (n) of expressed is indicated. Gene counts from all analyzed mouse cell lines and human cell lines are shown in a separate category bined). **b,** Fraction of A-485 downregulated genes in the indicated cell lines. HD, highly downregulated (≥2-fold ased); ID, intermediate downregulated (>1.5-2.0-fold decreased); SD, slightly downregulated (>1.2-1.5-fold decreased); D, not downregulated (≤1.2-fold decreased) after A-485 treatment. Gene quantified from all analyzed mouse cell lines uman cell lines are shown in a separate category (Combined). (Biological replicates of EU-seq: n=5 for ESC, n=3 for and NPC, n=2 for human cell lines). **c,** Fractional distribution of genes with the indicated cell-type-specificity categon the indicated cell lines and the depicted A-485-regulated gene categories. **d,** Fractional distribution of A-485-regulated gene class, in the indicated cell lines, and within the indicated cell-type-specificity categories.

### Quantification of enhancer-regulated genes

We used the CBP/p300 catalytic inhibitor A-485^38^ to comprehensively map the enhancer dependency of genes. A-485 specificity has been demonstrated using CBP/p300 knockout cells^39^. We further validated it by comparing the nascent transcription changes caused by A-485 and the CBP/p300 bromodomain-based degrader dCBP1^40^ (**Extended Data Fig. 2b**). A-485 and dCBP1-induced gene expression changes were highly correlated, and A-485 showed no sign of off-target specificity. A-485-induced catalytic inhibition occurs more rapidly (<5 min) than dCBP1-induced CBP/p300 depletion (>60 min)^34, 40^; therefore, we chose A-485 for this study. A-485-induced nascent transcription changes were measured using metabolic RNA labeling with 5-ethynyl uridine and sequencing (EU-seq). In ESC, EpiSC, and NPC, transcription was measured at three different time points (30, 60, and 120 min), with a minimum of three biological replicates, generating 9 independent quantification data points. A-485 caused rapid and selective downregulation of CTS genes (*Cobl*, *Enpp3*, *Npas3*), but not HK genes (*Mcmbp*, *Cct4*) (**Extended Data Fig. 3a**). In human cells, nascent transcription changes were quantified after 60 min of A-485 treatment, in two biological replicates. A-485 robustly downregulated enhancer target gene (*MYC*) without affecting non-target housekeeping gene (*CCT4*) (**Extended Data Fig. 3b**), confirming the suitability of our approach for separating enhancer-dependent and – independent genes.

### Enhancers regulate genes of diverse cell-type-specificity

Based on A-485-induced nascent transcription changes, genes were classified as highly downregulated (HD; decreased by ≥2-fold), intermediate downregulated (ID; decreased by 1.5-2.0-fold), slightly downregulated (SD; decreased by 1.2-1.5-fold), and not changed (NC; decreased by ≤1.2-fold) (**Fig. 1b**). Within each category, gene numbers from all cell lines were summed and shown as a separate category (Combined).

Next, we analyzed the relationship between enhancer dependence and cell-type specificity of genes. With increasing A-485-induced downregulation, the fraction of CTS genes progressively increased, while the fraction of HK genes gradually decreased (**Fig. 1c**). In cell-type-specificity categories, a larger fraction of CTS genes was HD than HK genes, and most HK genes were ND (**Fig. 1d**). The relationship between cell-type-specificity and A-485-induced gene downregulation is highly similar in diverse mouse and human cell lines. These results show that enhancers more frequently regulate CTS genes, but not all CTS genes are enhancer regulated. Conversely, most HK genes are activated enhancer-independently, but as such HK genes are not insensitive to enhancers.

### Validation of A-485-induced nascent transcription changes

CBP/p300 inhibition specifically impairs RNA Polymerase II (Pol2) recruitment and pause release at enhancer-regulated genes^34^. To independently validate EU-seq-based gene regulation, we quantified Pol2 binding in ESC, EpiSC, and NPC. A-485-induced changes in Pol2 binding were correlated (r=0.87-0.92) in replicate measurements (**Extended Data Fig. 4a**). Notably, the A-485-induced decrease in Pol2 binding was strongly correlated (r=0.80-0.86) with the decrease in nascent transcription (**Extended Data Fig. 4b**). Pol2 binding, both at TSS and gene body regions, was correspondingly reduced at EU-seq downregulated gene categories (**Extended Data Fig. 4c**). Robust, genome-scale correlation between the independent assays (nascent transcriptome analyses and Pol2 ChIP-seq), and across diverse types, strongly supports the quantitative accuracy of the measurements.

### Enhancers mainly regulate proximal genes

Next, we investigated the relationship between enhancer strength, E-P distance, cell-type-specificity, and A-485-induced gene regulation. Previously, accurate estimation of enhancer strength had remained challenging. In the accompanying manuscript, we established histone H2B N-terminus multisite lysine acetylation (H2BNTac) as a genuine signature of active enhancers (Narita et al.). We used H2BK20ac as a representative H2BNTac mark to map enhancers. For comparison, we also included H3K27ac, a widely used hallmark of active enhancers. H2BK20ac and H3K27ac occupancy were analyzed by ChIP-seq in all 9 cell lines (**Extended Data Fig. 5a**). Regions commonly marked by H2BK20ac and H3K27ac were considered as candidate enhancers, and enhancer strength was separately estimated using (1) H3K27ac ChIP-seq signal, or (2) H2BK20ac ChIP-seq signal.

To infer the rules of native E-P specificity, we investigated the relationship between the following four parameters: (1) the cell-type-specificity of genes, (2) the E-P distance, (3) the enhancer strength, and (4) A-485-induced gene downregulation. First, in each cell type, genes were grouped based on the cell-type specificity (CTS, CTE, HK). Then, within each cell-type-specificity category, genes were further grouped based on the E-P distance (0-10kb, 10-50kb, or >50kb). Next, within the above categories, genes were further sub-grouped based on the strength of proximal enhancers. Finally, in each of these categories, the fraction of A-485-regulated genes was determined.

H2BK20ac intensity-based enhancer ranking showed a strikingly clear relationship with all the above parameters (**Fig. 2a**, **Extended Data Fig. 5b**). In ESC, EpiSC, and NSC, A-485-induced gene downregulation directly reflected the strength of enhancers, their distance from nearest promoters, and the cell-type-specificity of genes. The strongest enhancers (Top10%, Top25%) had the greatest impact on the regulation of the nearest genes. Indeed, regardless of the cell type specificity of genes, most genes are downregulated if they occurred in proximity to strong enhancers (<10kb, Top10%). The linear distance also shows an unambiguous relationship with gene regulation. Within the same enhancer rank category, the extent of A-485-induced gene downregulation drops markedly if enhancers are located further away from the TSS (compare 0-10kb versus 10-50kb groups), indicating a clear relationship between enhancer distance and gene regulation (**Fig. 2a**). Unlike H2B20ac, H3K27ac-based ranking showed no clear relationship between enhancer strength, gene distance, and A-485-induced transcription downregulation in ESC, EpiSC, and NPC (**Fig. 2b**). In particular, CTE, HK, and Combined gene categories displayed no apparent relationship with H3K27ac-based enhancer ranking and gene regulation. These results demonstrate that the relationship between enhancer position, strength, cell-type-specificity and proximal gene regulation is far simpler and more predictable than suggested in the current literature. However, these relationships are poorly exposed by the widely used enhancer marker H3K27ac, and only become apparent when enhancer strength is defined by H2BK20ac. This may explain why these simple relationships have remained hidden in plain sight.

**Fig. 2.**
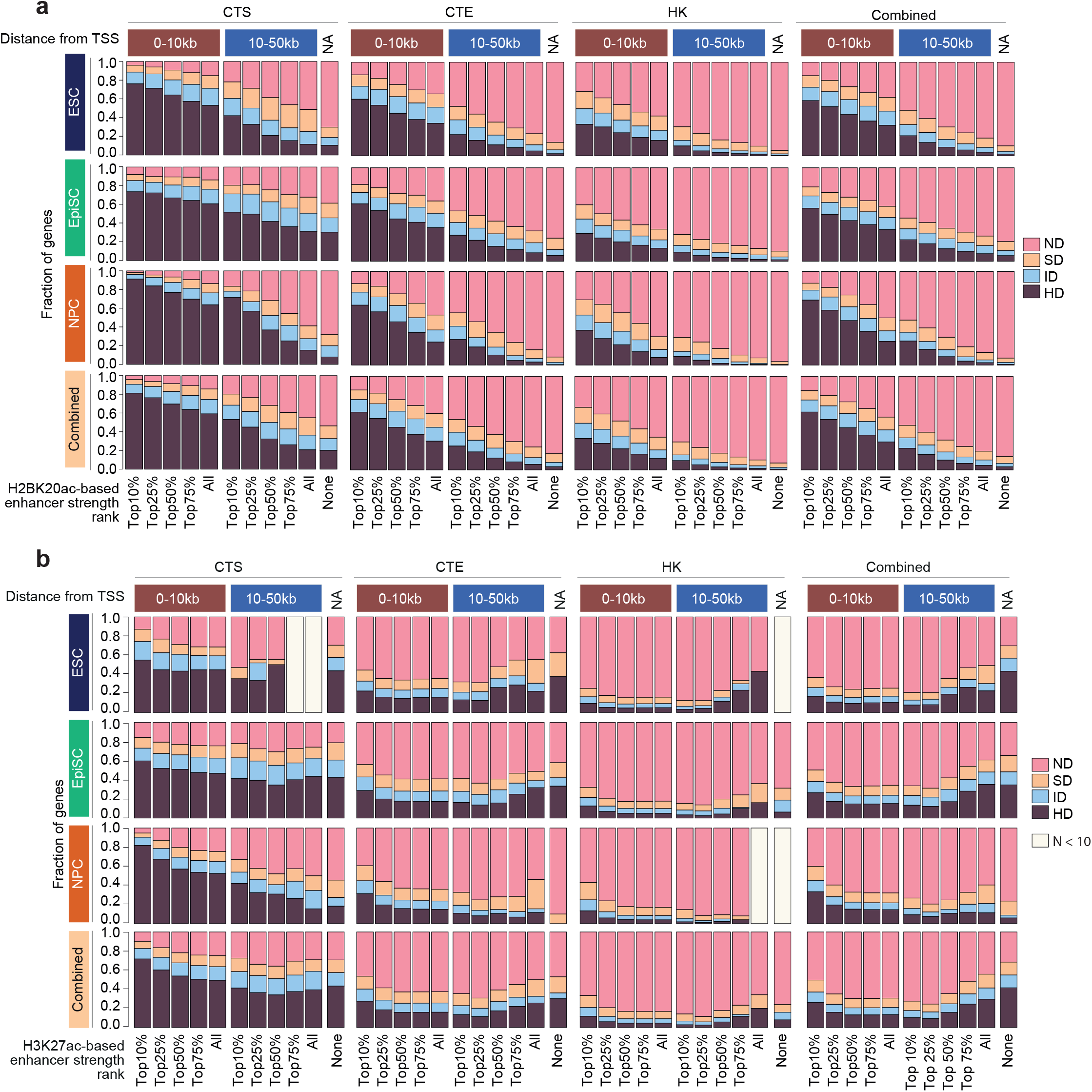
Enhancer-dependent gene regulation is strongly associated with enhancer strength, distance, and cell-type specificity of genes. **a,** Relationship between H2BK20ac intensity-based enhancer ranking and A-485-induced gene regulation. For the indicated cell lines, genes were grouped based on cell type specificity (CTS, CTE, HK, and Combined). Enhancers (marked with both H3K27ac + H2BK20ac) were ranked based on H2BK20ac ChIP intensity. Enhancers were rank-ordered and grouped into the indicated percentile categories. Genes were further grouped based on the shortest distance between TSS and the enhancer in each percentile category. Within different groups, the fraction of A-485 regulated gene categories are shown. All, all enhancers; None, no enhancer within 50kb from TSS. b, Relationship between H3K27ac intensity-based enhancer ranking and A-485-induced gene regulation. Enhancers used for these analyses are H3K27ac peak regions and enhancer strength was defined by the H3K27ac ChIP signal. The relationship between H3K27ac defined enhancer strength and distance with cell-type-specificity and A-485-induced gene regulation was analyzed as described in panel a. The groups where the number of genes is small (N< 10) are not shown.

### Enhancers are broadly compatible with diverse promoters

Of the all genes quantified in the mouse cell lines (n=33,616, **Fig. 1a**, Combined category), 1862 occurred in proximity to strong proximal enhancers (Top10%, <10kb), and 83% (HD 61%, ID 13%, SD 9%) of those were downregulated (**Fig. 2a**, Combined category). In contrast, 10152 genes lacked proximal enhancers (no enhancer within 50kb from TSS, NA category), and from those, only 14% (HD 3%, ID 4%, SD 7%) were downregulated. This shows that genes with strong proximal enhancers are more likely (odds ratio, 45, p < 2.2e-16, two-sided fisher test) to be HD than those lacking proximal enhancers. In ESC, EpiSC, and NPC, 60-70% of HK genes with strong proximal enhancers (Top10%, <10kb) are downregulated by A-485. In contrast, only 4-11% of HK genes lacking proximal enhancers are downregulated, demonstrating that HK genes occurring in proximity to strong enhancers are much more likely (odds ratio, ESC, 31, p < 2.2e-16; EpiSC, 13, p < 2.2e-16; NSC, 60, p < 2.2e-16, two-sided fisher test) to be downregulated than non-enhancer proximal genes. This difference would not be expected if HK CPs *per se* were insensitive to enhancers. If we consider a gene being enhancer-sensitive, even if it is only downregulated in one of the cell lines, about half of the expressed genes appear enhancer regulated by at least >1.2 fold (**Extended Data Fig. 5c**). Extrapolating these observations to hundreds of other cell types, it can be anticipated that a large majority of mammalian genes are impacted by enhancers in at least one cell type, even though in any one cell type, enhancers only regulate a small subset of genes.

To further understand enhancer compatibility, we manually inspected several genes in ESC that occurred proximally to strong enhancers (Top10%, <10kb) but were not downregulated by A-485. In most instances, the same enhancers occurred in proximity to other genes and regulated those genes. For example, *Rpl3* and *Polr2e* are insensitive to proximal enhancers. The same enhancers also localize in proximity to *Syngr1* and *Gpx4*, respectively, and regulate these genes, which occur more proximal to the enhancers (**Extended Data Fig. 6a**). Regulation of one of the proximal genes may explain a lack of regulation of the other proximal genes, which often localize more distally and have stronger promoters. Overall, of the 463 strong enhancer-proximal CTS and HK genes in ESC, ∼20 could be manually confirmed as enhancer insensitive, including *Eif4a2*, *Ube2s*, and *Zfp936* (**Extended Data Fig. 6b**). In these instances, no immediate enhancer-proximal genes were downregulated. Promoters of these genes could be incompatible with enhancers, but there can also be other reasons for their apparent insensitivity, such as long mRNA half-life or binding of insulators. Consistent with this idea, *MRPL23* is not regulated by the proximal enhancers because CTCF-Cohesin-mediated looping connects the enhancer to distally located *H19* and *IGF2*^41, 42^. *H19* and *IGF2* have short introns and very long mRNA half-life^43, 44^. Therefore, their nascent transcript levels are not reduced and they also appear enhancer insensitive. But a zoom-in view of the *IGF2* intron regions confirms that it is indeed downregulated in HepG2 (**Extended Data Fig. 6c**). *IL32* and *KCNQ* have proximal enhancers in more than one cell line yet show A-485 sensitivity in only one of them (**Extended Data Fig. 7**). This indicates that, as such, a lack of enhancer insensitivity in one cell line does not necessarily establish the insensitivity of a CP to enhancers. We conclude that enhancers have very broad, but not necessarily universal, compatibility with diverse promoters.

### Validation of enhancer-promoter compatibility

To confirm broad E-P compatibility, we used the *Nanog* upstream (-5kb) enhancer and positioned it in proximity to promoters of six different genes. The *Mcmbp*, *Sec23ip*, *Fbxo9*, *Ick*, and *Mthfsd* are broadly expressed and not regulated by A-485 in any of the analyzed cell lines in the native state (**Extended Data Fig. 8**). The *Foxc2* promoter is accessible in ESC, but the gene is marked with the repressive mark H3K27me3 and not expressed in the analyzed cell lines. The enhancer was stably inserted in the first intron of *Mcmbp*, *Sec23ip*, *Fbxo9*, and *Ick*. *Foxc2* is intronless, and thus, the enhancer was positioned downstream of the gene. For *Mthfsd*, the enhancer was inserted upstream of TSS. Reassuringly, in all cases, regardless of the cell-type-specificity of genes and orientation of the enhancer, enhancer insertion significantly (p=<0.0001) increased the expression of the proximal genes, supporting the idea of broad enhancer compatibility (**Fig. 3a**).

**Fig. 3.**
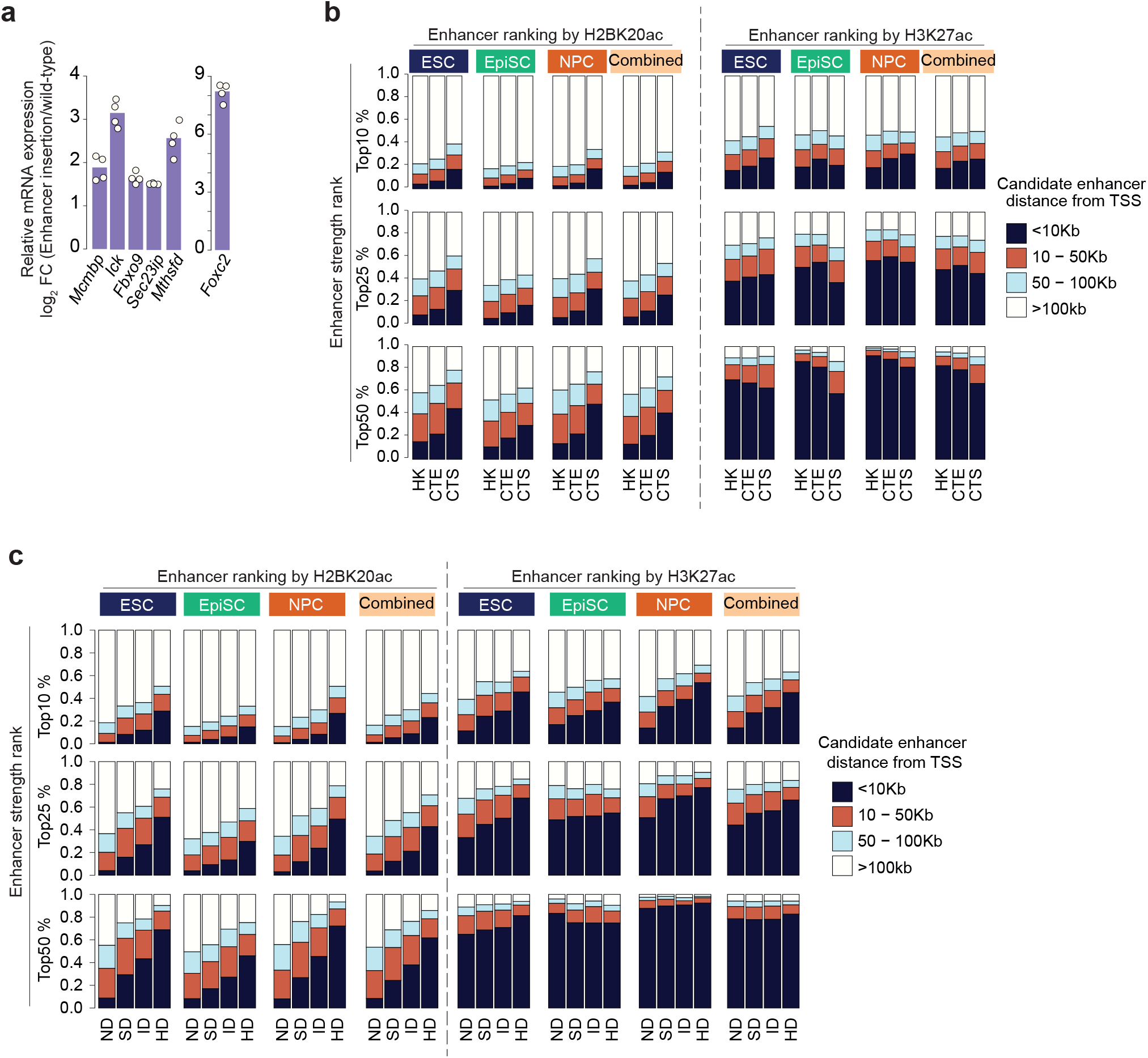
Strong enhancers frequently occur proximal to CTS and A-485 downregulated genes. **a,** Enhancer insertion increases expression of proximal genes. For five of the genes (*Mcmbp, Fbxo9, Ick, Sec23ip, Mthfsd*), the Nanog upstream (-5kb) enhancer was inserted in the first intron of the genes. Foxc2 is intronless, and hence the enhancer was positioned down-stream of the gene (see Methods). The relative mRNA expression was measured in wild-type and enhancer-inserted cells using quantitative RT-PCR. **b-c,** In the indicated mouse cell lines, the expressed genes were classified based on their cell type-specificity (b) or A-485-regulated gene class (c). Enhancer strength was determined using H2BK20ac (left panels) or H3K27ac (right panels) ChIP-signal, and enhancers were grouped into Top10%, 25%, and 50% categories. Only commonly marked H3K27ac and H2BK20ac regions were used in these analyses. Distance between candidate enhancer and TSS was determined by the distance between the commonly marked H3K27ac and H2BK20ac region and the nearest TSS. The bar charts show the relationship between cell-type-specificity (b) or A-485-dependent gene regulation (c), enhancer rank category, and enhancer-TSS distance.

### Strong enhancers frequently occur near their target genes

Next, we investigated the reasons for the relatively greater impact of enhancers on CTS than HK genes (**Fig. 1d**). H2BK20ac-based, but not H3K27ac-based, enhancer ranking shows that the strongest proximal enhancers occur more frequently in proximity to CTS genes than to HK genes (**Fig. 3b, Extended Data Fig. 9a**). For example, in ESC, EpiSC, and NPC, 17-32% of CTS genes harbored Top25% enhancers within 10kb, whereas only 6-9% of HK genes had Top25% enhancers within 10kb of their TSS. This shows that strong enhancers are positioned more frequently (odds ratio, ESC, 3.9, p <2.2e-16; EpiSC 3.4, p <2.2e-16; NSC 6.7, p <2.2e-16, two-sided fisher test) near CTS genes as compared to HK genes. Notably, this difference was negligible or very small (odds ratio, ESC, 1.2, p =0.056; EpiSC 1.0, p =0.8411; NSC 0.80, p =0.0063, two-sided fisher test) if enhancers occurred further away (>10kb) from TSS, where a similar fraction of enhancers occurred near HK, CTS, and CTE genes. Analysis of the A-485 downregulated gene class confirmed this trend and showed that a greater fraction of HD genes harbor strong proximal enhancers (Top10%, 25%, <10kb) than ND genes (**Fig. 3c, Extended Data Fig. 9b**). The results show that enhancers are positioned non-randomly, occur less frequently near ND genes, and those that occur in proximity to ND genes are weaker and located distally from TSS. Thus, the infrequent occurrence of strong proximal enhancers is one of the reasons for the modest regulation of HK genes.

### Enhancers boost transcription of ubiquitously expressed genes

To confirm and further understand the reason for the quantitative differences, we manually inspected tracks of A-485-regulated genes. We find that, within the same cell type, neighboring genes are quantitatively differentially affected (**Fig. 4a**). As an example, A-485 almost completely ablates *WNT16* and *MIR31HG* expression in HCT116, but the proximally-occurring genes *FAM3C* and *MTAP* are only partially downregulated. *FAM3C* and *MTAP* expression is reduced to a level that is similar to other cell lines in which the gene is not enhancer regulated. We refer to this type of partial regulation (*FAM3C*, *MTAP*) as enhancer-dependent transcriptional ‘boosting’ to discriminate it from the canonical strong enhancer-dependent regulation (*WNT16*, *MIR31HG*), where virtually all transcription is dependent on enhancer activity (**Fig. 4a**). Of note, H3K4me3 indicates two alternative TSS for *MIR31HG*, expression from one of the promoters is fully ablated whereas the other promoter retains weak expression, indicating differences in the autonomous activities of the two alternative promoters. Enhancer-dependent boosting is observed in all cell lines and frequently impacts genes that are ubiquitously expressed, some of them conserved down to single-celled eukaryotes, or generally important for human cell proliferation, such as *RAD21*, USP7, *MPHOSPH10*, *PSMA6*, and *RNF168* (**Extended Data Fig. 10**). Enhancer-dependent boosting of HK genes, which are conserved in non-metazoan organisms, shows that regulatory enhancer capacity can be ‘retrofitted’ onto genes that evolved prior to the advent of enhancers and are generally transcribed enhancer-independently.

**Fig. 4.**
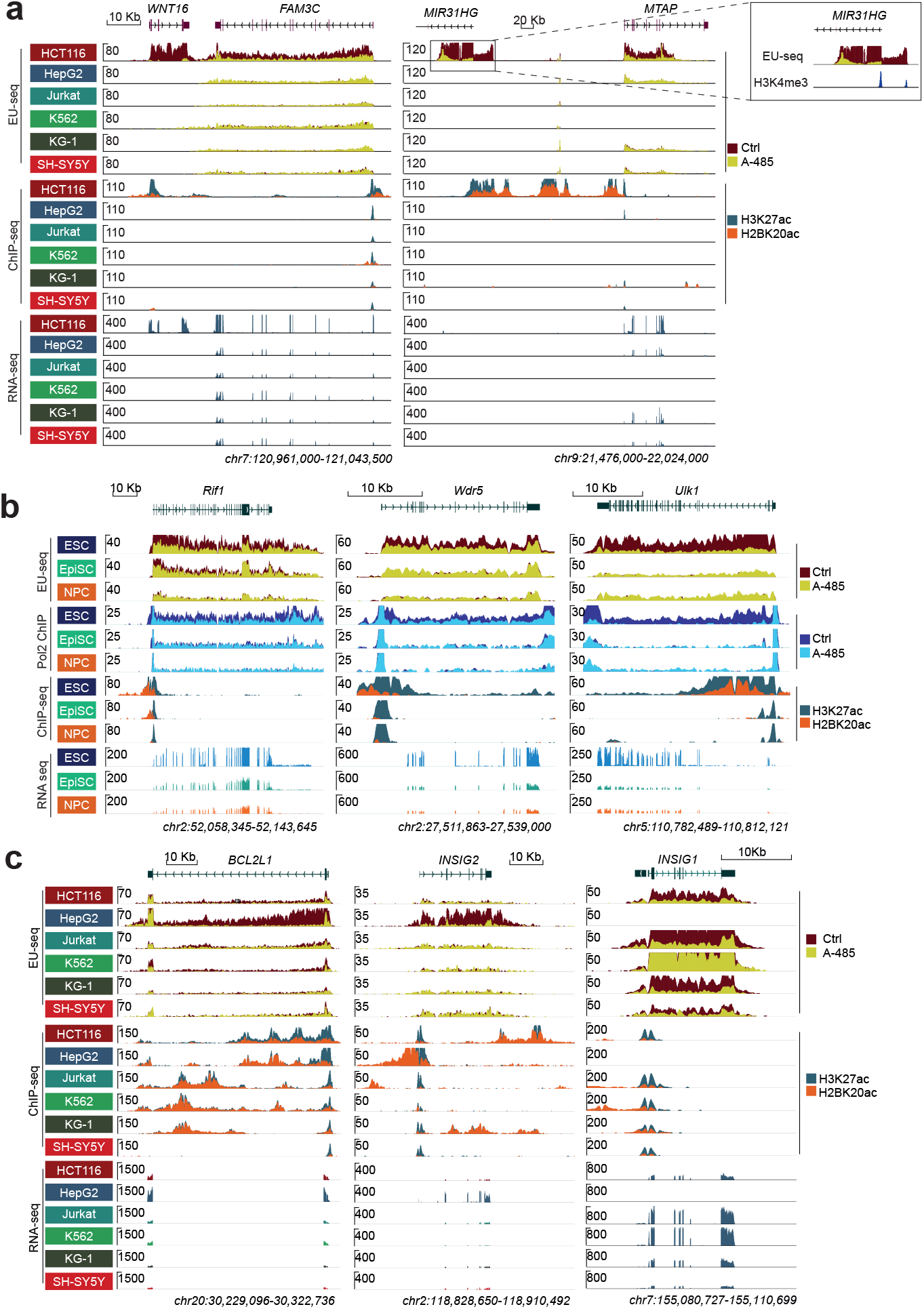
Enhancers cell type specifically increase the expression of ubiquitously expressed genes. **a,** Gene tracks showing quantitative differences in A-485-induced gene downregulation. *WNT16* and *MIR31HG* are cell type specifically expressed in HCT116, and the A-485 treatment completely abrogates their transcription. In contrast, the proximal genes, *FAM3C* and *MTAP*, respectively, are expressed in other cell lines, but their expression is elevated in HCT116, and A-485 treatment specifically reduces FAM3C expression in HCT116. Unlike *WNT16 and MIR31HG, FAM3C* and *MTAP* expression in HCT116 is not abrogated by A-485 but quantitatively reduced. After A-485 treatment, the expression of *FAM3C* and *MTAP* in HCT116 becomes similar to that of other cell lines in which the gene is not affected by A-485. **b,** Genome browser tracks of *Rif1, Wdr5,* and *Ulk1* showing their cell-type-specific differential expression and A-485-induced regulation in ESC, EpiSC, and NPC. Overlaid tracks show nascent transcription and Pol2 binding in untreated and A-485 treated (1h) cells. *Rif1* is strongly down-regulated in ESC, weakly downregulated in EpiSC, and remains unaffected in NPC. *Wdr5* and *Ulk1* are downregulated in ESC only. Changes in nascent transcription are mirrored in a cell-type-specific decrease in Pol2 binding. H3K27ac and H2BK20ac mark proximal candidate cis-regulatory elements, and RNA-seq shows cell-type-specific quantitative variation in the expression of the indicated genes. **c,** Genome browser view of *BCL2L1, INSIG1,* and *INSIG2* showing a loss of *INSIG1* expression, and a greatly elevated expression of *BCL2L1* and *INSIG2* in HepG2. *BCL2L1* is variably expressed, with HepG2 showing the strongest expression and the highest A-485-induced downregulation. *INSIG2* expression is greatly elevated in HepG2, and its expression is specifically downregulated in HepG2. H3K27ac and H2BK20ac show candidate cis-regulatory regions in each cell line.

### Enhancer-dependent boosting is functionally consequential

To consider the biological implications of enhancer ‘retrofitting’, we analyzed HK genes with known heightened cell-type-specific roles. *Wdr5*, *Rif1*, and *Ulk1* are evolutionarily conserved from budding yeast to humans and their elevated expression is linked to their heightened function in stem cells. In DepMap analyses^45^, *WDR5* is essential (1054/1054) in human cell lines, and its increased expression regulates pluripotency^46^. Autophagy, and the autophagy-regulating kinase ULK1, regulate pluripotency and early stages of embryonic development^47, 48^. RIF1 is a conserved regulator of replication timing, ESCs express a high level of RIF1, and its deletion causes the most dramatic changes in the replication timing in stem cells^49, 50^. *Rif1* knockdown impairs mESC proliferation and pluripotency^51^. Consistent with their heightened functional importance in stem cells, ESC expresses the highest levels of *Rif1*, *Wdr5*, and *Ulk1* (**Fig. 4b**). *Wdr5* and *Ulk1* expression is selectively enhancer-boosted in ESC. *Rif1* expression is strongly enhancer-boosted in ESC, modestly boosted in EpiSC, while the gene is expressed enhancer independently in NPC (**Fig. 4b**). Notably, in all these instances, expression of the genes is not ablated, but reduced, and equilibrates at a similar steady-state level in all 3 cell types after the A-485 treatment. This demonstrates that these genes are broadly expressed independently of enhancers and mounting of enhancers onto them increases the transcript dosage cell-type specifically.

To verify the functional relevance of enhancer-dependent *Rif1* boosting in ESC, we conditionally deleted the *Rif1* upstream enhancer (**Extended Data Fig. 11a**). *Rif1* expression was reduced ∼80% after the enhancer deletion (**Extended Data Fig. 11b**). For comparison, we also fused RIF1 with GFP-FKBP12^F36V^ to acutely deplete RIF1 protein using the dTAG approach ^52^ (**Extended Data Fig. 11c-d**). This allowed us to compare the phenotypic effect of enhancer-dependent *Rif1* boosting with RIF1 depletion. Within 3-5 days after inducing enhancer deletion or RIF1 depletion, ESC growth was reduced (**Extended Data Fig. 11e-f**), indicating the importance of enhancer-dependent *Rif1* boosting for ESC proliferation.

To investigate the potential relevance of enhancer-dependent boosting in the disease context, we analyzed genes that show preferential essentiality in HepG2, HCT116, Jurkat, K562, and SH-SY5Y in DepMap^45^ (gene essentiality data were not available for KG-1). Except for HCT116, other cell lines included between 2-5 enhancer-regulated genes among the Top10 cell line preferential essential genes, including cell-type-specific TFs (*MYB, RUNX1, HNF1A, ISL1*) and signaling proteins (*ALK, IRS1, IRS2*) (**Extended Data Fig. 12a**). Interestingly, ubiquitously expressed *INSIG2* and *BCL2L1* score among the Top10 preferentially essential genes in HepG2. *BCL2L1* expression is strongly boosted in HepG2 by proximal enhancers (**Fig. 4c**). Most notably, *INSIG2* scores as an essential gene in just 2 (out of 1054) cell lines, with HepG2 showing the highest dependency. INSIG2 and INSIG1 are crucial regulators of cholesterol homeostasis^53^. *INSIG1* and *INSIG2* are expressed in virtually all tissues (**Extended Data Fig. 12b**). Conspicuously, HepG2 lacks *INSIG1* expression and shows dramatic enhancer-dependent boosting of *INSIG2*, whereas in all other cell lines *INSIG2* is expressed enhancer-independently (**Fig. 4c**). This implies that enhancer-mediated *INSIG2* boosting provides a compensatory mechanism for the loss of *INSIG1* in HepG2, offering a plausible explanation for the unique essentiality of *INSIG2* in HepG2.

Further supporting the functional relevance of enhancer-dependent boosting, a recent study showed that an oncogenic enhancer increases the expression of *SEPHS2* in acute myeloid leukemia (AML) and creates unique sensitivity of AML cells to dietary selenium restriction^54^. Enhancer deletion only partially (<50%) reduces *SEPHS2* expression, but significantly impairs AML cell proliferation^54^. Our work independently confirms enhancer-dependent boosting (∼50%) of *SEPHS2* in AML-derived KG-1, but not in HCT116, SH-SY5Y, or Jurkat (**Extended Data Fig. 12c**). We extend the prior findings and show that *SEPHS2* expression is also enhancer-boosted in the non-AML cell line HepG2, but by a different enhancer than the one used in AML cells (**Extended Data Fig. 12c**). Previously, it remained unclear why deletion of the enhancer only partially reduced *SEPHS2* expression in AML cells, and why deletion of the same enhancer did not decrease *SEPHS2* expression in non-AML cells^54^. We find that *SEPHS2* can be expressed enhancer-independently, and its expression can be boosted cell-type-specifically, by distinctly positioned enhancers, offering plausible answers to the unresolved questions^54^ (**Supplementary note 2**).

### Enhancer and PRE strengths are tuned independently and cell-type-specifically

Next, we sought to quantitatively assess the relative impact of enhancers and PREs in gene activation. Nascent transcript abundance provides a reasonably direct measure of the promoter output, which in turn reflects the sum of enhancer and PRE contribution. A-485-induced decrease in nascent transcription should reflect enhancer contribution, and the remaining gene expression after A-485 treatment should reflect PRE-dependent autonomous promoter strength. In unperturbed cells, nascent transcript abundance of HD, ID, SD, and ND genes is quite similar and varies by >3 orders of magnitude, but after A-485 treatment, expression of SD and ID genes is sustained at a much higher level than HD genes (**Extended Data Fig. 13**). Interestingly, enhancers appear to similarly contribute to the expression of genes with varying promoter strength, but the relative impact of enhancers appear stronger in genes that have weaker promoters (**Extended Data Fig. 13b**). This shows that HD promoters are mainly activated by enhancers, ND genes are activated by PREs, and SD and ID genes are partially activated by PREs, and their expression is boosted by enhancers.

Enhancer strength is tuned greatly across cell types. For example, the TFs *Sox2*, *Spry2*, *Spry4*, *ID1*, and *SOX4* are regulated by enhancers in all cell lines expressing the genes, yet their nascent transcript abundance varies by up to ∼30 fold in different cell types (**Extended Data Fig. 14a-b**). Enhancer strength is also tuned for non-TF proteins, such as *LDLR*, *SLC43A3*, *HMGCS1*, and *CDK6* (**Fig. 5a, Extended Data Fig. 14c**). Thus, for the same genes, enhancer strength can be tuned by orders of magnitude. Similar to enhancers, PRE strength is also adjustable cell-type-specifically (**Fig. 5b, Extended Data Fig. 15**). Alternative promoters of the same genes, such as *Cbx7*, *Rbpj*, *ARHGEF18*, and *EIF2AK4*, are differentially activated by enhancers or PREs (**Extended Data Fig. 16**). Yet more remarkably, expression of cell-type-specific genes is adjusted by tuning either enhancer or PRE strength (**Fig. 5c, Extended Data Fig. 17**). The combined function of enhancers and PREs also impacts broadly expressed genes, and can even affect some of the most highly expressed genes, such as *MALAT1*, although in such instances the measurable enhancer impact appears relatively modest in cell lines that have strong PREs (**Extended Data Fig. 18**).

**Fig. 5.**
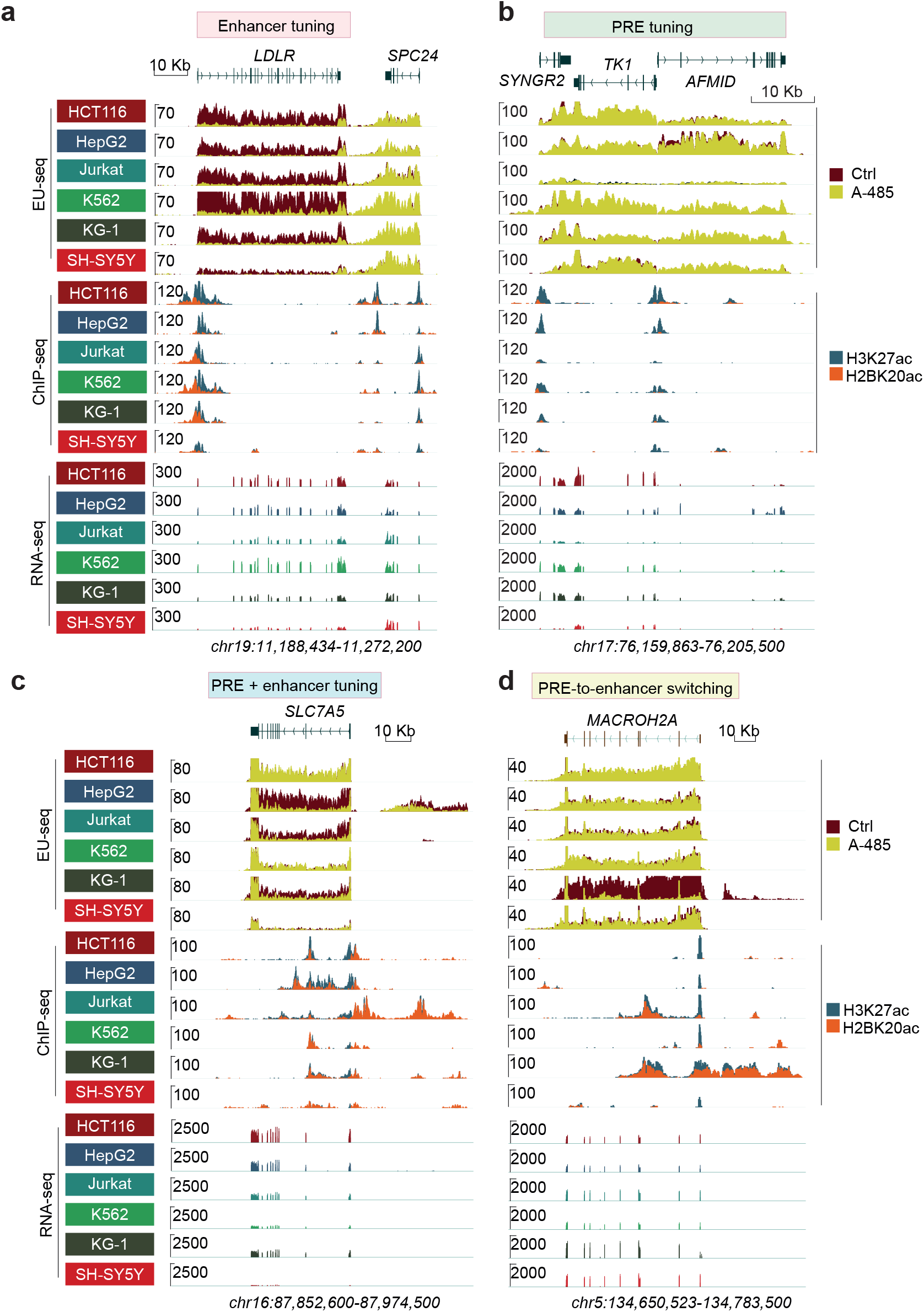
Tunable and integrated strengths of PREs and enhancers control cell-type-specific gene expression. **a,** Gene tracks showing cell-type-specific variability in enhancer-dependent LDLR expression. LDLR is downregulated by A-485 in all cell lines but its expression is highly variable across cell lines, showing gene expression tuning by cell-type-specific tuning of enhancer strength. Proximally located SPC24 is activated by PREs and remains unaffected by A-485. **b,** Genome browser tracks showing enhancer-independent, cell-type-specific quantitative variation in gene expression, likely through changing PRE strength**. c,** Genome browser tracks showing cell-type-specific quantitative variation in *SLC7A5* expression, either through changing PRE strength (SH-SY5Y, K562, HCT116) or through the combination of changing PRE and enhancer strength (KG-1, Jurkat, HepG2). **d,** Genome browser tracks showing promoter-enhancer switching of histone *MACROH2A* regulation. The expression of *MACROH2A* in KG-1 is abrogated by A-485, but its expression remains unaffected in all the other cell lines.

Strikingly, for the same genes, regulation can be switched cell-type-specifically. Histone *MACROH2A* is expressed enhancer-independently in 5 cell lines, including Jurkat, which has nearby enhancers (**Fig. 5d**). Yet, *MACROH2A* transcription is completely switched to being enhancer-dependent in KG-1. Although this type of switching is infrequent, we found several examples (*ISOC1*, *FECH*, *MED14*, *CSK, GSTO1*) showing this type of dramatic change in regulation (**Extended Data Fig. 19**). Cells showing PRE-to-enhancer switching harbor H2BK20ac marked strong proximal candidate enhancers. The concordance between the acquisition of cell-type-specific candidate enhancers and change in gene regulation suggests that PRE-to-enhancer switching occurs as a consequence of cell-type-specific enhancer mounting. Of note, the overall expression of some of the genes, such as *MED14* and *FECH*, is similar in all cell lines, yet the type of regulation differs, suggesting that PRE strength is reduced in cells that have acquired strong proximal enhancers. Collectively, these results show (1) that enhancer and PRE strength are tuned independently of each other, (2) enhancer and PRE strengths are adjusted gene-specifically and cell-type-specifically, and (3) the same genes can be activated jointly or interchangeably by PREs and enhancers. From this, we conclude that, in the same cell type, a single CP can accommodate input signals from enhancers and PREs.

### DNA sequences discriminate between autonomous and enhancer-dependent promoters

Ultimately, gene regulatory information must be encoded in the DNA sequences. We sought to identify sequence features that discriminate between enhancer-dependent and –independent promoters (i.e. ND versus HD promoters). First, we compared the frequency of two main mammalian core promoter motifs: TATA box and Initiator (Inr). TATA-box occurred more frequently among HD than ND genes (**Extended Data Fig. 20a-b**), but, even among HD genes, 84-91% lacked a TATA-box. CpG island (CGI) density is higher in promoters of HK genes than CTS genes^55, 56^. We found that CGI frequency gradually decreases (ND>SD>ID>HD1>HD2) with A-485-induced gene downregulation (**Fig. 6a; Extended Data Fig. 20c**). Normalized CpG density (observed to expected, O/E, ratio) is inversely associated with enhancer-dependent gene regulation. The O/E ratio was highest in ND promoters and gradually decreased with the extent of A-485-induced gene downregulation (**Fig. 6b; Extended Data Fig. 20c**).

**Fig. 6.**
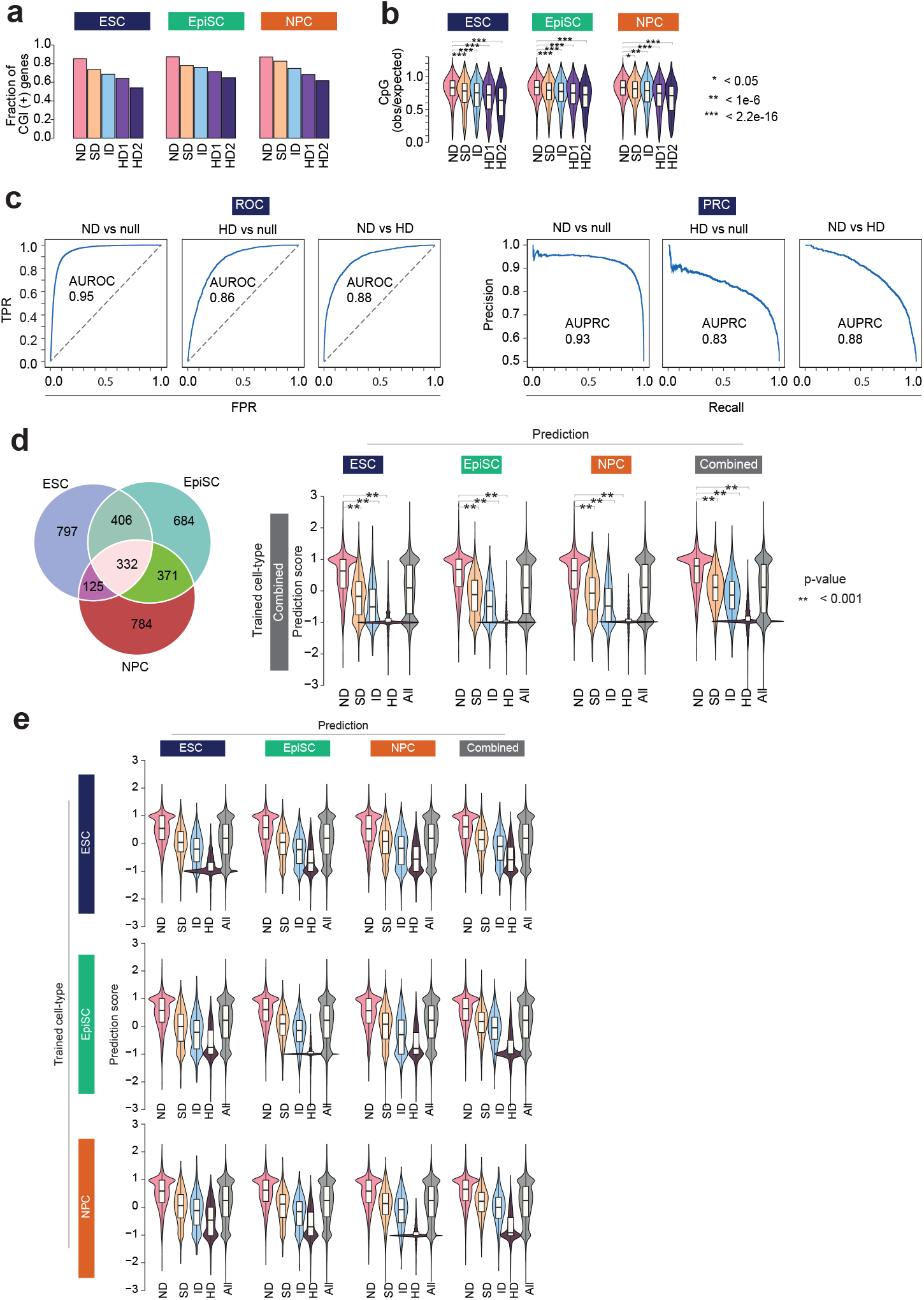
Sequence differences separate HK and CTS promoters. **a-b,** Promoter CpG island (CGI) density inversely relates to enhancer-dependent gene regulation. The fraction of CGI positive (CGI +) genes (a) and normalized (observed to expected, O/E) CpG density (b) among the indicated A-485 regulated gene categories. A-485-regulated genes were classified as specified in Fig. 1b, and HD genes were further subdivided into two groups: HD1 (downregulated ≥2-4-fold) and HD2 (downregulated ≥4-fold). ** p < 0.001, Mann–Whitney U test. c, Promoter sequences can discriminate ND and HD genes. The receiver operating characteristic (ROC, left panel) and precision-recall cure (PRC, right panel) show that promoters of ND and HD genes can be separated from each other, and random genomic sequences (null sequence). The classifiers were trained using DNA sequences, 500bp upstream from TSS of ND and HD genes from ESC, EpiSC, and NPC, or randomly selected genomic sequences of similar length and GC content (see Methods for details). The area under the ROC curve (AUROC) and area under the PRC (AUPRC) are shown. **d-e,** Promoter sequence differences predict enhancer-dependence of genes. The classifier was trained using ND and HD genes as positive and negative sets respectively. The Venn diagram shows number of HD genes in the three cell lines. A gkm SVM model was trained using HD genes combined from the three cell lines and performance was tested in each of the cell lines. **e,** A gkm SVM model was trained using ND (negative set) and HD (positive set) genes from one of the indicated cell lines. Performance of the model was tested on the trained cell line as well as in the other two held-out cell lines.

To test if promoter sequences can discriminate ND and HD promoters, we trained gapped k-mer support vector machine (SVM)-based kernels (gkm-SVM) (see Methods). The classifier was able to distinguish ND promoters from null sequences (i.e. randomly selected sequences of matching GC and repeat content) with very high accuracy (mouse: AUROC 0.95, AUPRC 0.93; human: AUROC 0.96, AUPRC 0.95) (**Fig. 6c; Extended Data Fig. 21a**). The classifier also separated HD promoters from null sequences, but with a lower accuracy (mouse: AUROC 0.86, AUPRC 0.83; human: AUROC 0.84, AUPRC 0.81), indicating that ND promoters are more information-rich than HD promoters. Notably, ND and HD promoters could be discriminated with good accuracy (mouse: AUROC 0.88, AUPRC 0.88; human: AUROC 0.85, AUPRC 0.83).

Next, we tested the performance of the sequence-based model in separating A-485 regulated gene classes. We used A-485-induced gene regulation information from the isogenic mouse cell lines and trained classifiers using the following three gene sets. (1) positive set: all HD genes combined from the three cell lines; negative set: ND genes in all the three cell lines. (2) positive set: HD genes from one of the cell lines; negative set: ND genes in all three cell lines. (3) positive set: HD genes exclusively downregulated in one of the cell types; negative set: ND genes in all three cell lines.

The model trained on the combined gene set performed best and separated HD and ND genes in all cell lines (**Fig. 6d**). Notably, the classifier also discriminated held-out SD and ID genes, which were not included in the training dataset. A model trained using ND and HD genes from individual cell lines also performed well, and separated genes in the trained cell type as well as in the held-out cell types (**Fig. 6e**). The model trained on genes that were exclusively HD in one of the cell types showed modest performance, both in the cell type in which it was trained and in the held-out cell types (**Extended Data Fig. 22**). A model trained in human cell lines also performed well (**Extended Data Fig. 23a-b**). The model trained in mouse cell lines could also separate A-485 regulated gene class in human cell lines, albeit with decreased performance (**Extended Data Fig. 23c**). These results indicate that promoter-proximal sequences include information that can reasonably predict their autonomous activation. This indicates that, to some extent, sequence features that segregate ND and HD promoters are evolutionarily conserved.

### Validating greater intrinsic autonomous activity of ND than HD promoters

To identify sequence motifs that segregate ND and HD promoters, we derived *de novo* motifs from importance scores using the trained gkm-SVM kernel. Sequences enriched in ND promoters are CG-rich and match motifs of TFs, such as NRF1, GABPA, YY1, ETS1, ZNF143, and THAP11, which are broadly expressed and known to bind promoters of HK genes ^57–61^ (**Extended Data Fig. 21b**). To confirm this prediction, we analyzed the binding of NRF1, GABPA, ZNF143, and YY1, as well as the chromatin remodeler INO80 that binds to YY1, in ESC. As a reference, we also included CTCF, which is directly implicated in chromatin loop formation and enhancer-promoter communication^28, 62^. Except for CTCF, a larger fraction of ND promoters were occupied by these factors than HD promoters, and YY1 and INO80 co-occupied a majority of ND promoters (**Fig. 7a**). Notably, at the occupied promoters, there was no difference in CTCF binding in ND and HD promoters, but GABPA, YY1, INO80, and NRF1 binding was significantly different and progressively decreased (ND>ID>SD>HD) with increasing A-485-induced gene downregulation (**Fig. 7b**). These results validate the model’s prediction and suggest that sequence differences in promoter-proximal regions possibly dictate differential recruitment of housekeeping TFs to ND and HD promoters.

**Fig. 7.**
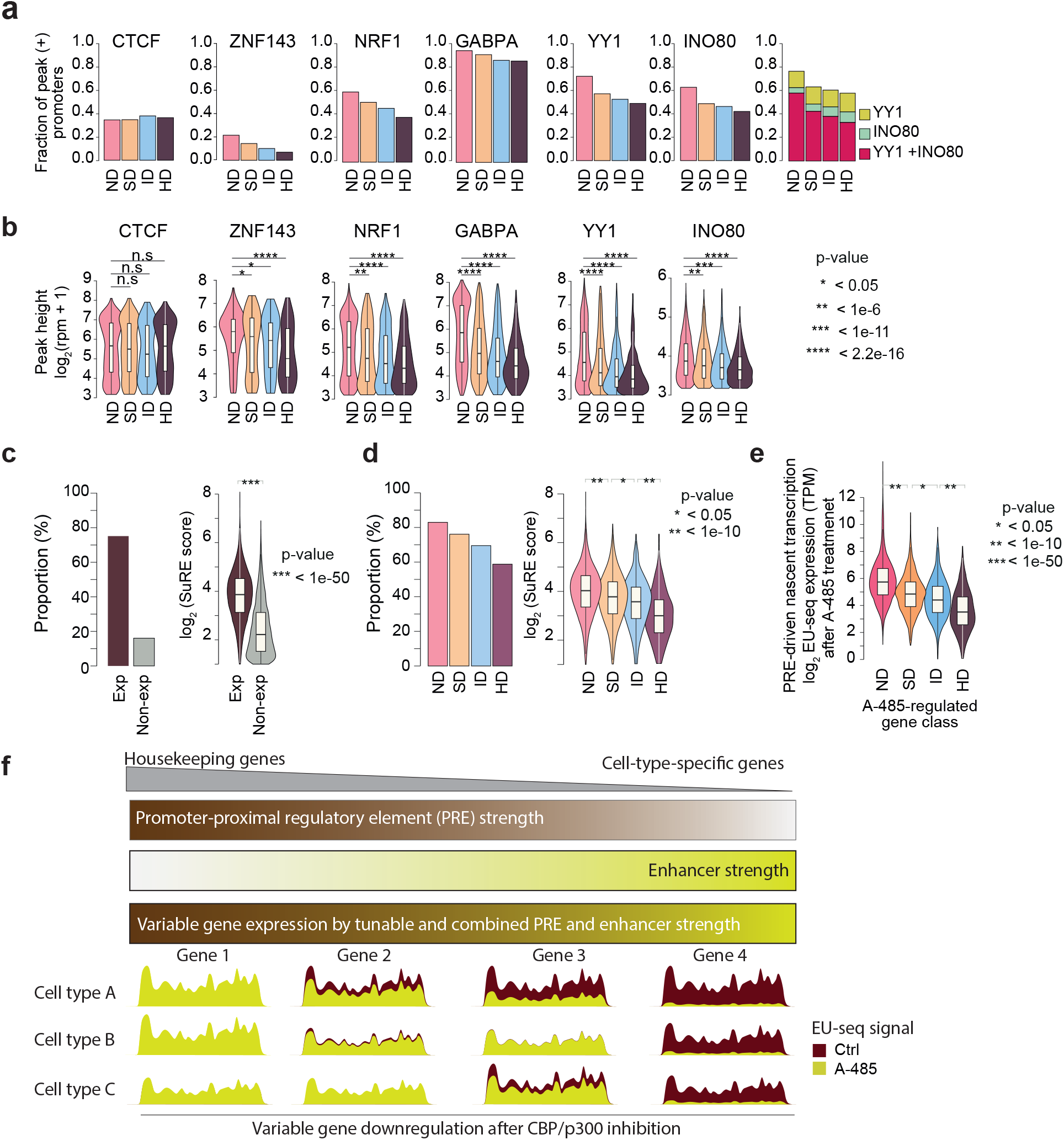
Distinct TF binding and autonomous activities of enhancer-dependent and -independent promoters. **a,** Shown is the promoter binding of the indicated transcription factors (TFs) and INO80 at the indicated A-485 regulated gene categories in mouse ESC. Bar charts show the fraction of promoters bound by the indicated proteins. Violin plots, below each bar chart, show the ChIP intensity of the indicated proteins at promoters of the respective gene class. ChIP-seq data were re-processed from references listed in the Methods section. **b,** Portion of K562 promoters whose activity was quantified in SuRE analyses^63^ (left panel) and SuRE scores for the indicated groups of promoters (right panel). Exp: promoters of genes expressed in K562; Non-exp: promoters of genes not actively transcribed in K562. **c,** Portion of the indicated A-485-regulated gene class promoters scored in the SuRE dataset (left panel), SuRE scores in the indicated groups of promoters (right panel). **d,** Shown are the nascent transcript expression levels (TPM) in the indicated A-485-regulated gene class. Nascent transcription is quantified using EU-seq, in K562 cells treated with or without A-485 (1h). This analysis only included promoters that scored in SuRE analyses. **e, f,** Proposed model for quantitative gene expression variation in mammalian cells through tunable and integrated strengths of PREs and enhancers.

The differences in TF binding are sportive, but not demonstrative, of the differences in ND and HD promoter activity. To directly assess the autonomous activity of ND and HD promoters, we used the SuRE (survey of regulatory elements) dataset, which possibly represents the most comprehensive measurement of autonomous promoter activity in mammalian cells^63^. SuRE covered 75% of promoters that are active in K562 and promoters of actively transcribed genes had higher SuRE scores than promoters of non-expressed genes (**Fig. 7c**). The fraction of SuRE^+^ promoters decreased with increasing A-485-induced gene downregulation; 83% of ND promoters were SuRE^+^ whereas only 59% of HD promoters were SuRE^+^ (**Fig. 7d**). Notably, in SuRE^+^ promoters, SuRE scores progressively decreased with the extent of A-485-induced gene downregulation. ND promoters had the highest SuRE scores, and HD promoters had the lowest SuRE scores. Mirroring significant differences in SuRE scores, the level of PRE-driven endogenous transcription (the remaining nascent transcription after A-485 treatment) at A-485-regulated gene class is significantly different (**Fig. 7e**).

The concordance between SuRE scores and A-485-induced gene downregulation is remarkable considering that SuRE assesses promoter activity in an ectopic reporter, whereas in EU-seq, we indirectly infer the autonomous activity of native promoters by eliminating enhancer function. Together, these findings support the model that enhancer-dependent and -independent gene expression is segregated through differences in PREs, which are predominantly positioned in proximity to core promoters. The autonomous promoter activity progressively decreases from ND>ID>SD>HD.

## Discussion

This work provides a first genome-wide view of the endogenous E-P specificity in mammalian cells, demonstrating that enhancers are broadly compatible and predominantly activate the nearest accessible promoters. Our findings revise the idea that the nearest promoter is not the true target of human enhancers^4, 32, 33^.

Reporter-based systems are powerful in revealing E-P specificity in isolation but do not directly inform how context-dependent native transcription landscapes are adjusted cell-type-specifically. In our approach, enhancer specificity is inferred indirectly, but a salient advantage is that it directly informs the functional impact of enhancers on endogenous genes. Thus, our approach complements the existing methods and offers much-needed in vivo validation for reporter-based observations. Importantly, our work goes beyond revealing native E-P compatibility; we use the compatibility rules to illuminate quantitative gene expression variation within and across cell types. The work showcases the usefulness of improved enhancer markers and comparative analyses for uncovering and explaining the complexities of native gene expression. For example, had we not used H2BK20ac as an enhancer marker, the striking relationships between enhancer distance, strength, and gene regulation could not have been uncovered. Similarly, without studying multiple cell types, it would have been impossible to uncover the regulatory intricacies and explain how E-P compatibility shapes gene expression variability across cell types.

Below, we discuss the major findings of our work and their conceptual implications in understanding gene regulation and the evolution of new regulatory networks.

### Enhancers prominently act locally

High-resolution Micro-C-based chromatin contact maps suggest a median enhancer-promoter distance of ∼100kb ^64, 65^, and lower resolution enhancer-promoter interaction analyses imply that as many as half of enhancers skip their proximal genes to activate distally located genes^4–10^. If enhancers were to frequently skip their nearest active promoter, we would expect many enhancer-proximal genes to remain unaffected by CBP/p300 inhibition, and many non-enhancer proximal genes to be downregulated. In all cell lines investigated, most of the genes occurring in proximity to strong enhancers are downregulated after CBP/p300 inhibition, whereas genes that are located distally are seldom affected. If a large fraction of enhancers were incompatible with nearest promoters and frequently skipped their nearest active promoters, we would not expect such a clear relationship between enhancer-promoter distance and gene regulation.

Our finding, derived from tens of thousands of endogenous enhancers and promoters, suggests that enhancers act prominently on proximal open promoters. The local enhancer action may help explain why global ablation of TADs and large architectural loops have a limited impact on gene regulation ^66, 67^. We find that the activation potential of enhancers decreases sharply with the distance from the promoter, and weakly impacts genes beyond >50kb distance, providing genome-scale validation for reporter-based analyses in a single TAD^68^. We do note robust evidence showing that enhancers can and do act at long genomic distances ^3^. We acknowledge an important role of chromatin looping, but this mechanism may be more broadly used for reducing large enhancer-promoter distances rather than for skipping nearest genes and activating distal genes. Gene skipping may occur, but much less frequently than implied currently.

### Enhancers and PREs jointly shape transcription landscapes

Our work reveals that mammalian gene regulation is remarkable plastic and that gene regulation is not strictly segregated into ‘enhancer-dependent’ and ‘enhancer-independent’ types. We propose that PRE and enhancer function in gene regulation are not mutually exclusive; instead, cell-type-specific transcription landscapes are quantitatively shaped through tunable and integrated strengths of PREs and enhancers, providing crucial in vivo validation of reporter-based observations^12, 13^. Notably, enhancer and PRE strength are tuned independently of each other, across diverse cell types, providing a highly modular mechanism for dynamic gene regulation. We find hundreds of genes (SD and ID gene class) that are not strictly enhancer-dependent, but whose expression is boosted by enhancers, showing that the combined gene activation by enhancers and PREs is not a rarity, but a widespread feature of mammalian genes. This finding demonstrates that, in the same cell type, in its native context, a single CP can accommodate input signals from PREs and enhancers. We suggest that enhancers and PREs can act as independent regulatory units and broad compatibility is likely a general property of mammalian CPs.

### Broad enhancer compatibility expands the regulatory complexity

Qualitatively, many human genes are ubiquitously expressed, but their expression is quantitatively altered across tissues^69^. If enhancer compatibility was constrained, as implied in the enhancer specificity model, it would render a large portion of human genes beyond the scope of enhancers. Broad compatibility expands enhancer function beyond CTS genes and affords a widespread role to enhancers in quantitative boosting of HK genes across tissues in development and disease. In the context of evolution, we suggest that broad E-P compatibility provided an elegant mechanism for ‘retrofitting’ new regulatory complexity onto HK genes, whose origin pre-dates the advent of enhancers. At the same time, keeping the CP architecture preserved allowed the cell to use the same conserved basal machinery for transcribing all genes. We suggest that broad enhancer compatibility is not an accidental invention but a key design principle with important functional relevance.

It is important to emphasize that broad E-P compatibility implies that enhancers are capable of activating most promoters, but it does not mean that all promoters are quantitatively activated to the same extent. We find that CTS genes are most prominently regulated by enhancers, CTE genes are affected to an intermediate level, and HK genes are least affected. At least two different factors contribute to this. (1) Strong enhancers are more often positioned near CTS genes than CTE and HK genes. HK genes have weaker enhancers that are often located more distally. (2) Native genes are activated by the integrated activity of enhancer and PRE. The relative impact of an enhancer on transcription depends on the autonomous strength of the promoter, and the relative impact of the enhancer is related to promoter strength. Because the relative strength of PREs can change in different cell types, relative enhancer responsiveness can be variable across cell types.

### PREs can sustain partial gene activation without enhancers

The combined function of PREs and enhancers has implications for understanding transcript dosage regulation. It has been noted that deletion of well-defined enhancers often causes only modest or no downregulation of target genes^70^. Currently, this phenomenon is often rationalized by transcriptional buffering by redundantly functioning ‘shadow enhancers’^71, 72^. Our findings show that hundreds of enhancer target genes are only partially downregulated even when enhancer function is ablated globally. This suggests that along with shadow enhancers, PREs have a previously unappreciated, widespread role in sustaining partial gene activation in the absence of enhancer activity. Thus, just like shadow enhancers, dual regulation by enhancers and PREs may provide robustness in gene regulation by buffering against deleterious mutations in one of the regulatory elements.

### Broad compatibility lowers the threshold for evolving new regulatory networks

Non-coding genetic variants can create de-novo enhancers that can act as new regulatory modules^73–75^. Indeed, the evolution of organismal complexity primarily involves mutations in the cis-regulatory sequences that quantitatively alter the expression of functionally conserved proteins^76^. Restricted enhancer compatibility imposes greater constraints on the emergence of new gene regulatory networks because it requires a simultaneous change in incompatible CP sequences. Broad E-P compatibility lowers the threshold for implementing new regulatory capacities and facilitates the emergence and re-wiring of regulatory networks in disease and organismal evolution.

### Sequence differences in PREs segregate autonomous and non-autonomous promoters

We propose that enhancer-dependent and -independent genes are mainly segregated by the sequence-encoded differences in their PREs. Enhancer-independent genes, which prominently include HK genes, harbor strong PREs that potently recruit ubiquitously expressed TFs and act as ‘built-in’ activation modules. Enhancer-dependent genes, which mostly include CTS genes, contain weaker PREs. Consequently, CTS promoters have weaker autonomous activation potential. This brings two advantages (1) weaker autonomous promoter activity avoids ubiquitous expression of CTS genes, and (2) weaker autonomous promoter activity generates a dependence on enhancers for their activation. CGI density appears to be one of the features that afford greater autonomy to PREs. This can rationalize higher CGI density in HK promoters than in CTS promoters^57, 61, 77, 78^, stronger binding of ubiquitously expressed TFs to CGI-rich promoters^58–60, 63^, the stronger activity of CGI promoters in reporter assays^63^, and increased promoter activity with increasing CG dinucleotide content^79^. Of note, for simplification, we refer to promoters as autonomous and non-autonomous, but this binary classification is inaccurate. Promoters have a graded extent of autonomous activation potential. Importantly, the autonomous activity of promoters is not fixed; for the same promoters, activation strength varies across cell lines. Thus, the autonomous activity of promoters neither means that promoter strength is untunable nor does it imply insensitivity of such promoters to distal enhancers.

Together, the presented findings provide major conceptual insights into endogenous enhancer specificity and cell-type-specific gene regulation and will facilitate efforts toward accurate modeling of enhancer-dependent transcription and functional interpretation of genetic variants. A remaining goal for future studies is to clarify whether there are other classes of distal enhancers that act independently of CBP/p300 and if their regulatory principles are any different from CBP/p300-dependent enhancers investigated in this work.

#### Supplementary note 1

Recently it was shown that different groups of human enhancers require different COFs ^21^. To evaluate the requirement of CBP/p300 enhancer activity in STARR-seq, we re-analyzed the published data. Specifically, we analyzed the requirement of CBP/p300 for enhancers that are accessible in the native context. The reason we focused on chromatin accessibility is that chromatin accessibility is a hallmark of endogenously active enhancers, and it is one of the most fundamental differences between native enhancers and enhancers cloned in plasmids. Surprisingly, among all COFs analyzed, only CBP/p300 showed a clear relationship with endogenous chromatin accessibility and enhancer reporter activity (**Extended Data Fig. 1**). Enhancers showing the greatest CBP/p300 dependence had the highest chromatin accessibility, enhancers with intermediate CBP/p300 dependence had an intermediate level of accessibility, and enhancers that were unaffected by CBP/p300 depletion lacked chromatin accessibility. This shows that, even in the ectopic plasmid-based reporters, CBP/p300 is required for the activity of enhancers that are open in the native chromatin context, supporting our assumption that CBP/p300 is required for the activation of natively accessible enhancers. The lack of association between STARR-seq activity and chromatin accessibility is consistent with the findings that STARR-seq can identify open and closed enhancers^13, 80^. Pioneering factors can bind to closed chromatin, but we are not aware of concrete evidence showing a major contribution of closed enhancers to native gene regulation, without acquiring accessibility under the conditions in which they are active.

#### Supplementary note 2

Deletion of the oncogenic *SEPHS2* enhancer did not impair expression in normal hematopoietic cells or cells that originated from other tissues^54^. Our work independently confirms this finding and shows that the enhancer boosts (∼50%) *SEPHS2* expression in the AML-derived KG-1, but not in HCT116, SH-SY5Y, or Jurkat (**Extended Data Fig. 12c**). Extending the prior findings, we find that *SEPHS2* expression is also enhancer-boosted in the non-AML cell line HepG2, but the position of enhancers is completely different in KG-1 and HepG2. Our results offer a plausible explanation for some of the questions that remained unresolved in previous work. For example, why is *SEPHS2* expression only partially reduced in enhancer deleted cells, and is the remaining *SEPHS2* expression sustained by shadow enhancers or other mechanisms? In our analyses, *SEPHS2* expression in KG-1 is only reduced by ∼50%, even when CBP/p300 activity is ablated globally, suggesting that remaining *SEPHS2* expression is likely sustained by PREs, not by yet undefined distal shadow enhancers. Why did deletion of the enhancer not impact *SEPHS2* in non-AML cells? Our findings suggest two different reasons for this. (1) Enhancers positions in KG-1 and HepG2 are completely different, showing that *SEPHS2* expression can be regulated by different enhancers in different cell types, which can explain why deletion of an enhancer sequence may affect expression in one cell type but not in another, even if the gene is regulated by enhancers in both the cell types. (2) *SEPHS2* is expressed enhancer-independently in HCT116, SH-SY5Y, and Jurkat. This demonstrates that the same gene can be activated by an enhancer in one cell type but not in others, and thus, the effect of enhancer deletion will be cell-type-specific. This exemplifies how studying gene regulation in multiple cell types can provide an improved mechanistic understanding of quantitative gene regulation. Had we used only one cell type, or simply analyzed enhancer activity with MPRA, it would be almost impossible to obtain these insights. Together, the above results support the idea that enhancer-dependent boosting of HK genes is an important feature of mammalian gene regulation, and it provides a mechanism for quantitatively adjusting transcript dosage in physiology and disease.

## Acknowledgments

We thank the members of the Choudhary lab for their helpful discussions. The Novo Nordisk Foundation Center for Protein Research is financially supported by the Novo Nordisk Foundation (NNF14CC0001). C.C. was supported by the Hallas Møller Investigator Award (NNF14OC0008541) from Novo Nordisk Foundation. This work was funded by the European Commission FP7 grant (SyBoSS FP7-242129). Y.H. was supported by a Grant-in-Aid for JSPS Overseas Postdoctoral Fellows. S.K. was supported by a Lundbeck Foundation Fellowship (R347-2020-2170). We thank the SyBoSS partners Konstantinos Anastassiadis, A. Francis Stewart, Austin Smith, and Bill Skarnes for sharing E14TG2a Oct4-IRES-Puro mouse ESCs. We thank Jonas Walter for generating Cre-ERT2 expressing mESCs. We thank Dr. Christopher J. Ott for providing dCBP1. We thank the CPR Imaging Platform, the CPR Big Data Management Platform, and the CPR and DanStem Genomics Platform for their assistance. We thank the ENCODE Consortium and the ENCODE production laboratory for generating and sharing the data.

## Author contributions

T.N. and C.C. conceived the project. K.N. differentiated ESCs and generated the isogenic EpiSC and NPC cell lines. T.N., Y.H., S.K., E.M. performed experiments and analyzed data. T.N. performed bioinformatic analyses. T.N. and C.C. wrote the manuscript with input from all co-authors.

## Material and Methods

### Cell lines and culture conditions

E14TG2a Oct4-IRES-Puro mouse ESC (mESC)^81^ were used for all genomics experiments. Where indicated, E14TG2a mESC (Sigma-Aldrich, Cat# 08021401) were used for manipulation (insertion or deletion) of enhancers. mESC were cultured in a custom-made (C.C.Pro GmbH, Oberdorla) N2B27 medium consisting of a 1:1 mix of DMEM/F12 and Neurobasal medium, but lacking Arginine and Lysine. Before use, the media was supplemented with 640 µM L-Arginine, 550 µM L-Lysine, 100 µM 2-mercaptoethanol, 150 µM sodium pyruvate, 0.5x B27 supplement (Thermo Fisher Scientific), 0.5x N2 supplement (Thermo Fisher Scientific), 1% penicillin/streptomycin, and cell type-specific growth factors or inhibitors. For mESCs the medium was supplemented with 1000 units/ml leukemia inhibitory factor (LIF) (Merck Millipore), 1 µM PD0325901, and 3 µM CT-99021 (custom-made by ABCR GmbH, Karlsruhe).

EpiSCs were derived from E14Tg2A Oct4-IRES-Puro ESCs by culturing them on dishes coated with 10ng/ml fibronectin (Merck Millipore), in N2B27 supplemented with 20ng/ml Activin A and 12ng/ml Fibroblast growth factor 2 (FGF2) (MPI-CBG protein facility, or Peprotech) for 14-15 passages. NPCs were generated from EpiSCs by seeding 0.5-1x10^5^ cells per cm^2^ to fibronectin-coated dishes in N2B27 without growth factors^82, 83^. After 5 days, the developed neural rosettes were dissociated with Accutase (PAA) and seeded to 0.1% gelatinized tissue culture plates in N2B27 supplemented with 10ng/ml FGF2 and 10 ng/ml Epithelial growth factor (EGF) (PeproTech). All cell lines were passaged regularly (every 2 to 4 days) with Accutase (PAA) or TrypLE (Thermo Fisher Scientific), and ESCs and EpiSCs were occasionally selected for Oct4 expression with 1µg/ml puromycin (Sigma). HepG2 and HCT116 were cultured in DMEM media supplemented with 10% FBS and 1% penicillin/streptomycin. Jurkat, KG-1, K562, and SH-SY5Y cells were cultured in RPMI-1640 media supplemented with 10% FBS, and 1% penicillin/streptomycin. All cells were cultured at 5% CO_2_, 37 °C. Where indicated, the cells were treated with the following inhibitors: A-485 (10µM), or dCBP-1 (100nM).

### RNA-seq

Following cell lysis with RLT lysis buffer (Qiagen), total RNA was extracted using RNeasy Mini Kit (Qiagen). 1ug of total RNA was used for enriching polyadenylated mRNA, the samples were processed using TruSeq Stranded mRNA Library Prep Kit (Illumina) as per manufacturer’s instructions and sequenced using a Next-Seq 500 Sequencer (Illumina).

### Nascent transcription analyses by EU-seq

Nascent transcription analyses were performed as described previously^34^. ESC EU-seq data was previously published (GSE146328) and re-processed for the downstream analysis. EpiSC, and NPC were treated with DMSO or A-485 (10µM) for 30, 60, and 120 min. dCBP1-induced nascent transcription changes were quantified after treating ESC with DMSO or dCBP1 (100nM) for 120 min. In all human cell lines, nascent transcription changes were quantified in cells treated with DMSO or A-485 (10µM) for 60 min. Cells were pulse-labeled with 0.5 mM 5-EU for 20 min before sample collection. Following 5-EU labeling, cells were quickly washed (1x) with PBS, and immediately lysed in RLT buffer, and RNA was prepared using RNeasy Mini Kit (Qiagen). The labeled RNA was subsequently enriched using the Click-iT® Nascent RNA Capture Kit (Thermo Fisher Scientific) according to the manufacturer’s instructions. RNA-seq libraries were prepared using the NEBNext Ultra kit (NEB) as per the manufacturer’s instructions and subsequently sequenced using a Next-Seq 500 Sequencer (Illumina).

### Reference genome annotation

Mouse and Human genome annotation and reference genome were downloaded from the GENCODE (mouse; GRCm38 release 25, human; Grch37 release 29)^37^. As a reference, the transcript type of “protein coding” and “lincRNA” was chosen as representative genes. For building cell-type-specific gene models, the expressed transcripts were selected using the criteria of Transcripts per million transcripts (TPM) > 2 in RNA-seq and the presence of H3K4me3 ChIP seq peak within 1Kb from TSS. The longest isoform was selected if a single H3K4me3 peak is proximal to multiple transcripts.

### RNA-seq

Adaptors and low-quality sequences (-q 20) were trimmed using Cutadapt (doi:https://doi.org/10.14806/ej.17.1.200). Reads were mapped to the reference genome using STAR (version 2.6.1a)^84^ with removing the non-canonical junctions. Reads mapped to tRNA and rRNA regions were removed using Bedtools (version 2.23)^85^. The number of reads mapped to exon was counted on a gene-basis using HTseq (version 0.11.1)^86^ TPM was calculated using R.

### EU-seq

Adaptor and low-quality sequence trimming was performed as described in the RNA-seq section. Read sequences were aligned to mm10 (mouse) using bwa aln with default parameters (BWA version 0.7.10^87^. Multi-mapped reads and reads with more than three mismatches were removed using samtools (version 1.4)^88^. Reads mapped to rRNA and tRNA region obtained from UCSC genome browser were removed using Bedtools. For 30, 60, and 120 min timepoint data, based on the Pol2 elongation rates, maximum 30, 90, and 240 Kb gene body regions from TSS were used for differential gene expression (DGE) analysis. The number of reads mapped to defined regions was counted using HTseq. Log2 fold-change (FC) and p values were calculated in individual time points using DEseq2^89^ with the default scaling method (median of relative abundance). The fold changes from distinct timepoints were summarized by calculating the averages. Enhancer dependencies were determined by enhancer regulation based on the EU-seq DGE changes after the A-485 treatment. Expressed genes in each cell line were determined as described in the reference genome annotation section and further filtered the low expressed genes if the mean EU-seq read count of control and A-485 treatment condition is less than 20. ND as following criteria: < 1.2-fold changes of down-regulation after A-485 treatment. SD: < 1.5- and >= 1.2-fold changes of down-regulation. ID: < 2- and >= 1.5-fold changes of down-regulation. HD: >= 2-fold changes of down-regulation. For gene track visualization, we used deeptools and bigWigMerge (UCSC tool)^90^ to generate bigwig files using scaling factor calculated from DESeq2.

### ChIP-Seq

Reads were mapped to the reference genome by using bwa aln with default parameters (BWA version 0.7.10). Multi-mapped reads, duplicated reads, or reads with more than three mismatches were removed by samtools^88^. Reads mapped to the DAC Blacklisted Regions (https://www.encodeproject.org/annotations/ENCSR636HFF/) were omitted from the downstream analysis. Peak regions were called using LanceOtron with the default model (wide-and-deep_jan-2021)^91^. The peaks proximal within 2kb are merged using Bedtools^92^, and poorly enriched peaks of maximum peak height < 8 reads mapped per million (rpm) were omitted. Peak height was calculated using bamCompare^93^ with the following parameters (centerReads, minMappingQuality 10, 20bp bin, smooth length 400bp, extend reads 200bp, rpm normalization, and input rpm value is subtracted). Where indicated, peak regions were classified into three classes “Promoter” (TSS ±1kb), “Gene body” (Exon, intron, 5ʹ UTR, and 3ʹUTR, excluding “Promoter” regions), and “Distal” by using ChIPseeker^94^. IGV^95^ was used for the gene track visualization. For Pol2 ChIP-seq, we applied scaling factors based on the EU-seq results. We had assumed that Pol2 binding remains unaffected at promoters of genes that are not downregulated (ND) by A-485 in EU-seq analyses, and the scaling factors are calculated by DE seq default normalization methods using the Pol2 ChIP read counts at ND gene promoters (±2kb from TSS). For the quantification of Pol2 binding changes in the gene body regions, a maximum of 90kb gene body regions from TSS were used as described in the EU-seq section.

### Defining enhancer strength

Enhancer regions were defined as overlapping regions of H2BK20ac and H3K27ac peaks. For H2BK20ac-based analyses, enhancer strength was calculated by subtracting input rpm from the H2BK20ac rpm value. For H3K27ac-based analyses, enhancer strength was calculated by subtracting input rpm from the H3K27ac rpm value.

### Prediction model for ND and HD promoters

Support vector machines with gapped k-mer kernels (gkm-SVMs)^96^ were used to learn predictive models of ND and HD promoter sequences. Genes categorized as ND in all the cell lines (mouse: ESC, EpiSC and NPC, human: HCT116, HepG2, Jurkat, K562, KG-1, and SH-SY5Y) are defined as combined ND genes (mouse: N= 5603, human: N= 3258), and genes categorized as HD genes in any cell lines are defined as combined HD genes (mouse: N=3499, human: N=3154). Combined SD and ID genes are determined by the highest down-regulation of EU-seq DGE after A-485 treatment in all the cell lines; combined ID: < 2- and >= 1.5-fold changes of down-regulation in any cell lines and not classified as combined HD genes, combined SD: < 1.5- and >= 1.2-fold changes of down-regulation regulation in any cell lines and not classified as combined HD nor combined ID genes. DNA sequences of 100, 250, 500, and 1Kb upstream or downstream from TSS were used for generating prediction models. Null sequences were obtained by randomly selecting repeat- and GC-matching sequences from the corresponding genome using the genNullSeqs function of R package gkmSVM^96^. The approximately equal number of positive and negative sequence sets were used as input (<10% batch size difference). We tested gkm kernels and gkm radial basis function (gkm RBF) kernel using the gkmtrain function (lsgkm 1.10)^97^. The model performance was evaluated by the area under the precision-recall curve (AUPRC; calculated using ROCR R package)^98^ by 10-fold cross-validation. Gkm kernel trained by sequences of upstream 500bp from TSS with word length 10, maximum mismatch 3 and informative column 7 was chosen as the best model based on the AUPRC. The trained model was used for scoring all the expressed gene promoters by gkmpredict function (lsgkm)^97^.

Nucleotide contribution scores were calculated for all the nucleotides in all the positive and dinucleotide shuffled negative sequence sets (gkm explain function from lsgkm). We used TF– Modisco^99^ to obtain de novo motifs from the contribution scores of positive and negative sequence sets. We used TOMTOM (version 5.4.1)^100^ for comparing the computed motif position weight matrix (PWM) with the motif references. The motif references included JASPAR2022_CORE_vertebrates_non-redundant, uniprobe_mouse, jolma2013, HOCOMOCOv11_core_HUMAN_mono (for human motif), HOCOMOCOv11_core_MOUSE_mono (for mouse motif), and JASPAR2018_POLII.

### Annotation of CpG island, TATA box, and Inr promoter elements

250bp upstream and downstream sequences from TSS were used to calculate GC content and CpG dinucleotide frequencies. CpG island was defined as GC content > 0.5 and observed CpG/expected CpG > 0.6 across the 500 bp region. TATA box and Inr promoter elements are referenced from Eukaryotic Promoter Database (EPD)^101^.

### Overlap of promoter regions with SuRE peaks

K562 SuRE peaks were downloaded from the supplemental section of van Arensbergen et al. ^63^. SuRE^+^ active and inactive promoter, all the promoter regions in the reference genome annotation were classified into SuRE^+^ and SuRE^-^ based on the criteria where it is present within the range of -250 – +50bp from TSS. The proportions of SuRE^+^ active and inactive promoters are calculated by gene basis. SuRE peak overlap with A-485 downregulated gene class promoter was calculated by promoter-basis.

### Evaluating chromatin accessibility of COF-AID STARR-seq regions

COF-AID STARR-seq regions were downloaded from the supplemental information of Neumayr et.al. ^21^. HCT116 ATAC-seq data were downloaded from the GEO portal (GSE97889). Accessibility was calculated by maximum peak height and normalized reads density within STARR-seq regions as described in the ChIP-seq section.

### Gene classification by cell type-specificity

We used the FANTOM5 CAGE dataset as a reference to classify cell-type specificity of genes ^69, 102^. In the mouse reference, we excluded data from “whole,” “lactating,” and “pregnant” stages. The resultant 151 mouse tissue expression profile (44 tissues from 11 developing stages) and 76 human tissue expression profiles (68 tissues from adult and fetal stages) were converted into a binary expression matrix using a gene expression threshold of TPM ≥ 2. Genes were classified in mouse and human cell lines using the following criteria. [mouse] HK: genes expressed in all the three used cell lines and expressed in >= 80% of tissues in the CAGE dataset. CTE: genes expressed in all the three used cell lines and expressed in < 80% of tissues. CTS: genes expressed in =< 2 used cell lines. [human] HK: genes expressed in all the six cell lines and expressed in >= 80% of tissues. CTE: genes expressed in 4 or 5 used cell lines and expressed in >= 80% of tissues or genes expressed in 6 used cell lines and expressed in < 80% of tissues. CTS: genes expressed in =< 3 (out of 6) used cell lines.

### Statistical analysis

Otherwise mentioned, p-values are calculated using Mann–Whitney U test and corrected for multiple comparisons using Benjamini & Hochberg method (R package stats version 3.6.2).

### Use of publically available data

The following publicly available data were used: Mouse and Human gene annotations were from Genecode (https://www.gencodegenes.org); FANTOM5 CAGE dataset was from ArrayExpress (https://www.ebi.ac.uk/arrayexpress/); Promoter Elements were referenced from EPD (https://epd.epfl.ch//index.php); Motif references were downloaded from MEME Suite (https://meme-suite.org/meme/index.html) and JASPAR (https://jaspar.genereg.net/). The following publicly available sequencing datasets were reanalyzed: the RNA-seq in KG-1 (GSE171552), HepG2 (GSE134745), Jurkat (GSE134745), K562 (GSE134745), SH-SY5Y (GSE162644) and HCT116 (GSE144165); EU-seq in ESC treated without and with A-485 (GSE146328); the ChIP-seq on ZFP143 (GSE39263), NRF1 (GSE678674), GABPA (GSE116704), INO80 (GSE158533), YY1 (GSE178982), CTCF (GSE178982) in ESC; H3K4me3 in K562 (GSE163049), HepG2 (GSE111000), HCT116 (GSE127960), Jurkat (GSE164184), and SH-SY5Y (GSE90045), ATAC-seq of HCT116 (GSE97889).

### Cloning of plasmids for genome editing

Oligonucleotide pairs encoding sgRNAs (primers 1-14) were phosphorylated, annealed, and cloned into plasmids pX330 or pX458 by one-pot BbsI digestion/T4 DNA ligation as described^103^ followed by exonuclease digestion. Oligonucleotide pairs (primers 15-26) encoding intronic genomic target sites for the *Nanog* enhancer insertion (in proximity to *Mcmbp*, *Ick*, *Sec23ip*, *Foxo9*, *Mthfsd*, and *Foxc2*) were phosphorylated, annealed, and inserted by T4 DNA ligation into the pRRP recombination reporter^104^ digested with SacI and AatI. Templates for HDR-based editing were cloned by amplifying genomic regions with KOD Xtreme Hot Start DNA (primers 27-42) followed by amplification for homology arms with added multiple cloning site regions using PrimeStar MAX DNA polymerase (primers 43-88) and the pieces assembled by Gibson assembly into a linearized GeneArt pMA vector. A casette encoding an N-terminal Puromycin-P2A was inserted by restriction/ligation cloning to the homology arms containing loxP sites around the *Rif1* enhancer. For the degron editing of *Rif1*, a cassette encoding N-terminal Puromycin-P2A-GFP-FKBP12^F36V^ was similarly inserted to a Rif1 N-terminal homology region. A segment of the *Nanog* enhancer was inserted by digestion/ligation cloning into homology regions proximal to *Mcmbp*, *Sec23ip*, *Ick*, *Fbxo9*, *Foxc2*, and *Mthfsd*. A list of oligonucleotides used for genome editing, genomic confirmation of clones, and RT-PCR is provided in the Supplemental Table 1.

### Genome editing of mESC

For generating Cre-ERT2 expressing mESCs (E14TG2a, Sigma-Aldrich, cat# 08021401) were stably transfected with a PiggyBac CreERT2-expressing plasmid^105^. For generating *Rif1* enhancer fl/fl cells, the Cre-ERT2 expressing cells were seeded to gelatin-coated wells in a 6-well plate.

The day after, 1250ng pX330 and 1250ng HDR templates were mixed with 7.5µL Lipofectamine 2000 in a total volume of 500µL OptiMEM, allowed to stand 20min and then added to the cells. After 24h transfection, cells were trypsinized and split 1:20 to a gelatin-coated 10cm dish. After attaching 24h, the cells were selected with 0.5µg/mL puromycin for the next 6-8 days. Correctly modified clones were identified by genotyping PCR with KOD Xtreme Hot Start DNA polymerase.

For inserting *Nanog* enhancer in proximity to *Mcmbp*, *Fbxo9*, *Ick*, *Sec23ip*, *Mthfsd*, and *Foxc2*, mESC (E14TG2a, Sigma-Aldrich, cat# 08021401) were seeded to gelatin-coated wells in a 6-well plate for each transfection. The day after 200ng pRRP reporter, 1000ng pX330/458, and 1500ng HDR template with the *Nanog* enhancer were transfected with lipofectamine and OptiMEM as outlined above. 24h after transfection cells were trypsinized and split to 1:5 and 1:20 in gelatin-coated 10cm dishes. 2-3h after splitting, cells were transiently selected (24h) with puromycin (2µg/mL). Subsequently, the cells were washed once with PBS, and fresh media, without puromycin, was added. Colonies were picked 6-8 days after transfection and genotyped as described above (primers as for the cloning, or 91-94).

### mRNA isolation and RT-qPCR for mRNA quantification

Total RNA was isolated using an RNeasy mini kit (Qiagen) as described for EU-seq sample collection. Isolated RNA (500 ng) was reverse-transcribed to cDNA by using PrimeScript RT master mix (Takara). PCR was performed using Mx3005P (Agilent) with SYBR® Premix EX Taq^TM^ II (Takara) and primers listed in the table (99-112). The relative expression levels were calculated using *β-actin* mRNA as a reference.

### Confirmation of *Rif1* enhancer deletion and RIF1 depletion, and cell proliferation analysis

Following trypsinization, *Rif1* enhancer fl/fl or or GFP-FKBP12^F36V^-RIF1 mESC were re-seeded in gelatin-coated wells in 6-well plates with media supplemented with DMSO (1:1000) (both cell lines), 4-OHT (100nM) (*Rif1* enhancer fl/fl cells), or dTAG-13 (100nM) (GFP-FKBP12^F36V^-RIF1 cells). The cells were imaged 3 and 5 days after the treatment to monitor colony growth. For confirming dTAG13-induced depletion of GFP-FKBP12^F36V^-RIF1 cells, 1.2e4 cells were seeded in a Geltrex coated 96-well CellCarrier Ultra plate and allowed to attach overnight. The next day, the cells were stained with SiR-DNA (0.1µM) for 1h. Then dTAG-13 (100nM) was added to the cells. The cells were imaged immediately after adding d-TAG13 and again after 1h of dTAG-13 addition, on a Perkin Elmer Phenix Harmony confocal spinning disk microscope using 488nm and 647nm excitation.

To quantify cell viability, the cells were seeded in triplicates in 6-well plates as described above. After 3 days of treatment, the medium from the supernatant was collected, the cells washed once with PBS, wash collected, followed by TrypLE dissociation for 2-3min. Dissociated cells were collected in the same tube as the supernatant and the PBS wash. The wells were washed once more with PBS and collected in the same tubes followed by centrifugation 2min at 400g. The cells were dissociated with a 20µL pipette tip and resuspended in 1mL medium followed by dilution 1:2 with 0.4% trypan blue and loading of 10µL to a Countess cell culture chamber slide and quantification on a Countess 3 cell counter (Thermo Scientific).

---

**Extended Data Fig. 1.**
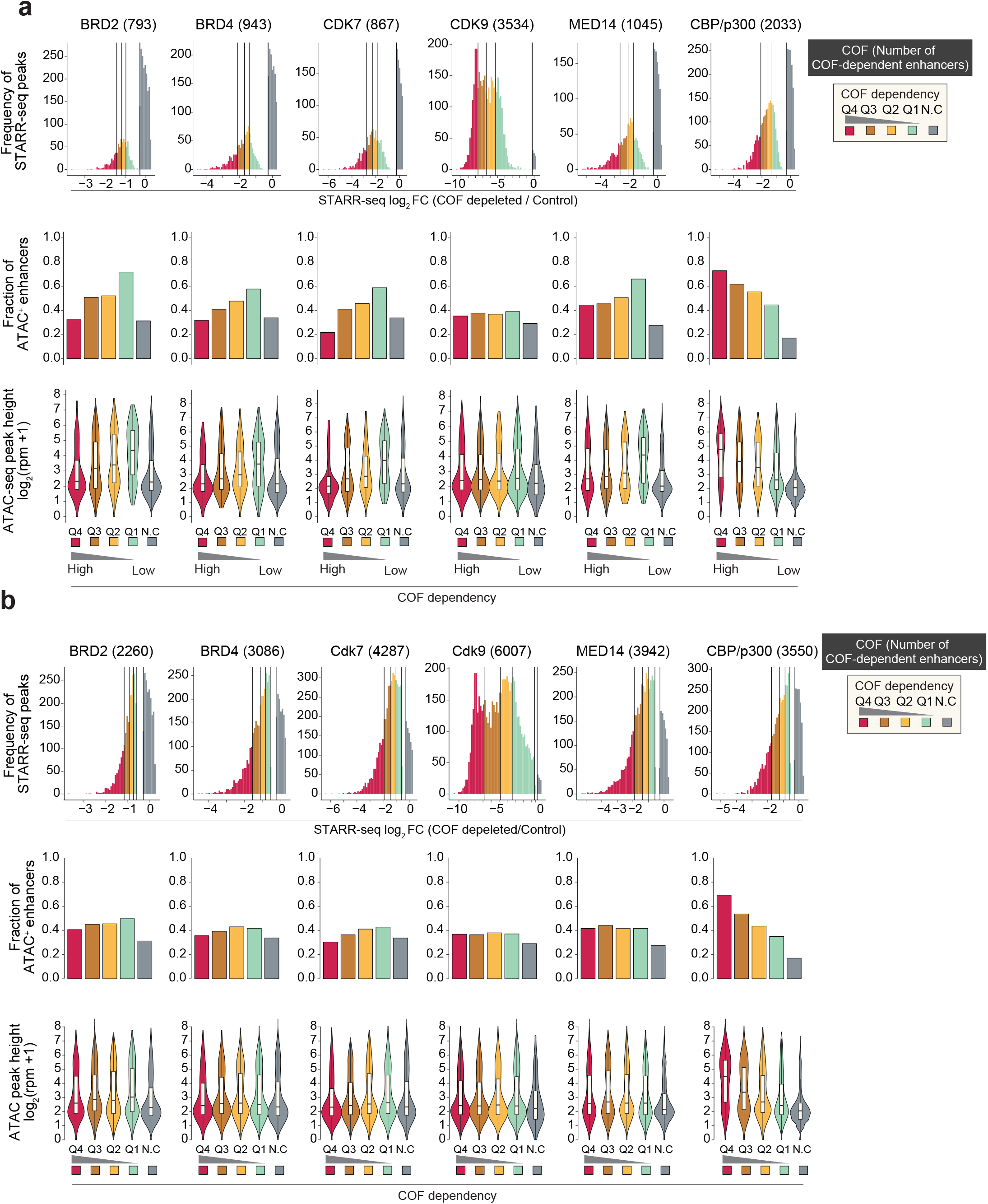
CBP/p300 is required for enhancer-dependent transcription activation in STARR-seq. **a,** The requirement of the six indicated cofactors (COFs; BRD2, BRD4, CDK7, CDK9, MED14, CBP/p300) for enhancer activity was assessed by Neumayr et al. using STARR seq21. The activity of 6249 candidate enhancers was quantified in wild-type cells or cells depleted of the indicated COFs. Based on the STARR-seq signal, the relative change in enhancer activity in wild-type and COF depleted cells was determined. For each COF, the number of enhancers that showed decreased activity in COF-depleted cells is indicated. The number of enhancers showing COF dependency was determined using statistical significance (p=<0.05) (shown in panel a) or fold downregulation (>1.5 fold decrease in enhancer activity in COF depleted cells) (shown in panel b). Based on enhancer activity fold-change in COF-depleted cells, enhancers were grouped into quartiles. Enhancers showing no change (N.C.) in activity after the depletion of the indicated COFs are shown as a separate group. In each of the above enhancer groups, native chromatin accessibility was analyzed in the cell line in which the enhancer activity was assayed (HCT116). Chromatin accessibility was determined by ATAC-seq. In the top panel, the frequency plot shows the fold-change distribution of enhancer activity in COF depleted and wild-type control. In the second raws, bar charts show the fraction of enhancers that are ATAC-seq^+^ in HCT116. In the last raw, violin plots show DNA accessibility by ATAC-seq in the corresponding group of enhancers. The fraction of enhancers showing a decrease in STARR-seq activity after the depletion of the indicated COFs is indicated. Analyses in panel a include enhancers whose activity in MPRA is statistically significantly decreased in the indicated COF depleted cells. Analyses in panel **b** include enhancers whose activity in MPRA is decreased by >1.5 fold (regardless of statistical significance) in the indicated COF depleted cells. Note that, among all analyzed COFs, CBP/p300 is the only COF that shows a strong association with the accessibility of enhancers in the native context. Enhancers whose activity is most strongly decreased after CBP/p300 depletion have the highest accessibility, enhancers that show intermediate dependency on CBP/p300 have intermediate accessibility, and enhancers that are insensitive to CBP/p300 depletion either lack or have very low DNA accessibility.

**Extended Data Fig. 2.**
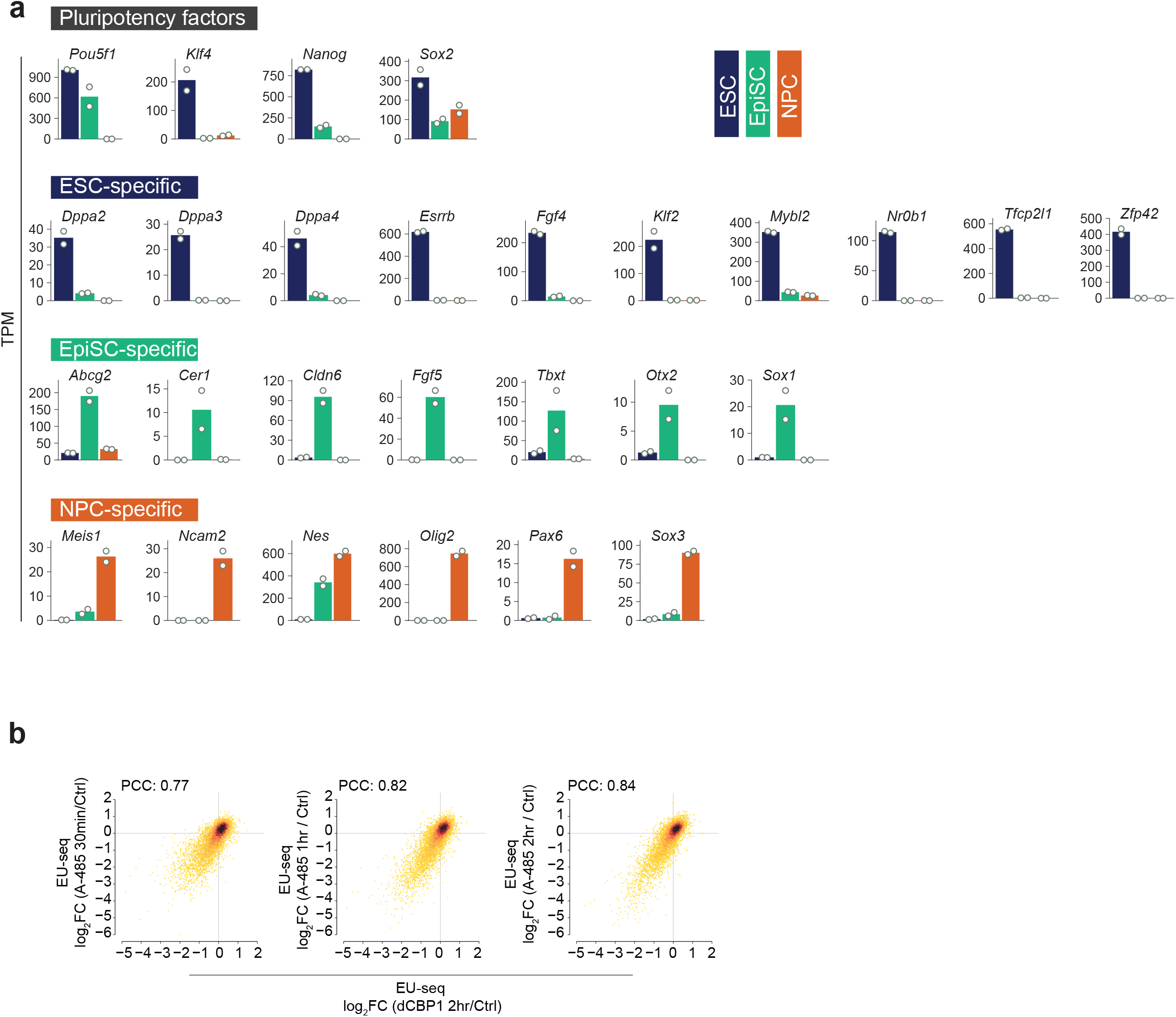
Confirmation of ESC, EpiSC, and NPC cell identity, and validation of A-485 target specificity. **a,** Expression profiles of cell type-specific transcription factors and signaling proteins in ESC, EpiSC, and NPC. Shown are the mRNA expression levels of the indicated genes, among the specified cell lines. Gene expression levels were quantified from two independent RNA-seq experiments. (TPM; transcripts per million). Two biological replicates were performed for RNA-seq in ESC, EpiSC, and NPC. **b,** CBP/p300 catalytic inhibition and acute CBP/p300 depletion cause similar gene transcription changes. ESCs were treated with the catalytic inhibitor A-485 for 30 min, 1h, and 2h, and with the CBP/p300 PROTAC dCBP1 for 2h. Nascent transcription changes were quantified using EU-seq. Shown are correlations between A-485 and dCBP1-induced transcription changes at the indicated time points. Note that A-485 and dCBP1-induced transcription changes are highly correlated, and A-485 shows no overt signs of off-target specificity. EU-seq data for A-485 treated ESC were re-processed from^34^. dCBP1-induced nascent transcription change was analyzed from a single experiment. PCC: Pearsońs correlation coefficient.

**Extended Data Fig. 3.**
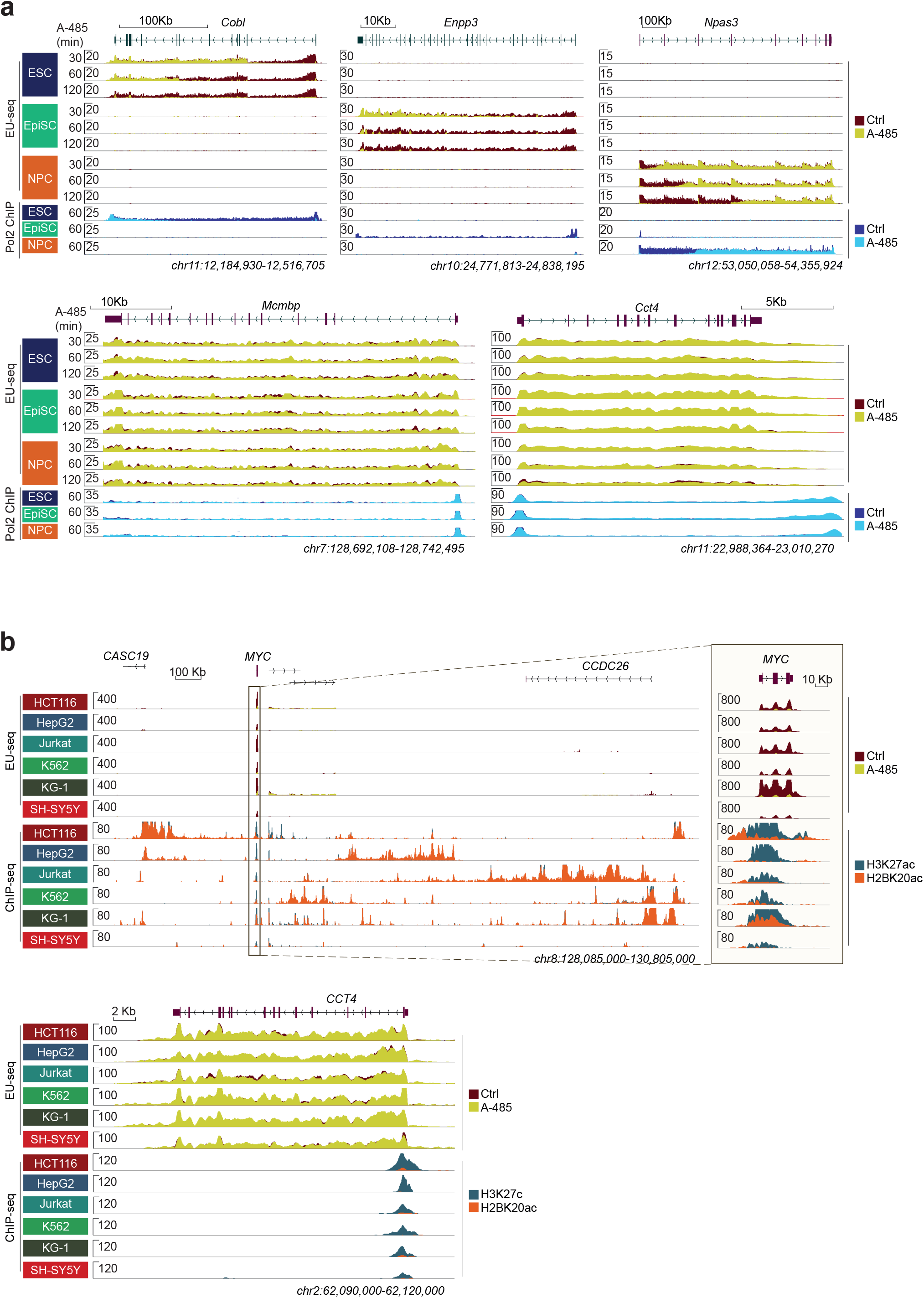
Confirmation of A-485-induced selective downregulation of enhancer-regulated genes in mouse and human cell lines. **a,** Representative nascent transcription tracks (top panels) in mouse cell lines showing A-485-induced downregulation of the indicated cell-type-specific, enhancer-dependent genes in (*Cobl, Enpp3, Npas3*), or housekeeping genes (*Mcmbp, Cct4*). The overlaid tracks show EU-seq signal in untreated and A-485-treated (30, 60, 120 min) cells, as indicated. Relatively long cell-type-specific genes were chosen as representative examples to demonstrate a time-dependent increase in the length of nascent transcript inhibition. The length of A-485-induced transcript inhibition roughly corresponds to the speed of Pol2 elongation. The overlaid Pol2 data (bottom panels) show Pol2 binding in the indicated treatment conditions and cell lines. Pol2 binding was analyzed by ChIP-seq, without or with A-485 treatment (60 min). Reduced Pol2 binding in ChIP-seq analyses independently confirms the above EU-seq results. *Mcmbp* and *Cct4* remain unaffected by A-485 in EU-seq and Pol2 ChIP-seq analyses in all cell lines. For ESC, EU-seq data were re-processed from^34^. EU-seq analyses in EpiSC and NPC were performed in minimum 3 independent biological replicates, and Pol2 ChIP-seq in ESC, EpiSC, and NPC were performed in two biological replicates. b, Confirmation of A-485-induced selective gene downregulation in human cell lines. Genome browser view of nascent transcription expression in the indicated human cell lines, and treatment conditions. For each cell line, overlaid tracks show EU-seq signals in untreated and A-485 treated conditions. Note that A-485 downregulates enhancer target (*MYC*), but not housekeeping (*CCT4*), genes. Nascent transcription was analyzed in the indicated 6 cell lines using EU-seq, without or with A-485 (60min). Two biological replicates were performed in all the indicated cell lines. Overlaid H3K27ac and H2BK20ac ChIP-seq tracks indicate the position of proximal candidate cis-regulatory elements.

**Extended Data Fig. 4.**
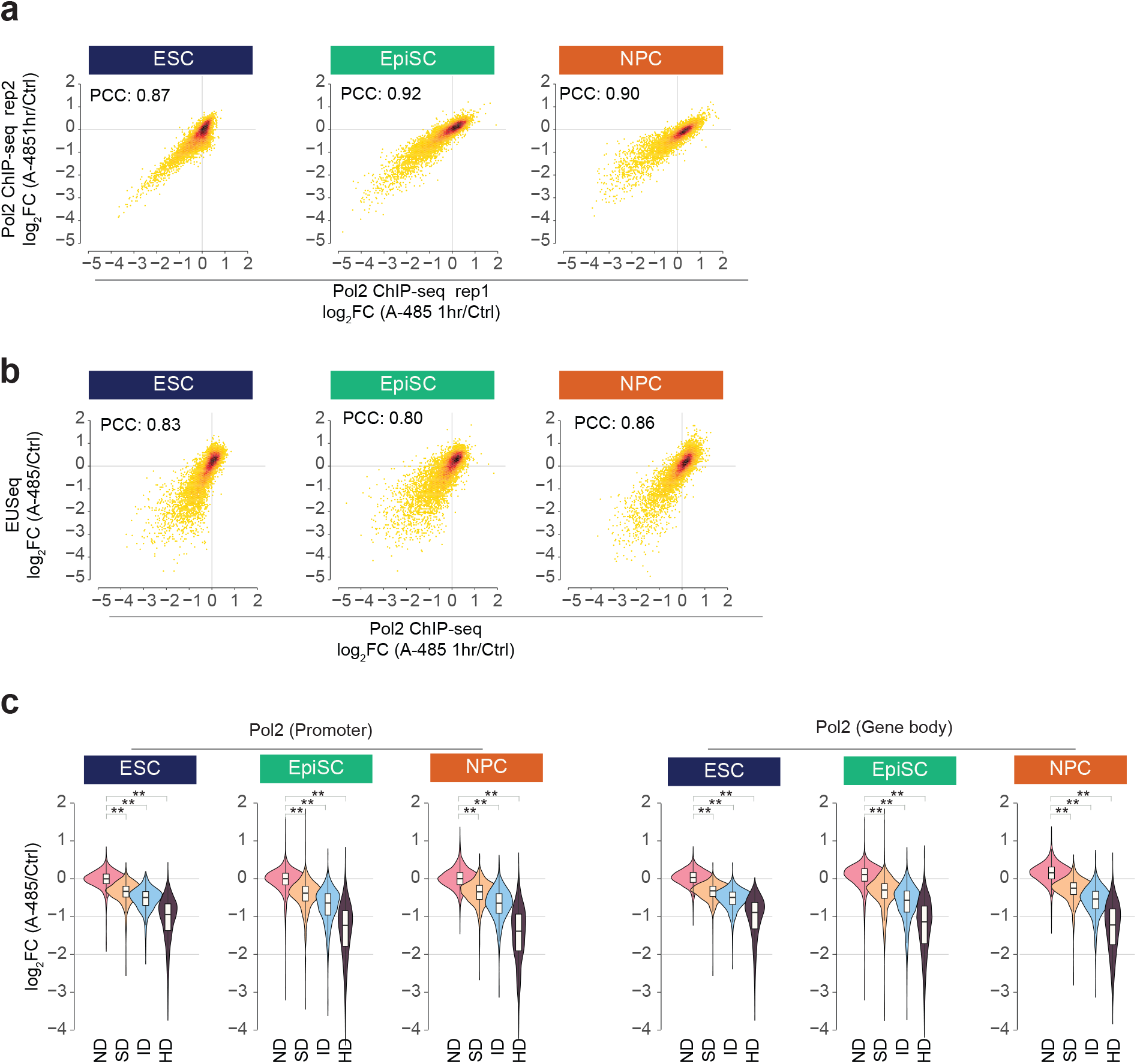
A-485-induced nascent transcription changes correlate with the changes in Pol2 binding. **a,** Shown are correlations between two independent Pol2 ChIP-seq replicate measurements in the indicated cell lines. Pol2 binding was analyzed without or with A-485 (1h) treatment. PCC: Pearsońs correlation coefficient. **b,** Correlation between A-485-induced nascent transcription changes (EU-seq) and Pol2 binding (Pol2 ChIP-seq) in the indicated cell lines. Note that, in all cell lines, A-485-induced change in Pol2 binding is strongly correlated with the change in EU-seq signal. **c,** A-485 treatment proportionally reduces Pol2 binding at A-485 downregulated gene class. Shown is the normalized Pol2 intensity in A-485 treated and untreated cells. Assuming that Pol2 binding remains unaffected at promoters of genes that are not downregulated (ND) by A-485 in EU-seq analyses, Pol2 binding untreated and A-485 treated cells were normalized based on Pol2 intensity at ND gene promoters (+/− 2kb from TSS). Shown are normalized Pol2 ChIP intensity at promoters (left panels) or gene body (right panels) regions in the indicated cell lines and the A-485 regulated gene class. A-485-regulated genes were classified based on A-485-induced nascent transcription regulation as defined in Fig. 1b. HD, highly downregulated (≥2-fold decreased); ID, intermediate downregulated (>1.5-2.0-fold decreased); SD, slightly downregulated (>1.2-1.5-fold decreased); and NC, not downregulated (≤1.2-fold decreased) after A-485 treatment. *** p < 1e-50, Mann–Whitney U test.

**Extended Data Fig. 5.**
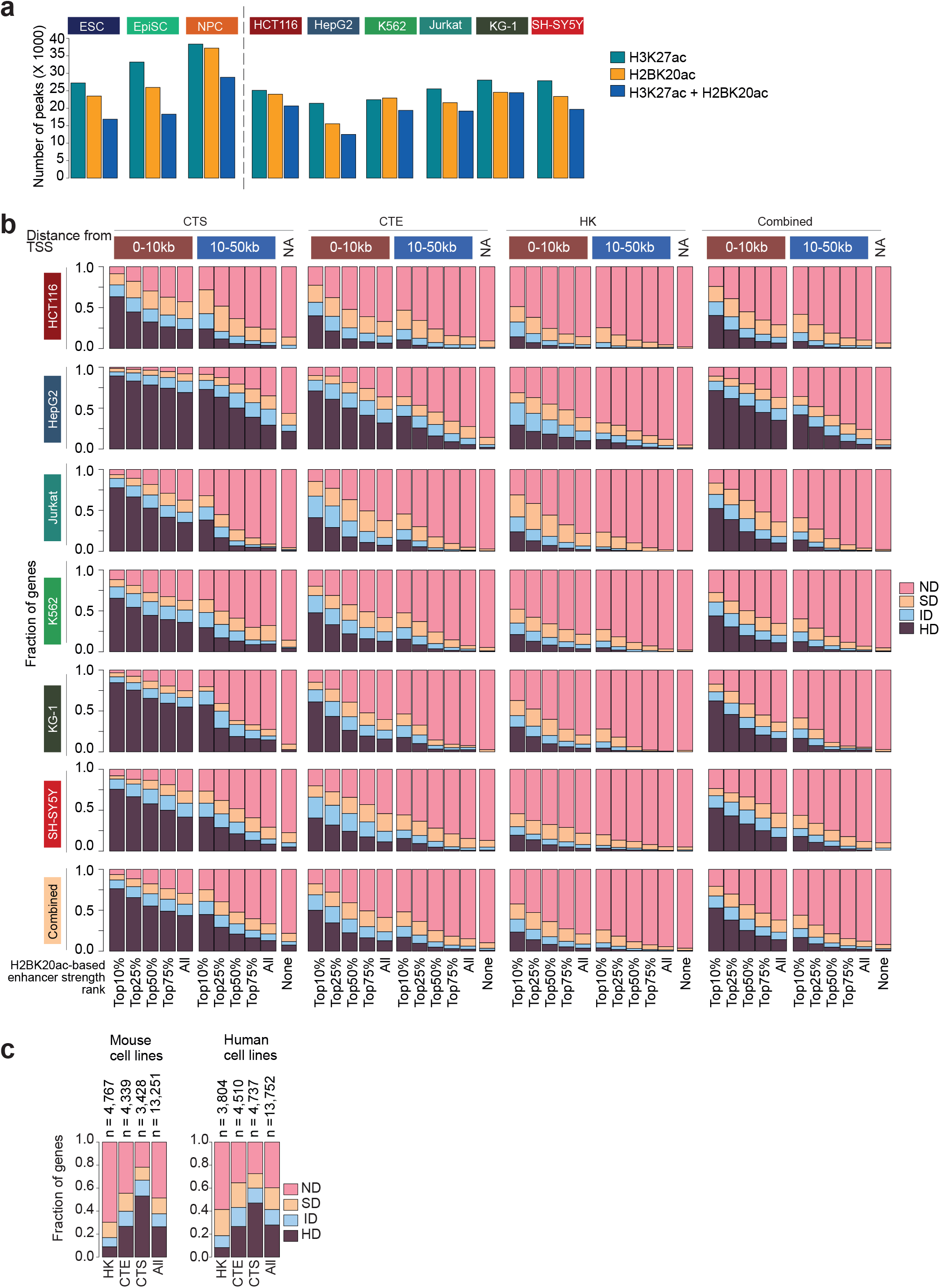
CBP/p300 inhibition downregulates enhancer proximal genes, and CTS shows greater enhancer dependence than HK genes. **a,** Shown are the number of H3K27ac and H2BK20ac ChIP-seq peaks identified, as well as the number of regions commonly marked with both H3K27ac and H2BK20ac, in the indicated cell lines. b, H2BK20ac intensity strongly relates to A-485-induced gene regulation in human cell lines. In the indicated cell lines, genes were classified based on their cell type-specificity as housekeeping (HK), cell-type-enriched (CTE), and cell-type-specific (CTE). A-485-downregulated genes were classified as defined in Fig 1b. Enhancers commonly marked with H2BK20ac and H3K27ac were ranked based on H2BK20ac intensity. Enhancers were grouped into Top10%, 25%, 50%, 75%, and All, based on H2BK20ac intensity. Enhancers were further grouped based on the distance from TSS of the nearest expressed genes. In each cell line, the bar charts show the relationship between cell-type-specificity class, enhancer rank category, enhancer distance, and A-485-induced gene regulation. Note that the fraction of A-485 downregulated genes progressively decreases with decreasing cell-type-specificity, decreasing enhancer strength, and increasing enhancer distance. In all cell-type-specificity categories, genes occurring in proximity to strong enhancers are more frequently downregulated by A-485 than genes occurring in proximity to weaker and distally located enhancers. c, Enhancer activity regulates a large fraction of mammalian genes. Fraction of A-485 regulated gene class among the specified cell-type-specificity categories in all mouse (left: ESC, EpiSC, NPC) and human (right: HCT116, K562, HepG2, Jurkat, KG-1, and SH-SY5Y) cell lines. The quantified genes were ranked based on their A-485-induced regulation, and assigned to the maximum downregulated category. For example, if a gene was downregulated ≥2-fold in one cell line but was regulated < 2-fold in other cell lines, the gene was assigned to the highest downregulation category, based on the maximum fold downregulation. The cumulative bar charts show the distribution of A-485 down-regulated gene categories in the specified cell-type-specific groups. The numbers shown (top) indicate the number of non-redundant genes in each cell-type-specificity category, as well as the total number of non-redundant genes quantified. The ‘Combined’ category includes genes from CTS, CTE, and HK categories, as well as ‘NC’ genes that were not classified in any of these categories.

**Extended Data Fig. 6.**
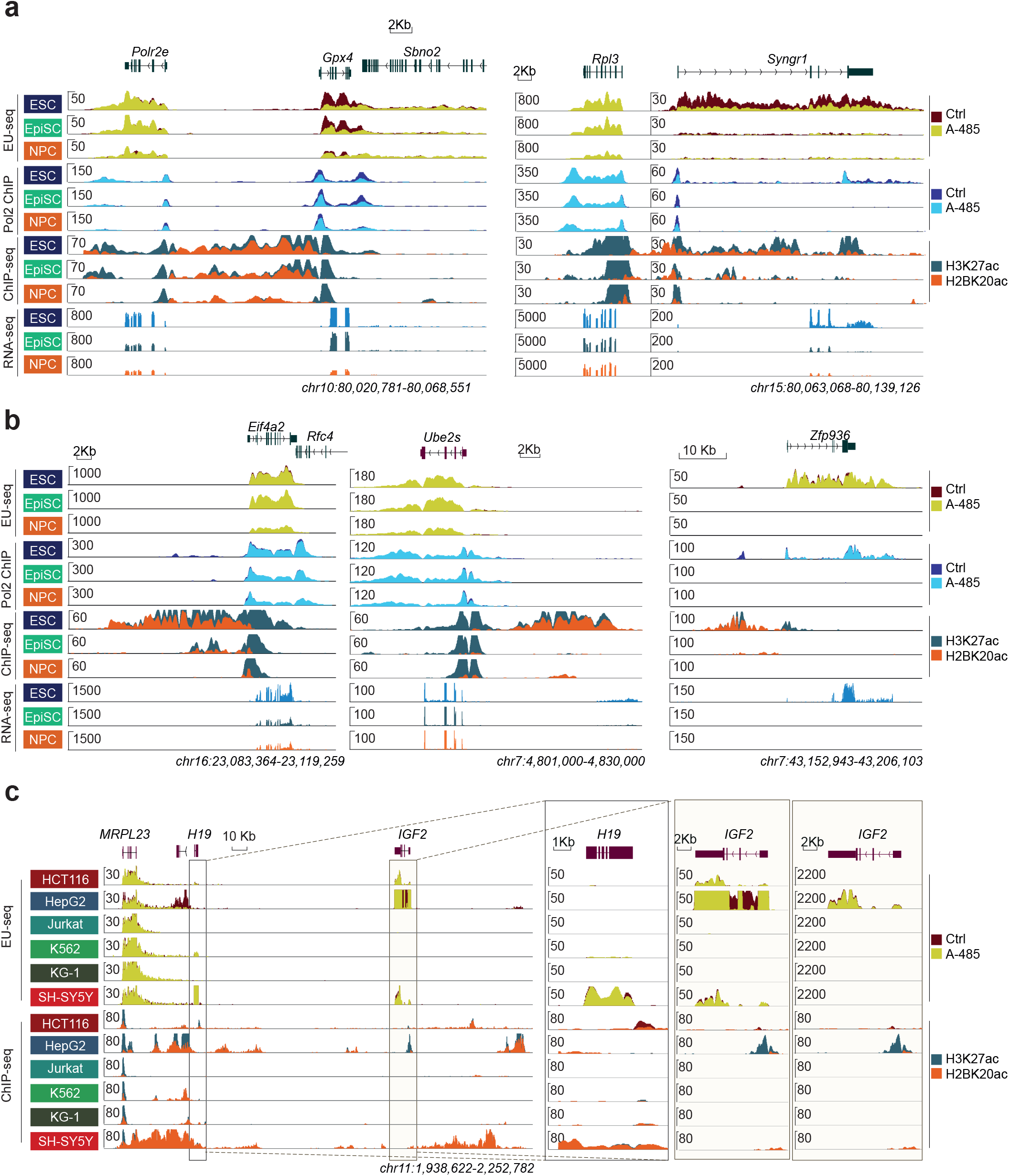
Gene-selective sensitivity to proximal enhancers. **a,** Representative examples of gene pairs that occur in proximity (<10kb) enhancers, but one of them is downregulated by A-485. In ESC and EpiSC, enhancer(s) is located between *Polr2e* and *Gpx4*. Only *Gpx4*, which is more closely positioned to the enhancer, is downregulated. Similarly, *Rpl3* and *Syngr1* occur in proximity to enhancers in ESC, but only *Syngr1*, which is more proximally positioned to the enhancers, is downregulated. Targeting of enhancers to *Gpx4* and *Syngr1* may explain the A-485 insensitivity of the other proximal genes. **b,** Examples of ubiquitously expressed (*Eif4a2* and *Ube2S*) and cell-type-specific (*Zfp936*) genes that appear insensitive to proximal enhancers. These genes represent exceptions whose enhancer insensitivity remains unexplained. **c,** 3D genome organization can promote preferential enhancer-promoter pairing, and genes with short (or none) intron sequences and stable mRNA can falsely appear as enhancer-insensitive in nascent transcription analyses. Through 3D genome organization, enhancers located between *MRPL23* and *H19* connect with and regulate *H19* and *IGF2*^41^, which may explain the A-485-insensitivity of *MRPL23*. Because of the long mRNA half-life, in EU-seq analyses, the overall expression of H19 and *IGF2* is not downregulated after A-485. But a zoom-in view of the *IGF2* intron region unambiguously demonstrates its A-485-induced downregulation. Note that at a higher scale, TPM 2200, overall IGF2 expression does not appear regulated by A-485. At a lower scale, TPM 50, EU-seq signal at IGF2 introns is abolished in A-485 treated SH-SY5Y, showing that the gene is indeed downregulated.

**Extended Data Fig. 7.**
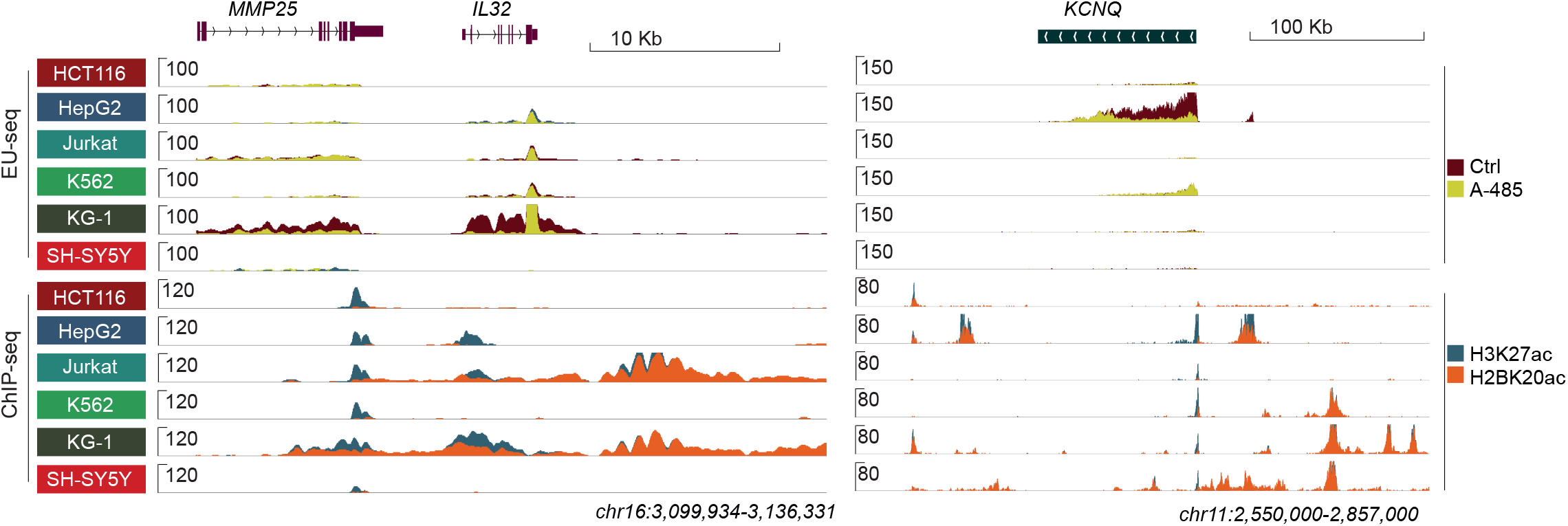
Cell-type-specific differences in regulation of genes with proximal enhancers. *IL32* harbors strong downstream enhancers in Jurkat and KG-1, as well as upstream enhancers in KG-1. However, *IL32* is more strongly expressed and downregulated by A-485 in KG-1 than in Jurkat. Similarly, *KCNQ* has proximal enhancer-like features in multiple cell lines, yet the gene is more strongly expressed and A-485 downregulated in HepG2. These examples indicate that *IL32* and *KCNQ* promoters are compatible with enhancers, and additional factors, such as 3D genome organization, presence of insulators, or accessibility of promoters, contribute to their cell-type-specific regulation by enhancers.

**Extended Data Fig. 8.**
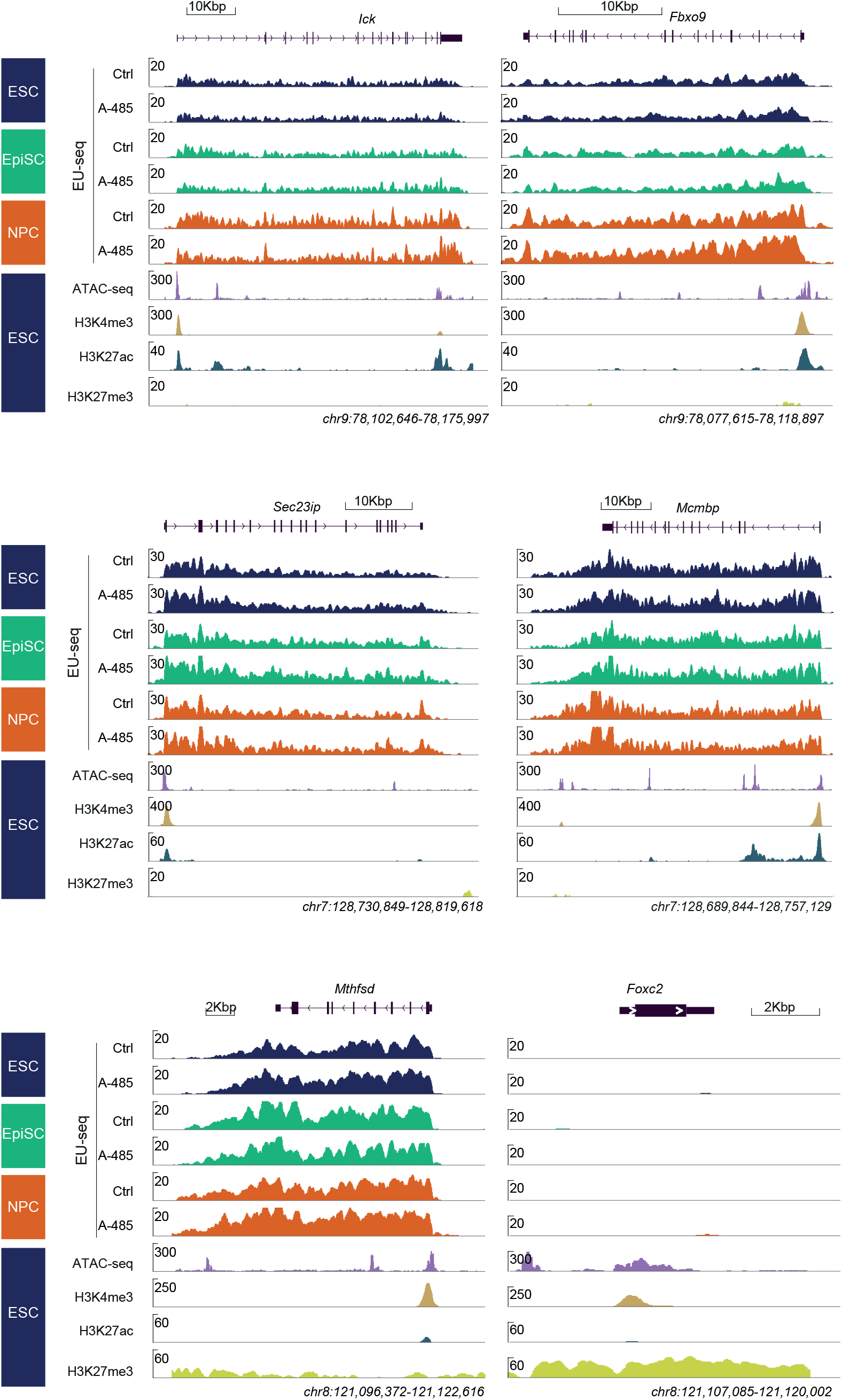
Expression of *Mcmbp, Ick, Fbox9, Sec23ip, Mthfsd,* and *Foxc2* in murine cell lines. Genome browser tracks show ubiquitous, enhancer-independent expression of *Mcmbp, Ick, Fbox9, Sec23ip, and Mthfsd,* in the indicated mouse cell lines. *Foxc2* is not expressed in any of the cell lines. *Foxc2* is marked with the repressive mark H3K27me3, but its promoter is accessible in ATAC-seq in mESC.

**Extended Data Fig. 9.**
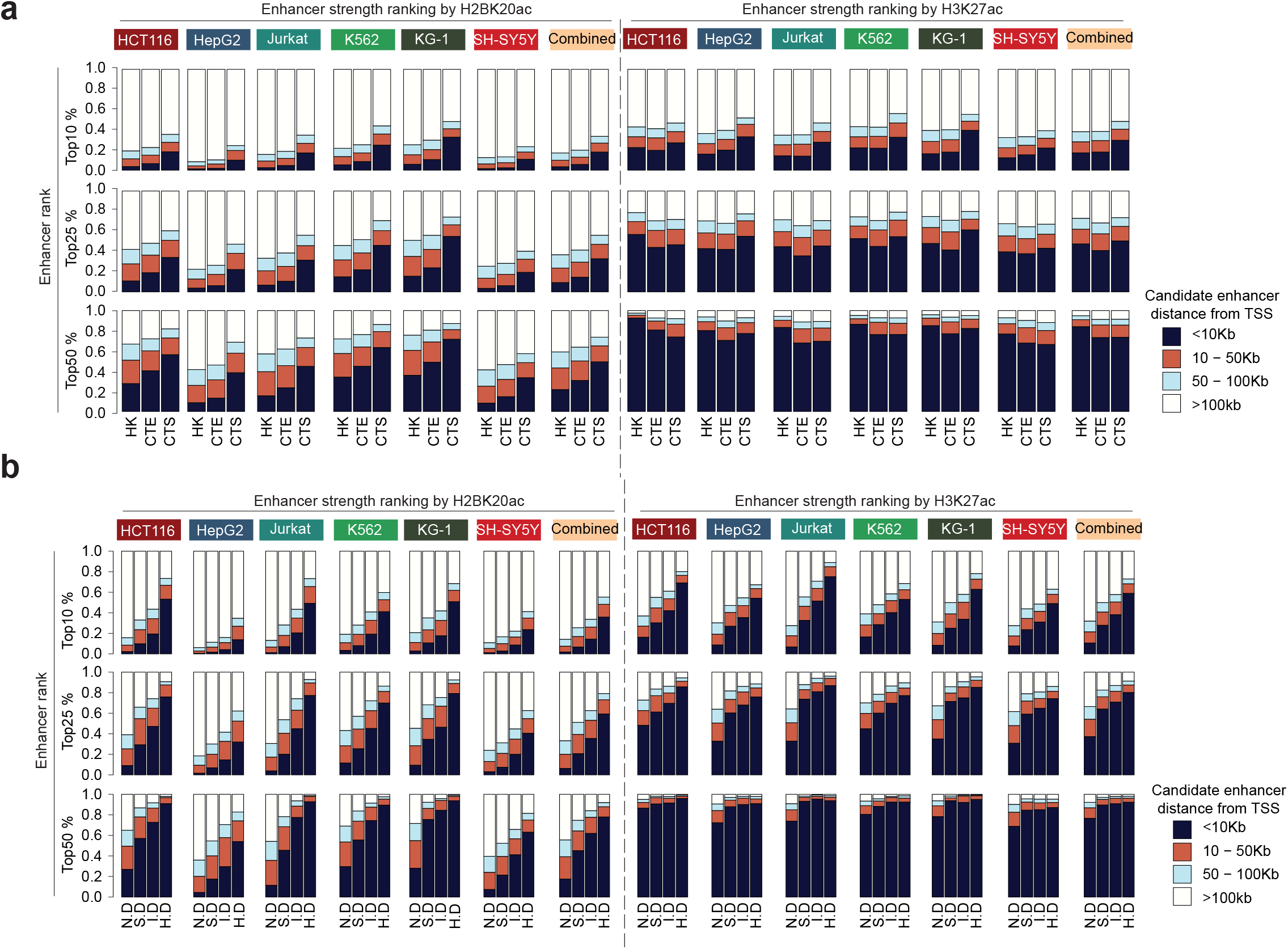
Enhancers are preferentially positioned in proximity to A-485 downregulated genes, enhancer-dependent gene regulation is associated with enhancer strength, distance, and cell-type specificity of genes. a, Genes expressed in the indicated human cell lines were classified based on their cell type-specificity (HK, CTE, CTS) as defined Fig. 1a (a) or A-485-induced downregulation (ND, SD, ID, HD) as defined in Fig1b (b). Enhancer strength was determined using H2BK20ac (left panels) or H3K27ac (right panels) intensity, as defined in Fig. 2. Based on the acetylation mark intensity, the Top50% of enhancers were ranked grouped into Top10%, 25%, and 50% categories. For calculating H2BK20ac-based enhancer strength, candidate enhancer regions commonly marked with H3K27ac and H2BK20ac were used in these analyses. Genes were further grouped based on the distance between the enhancer and the nearest TSS. The bar charts show the relationship between cell-type-specificity, enhancer rank category, and enhancer-TSS distance. Note that, in comparison to HK genes, a greater fraction of CTS genes harbor strong enhancers in their proximity (<10kb). Notably, unlike H2BK20ac intensity-based enhancer ranking, H3K27ac intensity-based enhancer ranking shows a poor relationship with cell-type-specificity, enhancer strength, and distance. b, Relationship between H2BK20ac (left panels) and H3K27ac (right panels) intensity-based enhancer ranking and A-485-induced gene regulation. In the indicated human cell lines, genes were classified based on A-485-induced downregulation (ND, SD, ID, HD) as defined in Fig. 1b. Enhancer strength was defined by the H2BK20ac or H3K27ac ChIP signal, Top50% enhancers were rank-ordered, and grouped into the indicated percentile categories. Candidate enhancers commonly marked with H3K27ac and H2BK20ac were used in these analyses. Genes were further grouped based on the distance between the enhancer and the nearest TSS. Within different groups, the fraction of genes in A-485 regulated categories is shown. A much greater fraction of HD genes harbors top-ranked H2BK20ac marked enhancers than ND genes, explaining why HD genes are more prominently downregulated by A-485 than ND genes. Of note, the relationship between H3K27ac intensity-based enhancer ranking and gene regulation rapidly deteriorates and in the Top50% rank category, H3K27ac-based ranking provides minimal distinction among different A-485 regulated gene classes.

**Extended Data Fig. 10.**
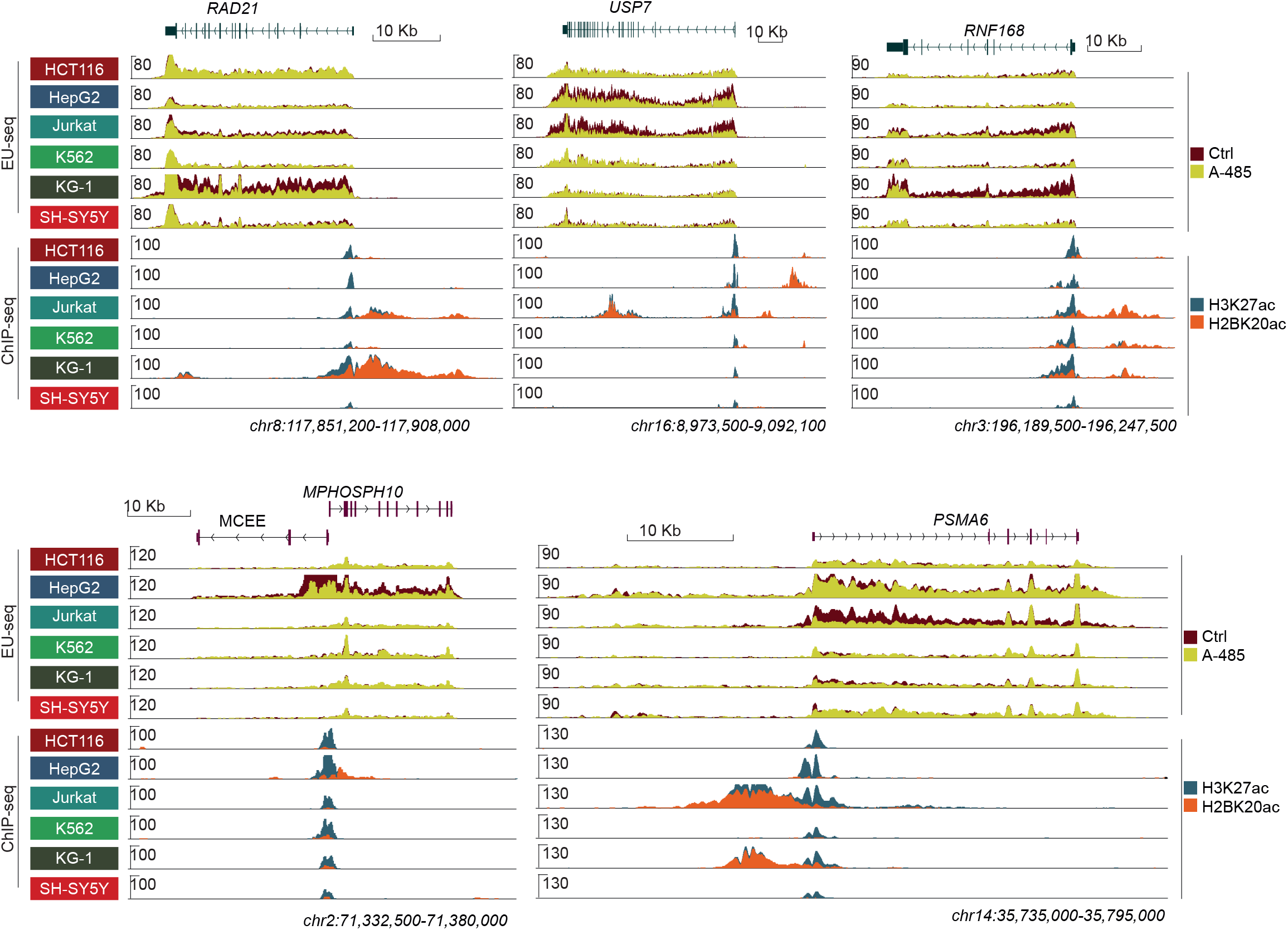
Enhancer-dependent quantitative boosting of housekeeping genes. Genome browser view of several housekeeping genes (*RAD21, USP7, RNF168, MPHOSPH10, and PSMA6*) showing cell-type-specific transcriptional boosting by enhancers. Note that cell line-specific enhancer-dependent boosting is associated with the occurrence of proximal enhancers marked with H2BK20ac.

**Extended Data Fig. 11.**
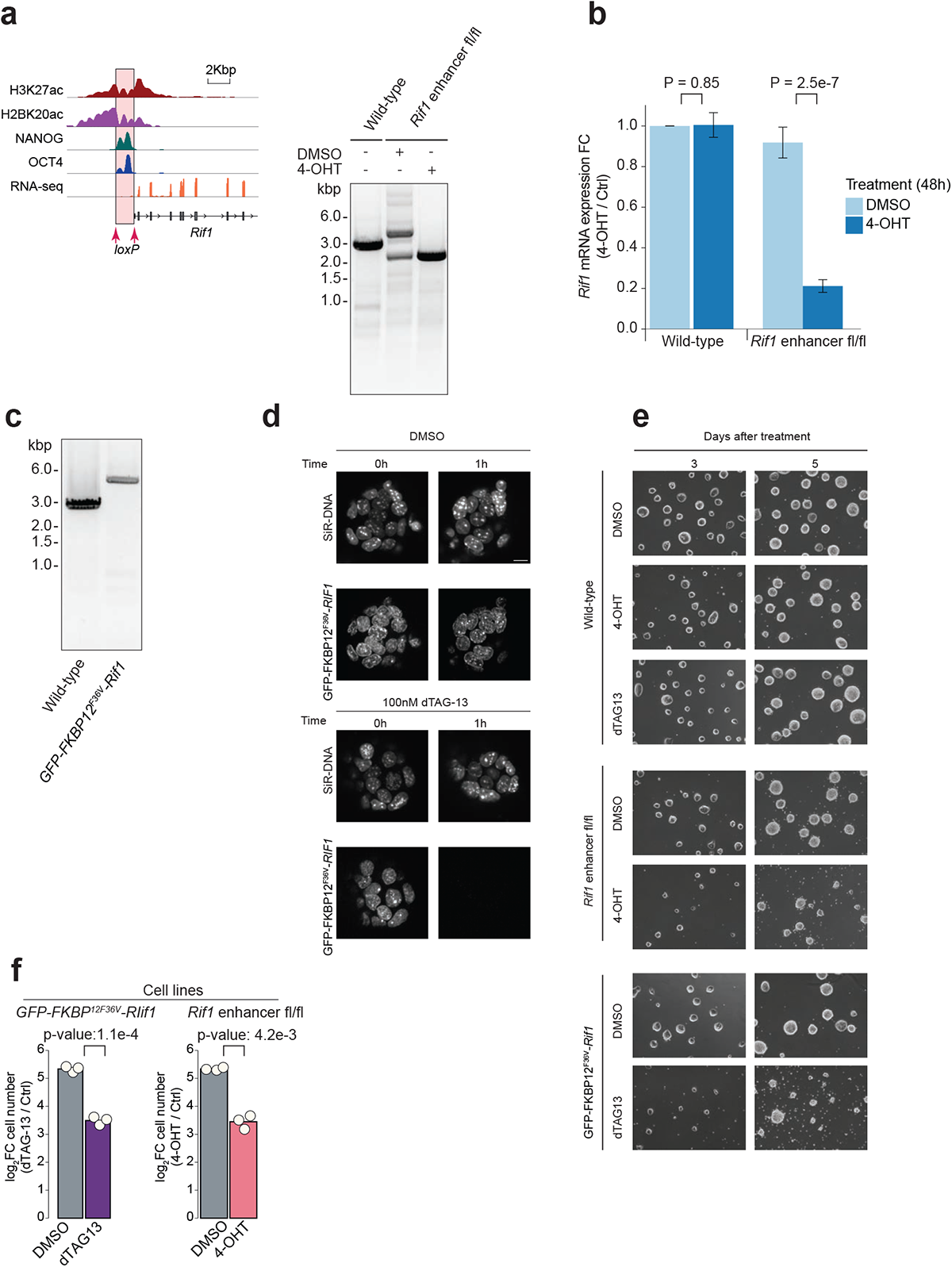
Enhancer-dependent RIF1 boosting is functionally relevant. Deletion of the upstream (-1kb) enhancer decreases *Rif1* expression in mESC. Using CRISPR, the *Rif1* upstream enhancer was flanked by loxP sites in mESC expressing Cre-ERT2 **(a)**. The enhancer was conditionally deleted by treating cells with 4-OHT (100nM, 48h) **(b)**. The bar charts show *Rif1* expression wild-type and Rif1 enhancer fl/fl cells, treated with DMSO or 4-OHT (100nM, 48h). **c-d,** Generation of GFP-FKBP12^F36V^-RIF1 expressing mESC **(c)**, and validation of GFP-FKBP12^F36V^-RIF1 degradation (dTAG-13, 100nM, 1h) **(d)**. **e-f,** Rif1 enhancer deletion impairs mESC proliferation. Rif1 enhancer fl/fl cells were treated with 4-OHT (100nM) to conditionally delete the enhancer, and GFP-FKBP12F36V-RIF1 cells were treated with dTAG13 (100nM) to conditionally deplete RIF1 protein. Cells were imaged 3 and 5 days after the treatment **(e)**, and the number of live and dead cells was counted on day 3 **(f)**. p-values: two-sided t-test.

**Extended Data Fig. 12.**
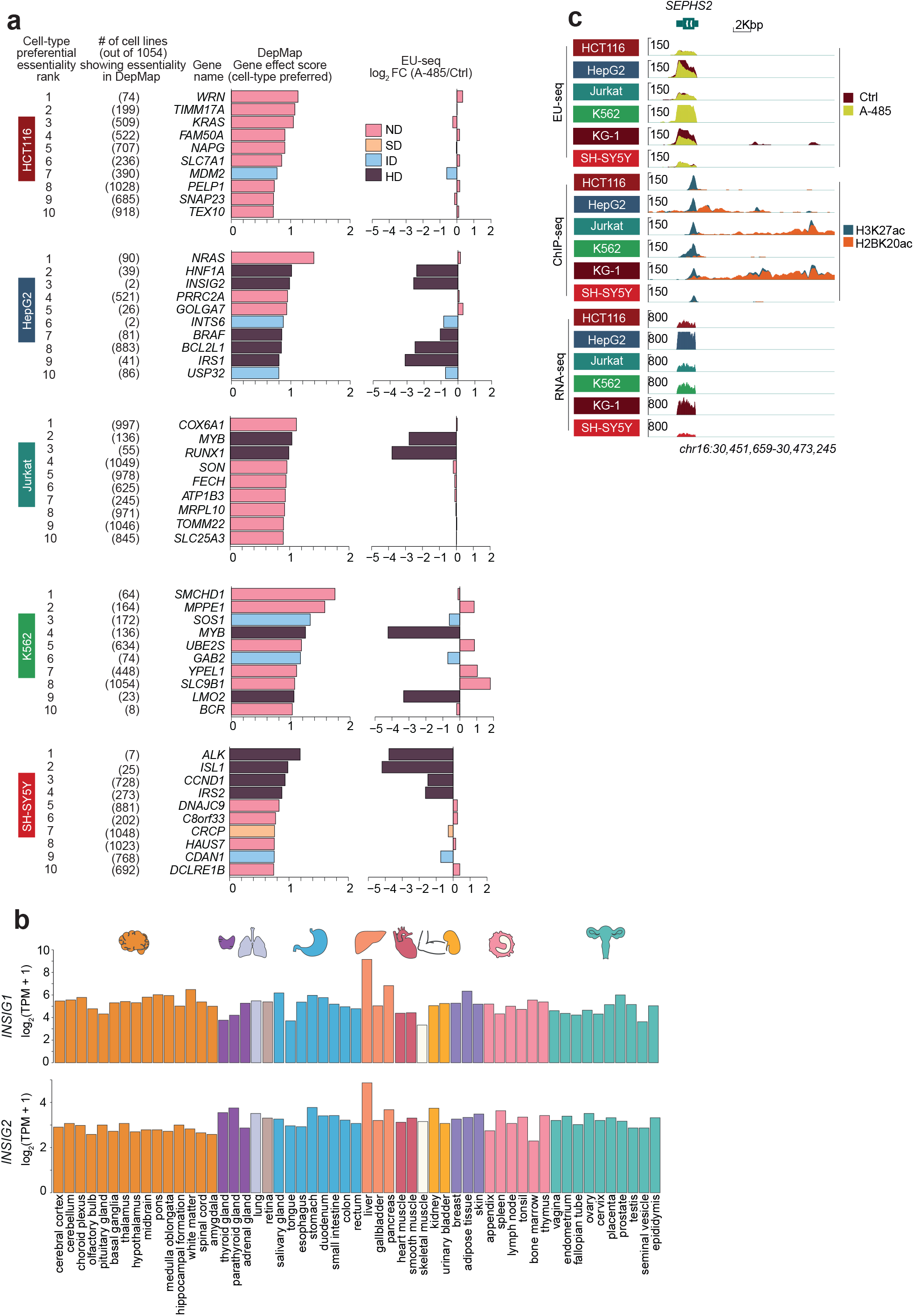
Expression profiles of *INSIG1* and *INSIG2,* and cell-type preferential essentiality of enhancer-regulated genes. a, Some enhancer-regulated genes score among the top cancer cell line preferential essential genes. The figure shows the Top10 cell-type preferential essential genes in the indicated cell cancer lines. For each gene, cell-type preferential essentiality rank, number of cell lines showing gene essentiality (out of 1054 cell lines), DepMap gene effect score, and A-485-induced nascent transcription downregulation in the respective cell lines, are indicated. Except for HCT116, all cell lines include 2-4 genes that are strongly enhancer-regulated and score among the Top10 essential genes in the respective cell lines. Most notably, *INSIG2* is essential in just 2 (out of 1054) cell lines. Cell-type preferential essentiality data were obtained from the DepMap CRISPR-based genome-wide essentiality database^45^. b, Shown are mRNA expression levels of INSIG1 and *INSIG2* across the indicated human tissues. Gene expression data was obtained from the human protein atlas^36^. c, Genome browser tracks showing the expression and A-485-induced regulation of *SEPHS2* in the indicated cell lines.

**Extended Data Fig. 13.**
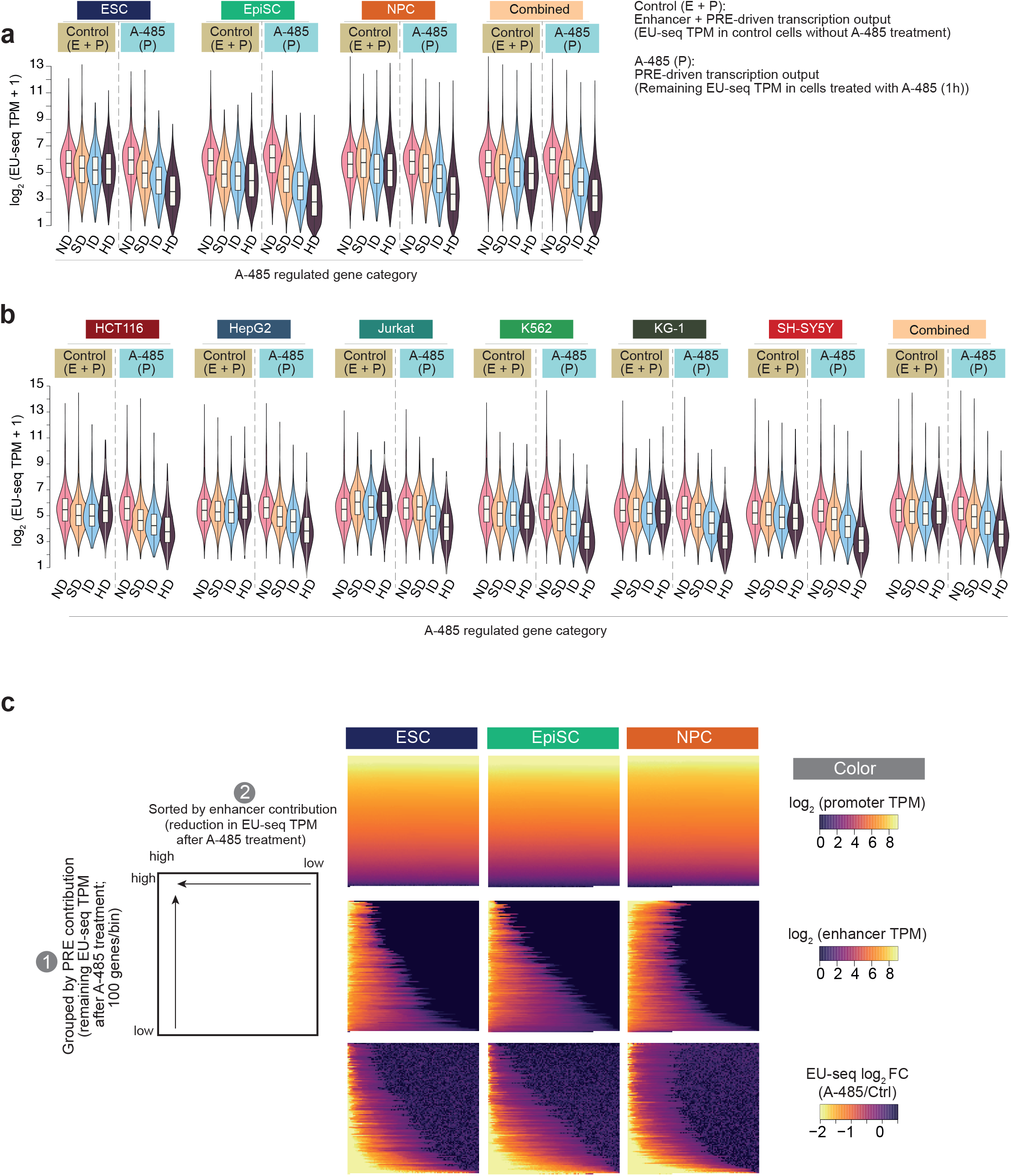
Enhancers and PREs, individually or in combination, drive gene expression over a large dynamic range. **a-b**, Nascent transcription levels in mouse (a) and human (b) cell lines. Shown are EU-seq TPM values for the indicated A-485 regulated gene class (ND, SD, ID, and HD), in the specified cell lines. In the untreated (Control) condition, gene expression is driven by the combined strength of enhancer (E) and PRE (P). After A-485 treatment, the contribution of enhancers to gene activation is ablated, and the remaining nascent RNA expression levels indicate promoter output sustained by PRE activity. In the control condition, gene expression levels in ND, SD, ID, and HD categories are roughly similar, albeit with some variability across cell lines. In each category, gene expression values vary by more than 3 orders of magnitude. After A-485 treatment, SD and ID genes are partially downregulated, and their expression is sustained at a much higher level than that of HD genes. Because CBP/p300 activity is globally inhibited, the remaining gene expression of SD and ID genes is sustained enhancer-independently. This shows that a large portion of genes is only partially activated by enhancers, and upon CBP/p300 inhibition, their remaining expression level is sustained by PRE activity. c, Absolute and relative contribution of enhancers differs for different genes. Gene expression in ESC, EpiSC, and NPC was quantified using nascent transcription analyses (EU-seq, TPM). Contribution of enhancer and PRE to gene expression was determined as described above. Genes were ranked from lowest to highest expression based on the PRE-driven gene expression level (i.e. remaining gene expression level after A-485 treatment), and binned (100 genes in each raw). Within each bin, genes were sorted based on the contribution by enhancers. Contribution of enhancer to gene expression is determined by the reduction in EU-seq TPM after A-485 treatment. Note that genes with relatively low contribution of PREs, show greater fold-change after A-485 treatment, indicating a relatively greater contribution of enhancers to genes with weaker PREs.

**Extended Data Fig. 14.**
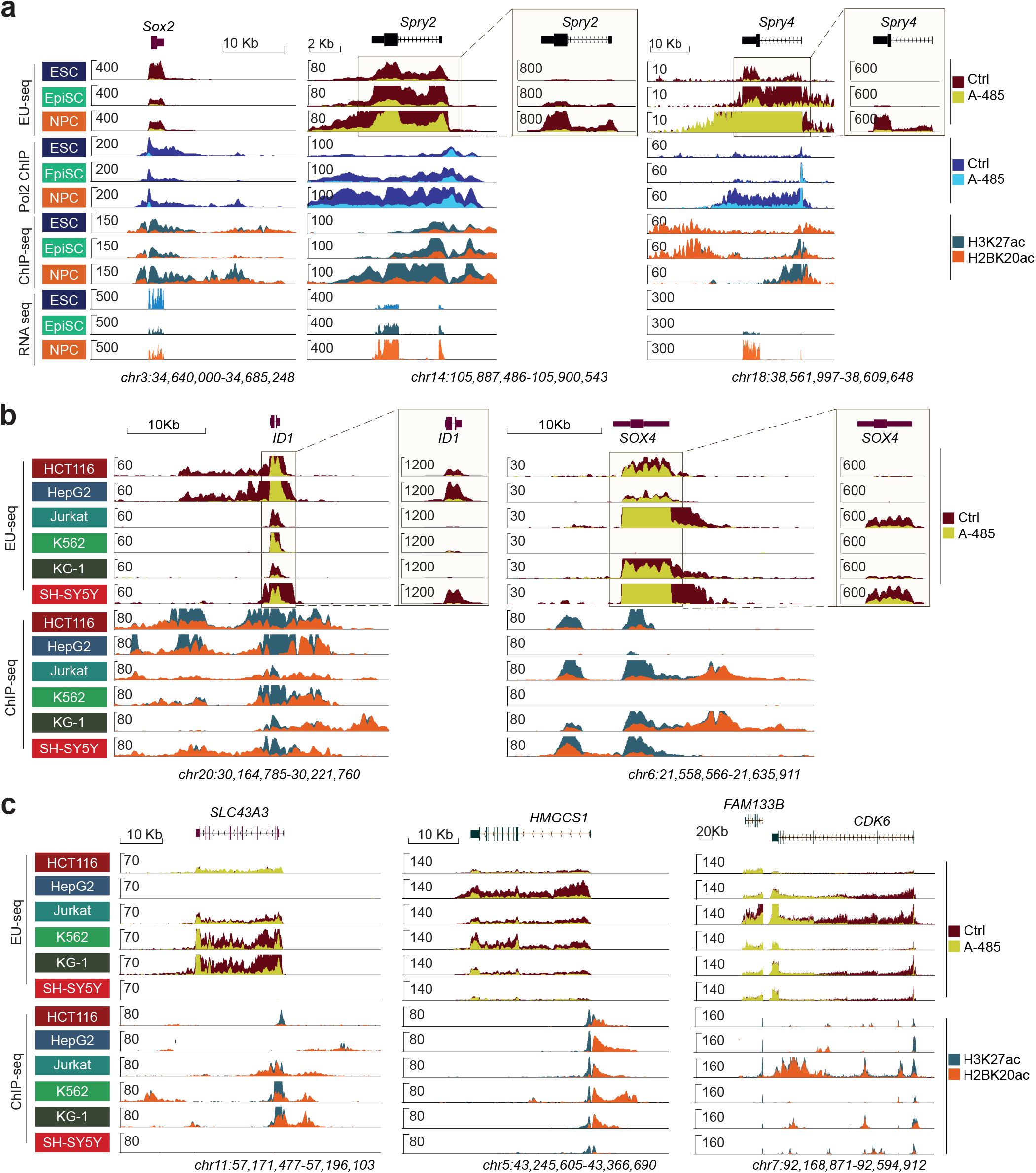
Enhancer strength is variable across different cell types. **a-c**, Shown are tracks of representative genes exhibiting very high variation in enhancer-dependent gene expression in mouse **(a)** and human cells **(b-c).** Note that genes are highly downregulated in most of the cell lines in which they are expressed, yet their expression levels vary greatly across the cell lines, demonstrating that gene-specific enhancer strength is highly variable across cell lines. Because of a very high dynamic range of expression, *Spry2, Spry4, ID1,* and *SOX4* expression is shown at two different scales. *FAM113B,* located downstream of *CDK6,* is only boosted by an enhancer in Jurkat, likely through the upstream enhancer located in this cell line.

**Extended Data Fig. 15.**
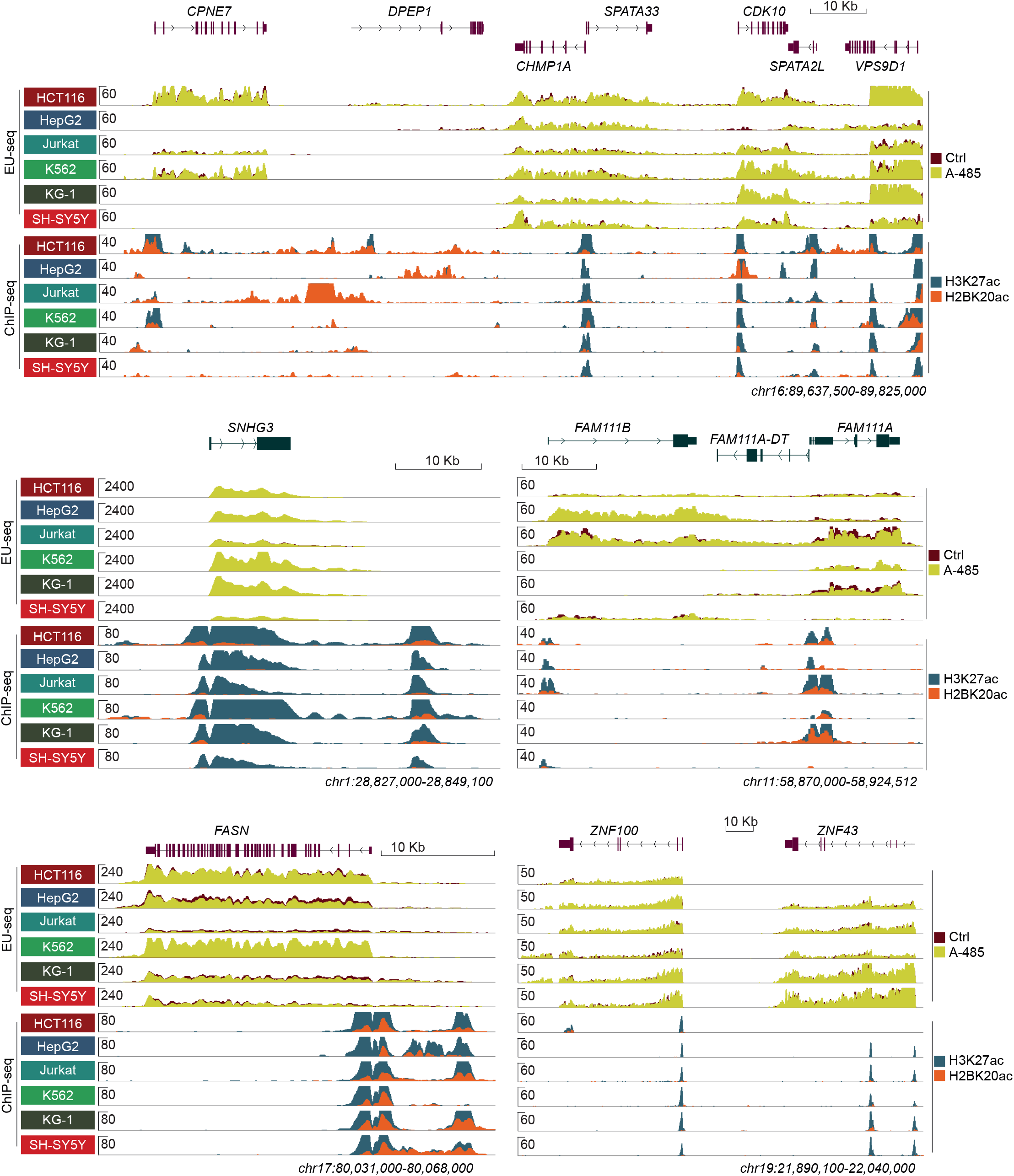
Cell-type-specific quantitative gene expression variation through alteration in PRE strength. Genome browser view of representative human genes showing cell-type-specific quantitative expression variation. Note that genes are expressed to variable levels in diverse cell lines, but they show no or only minor downregulation by A-485, demonstrating that their expression is quantitatively adjusted enhancer independently, likely through alteration of PRE strength.

**Extended Data Fig. 16.**
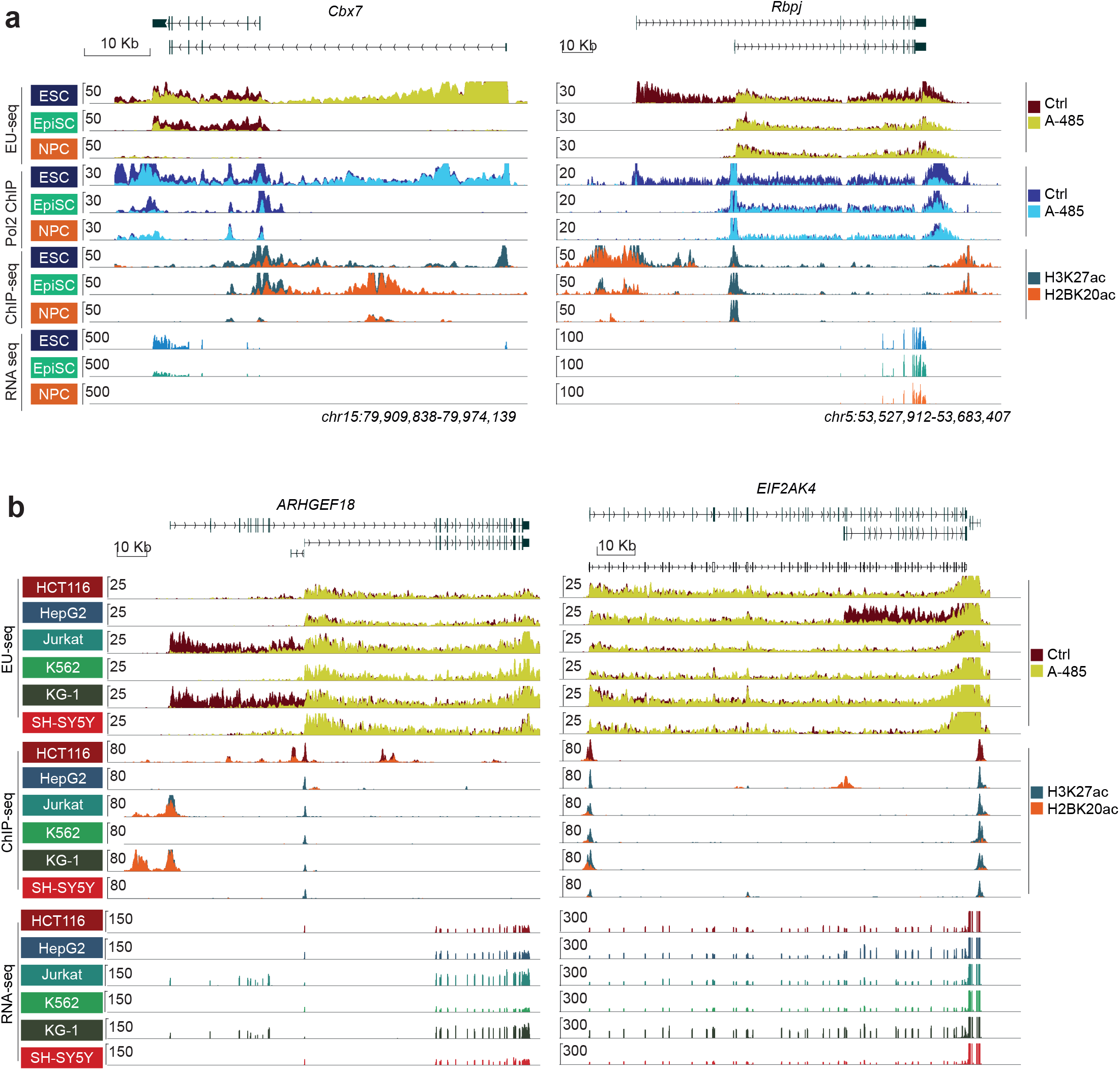
Enhancers and PREs differentially activate alternative promoters of the same genes. **a,**Genome browser view of representative human genes showing differential activation of alternative promoters by enhancers and PREs. *ARHGEF18* and *EIF2AK4* are expressed in all cell lines, but each of them expresses cell-type-specific isoforms. The promoter of the longer isoform of *ARHGEF18* is activated by an enhancer, whereas the promoter of the shorter isoform is activated by a PRE. Conversely, the promoter of the longer isoform of *EIF2AK4* is activated by PRE, whereas the shorter isoform is activated by an enhancer in HepG2. **b,** Differential activation of alternative promoters of the same genes in mouse cell lines. In ESC, *Cbx7* is expressed from two alternative promoters, whereas only the shorter isoform is expressed in EpiSC. The ESC-specific longer isoform of *Cbx7* is transcribed enhancer-independently by PRE, whereas the shorter isoform is activated by enhancers in both the cell lines. Of note, the remaining *Cbx7* expression in A-485 treated ESC likely reflects the EU-seq signal from the longer isoform of *Cbx7*. *Rbpj* is also expressed by two alternative promoters in ESC, but EpiSC and NPC only express the shorter isoform. While the promoter of the ESC-specific longer isoform is activated by an enhancer, the promoter of the shorter isoform is activated by PRE in all cell lines.

**Extended Data Fig. 17.**
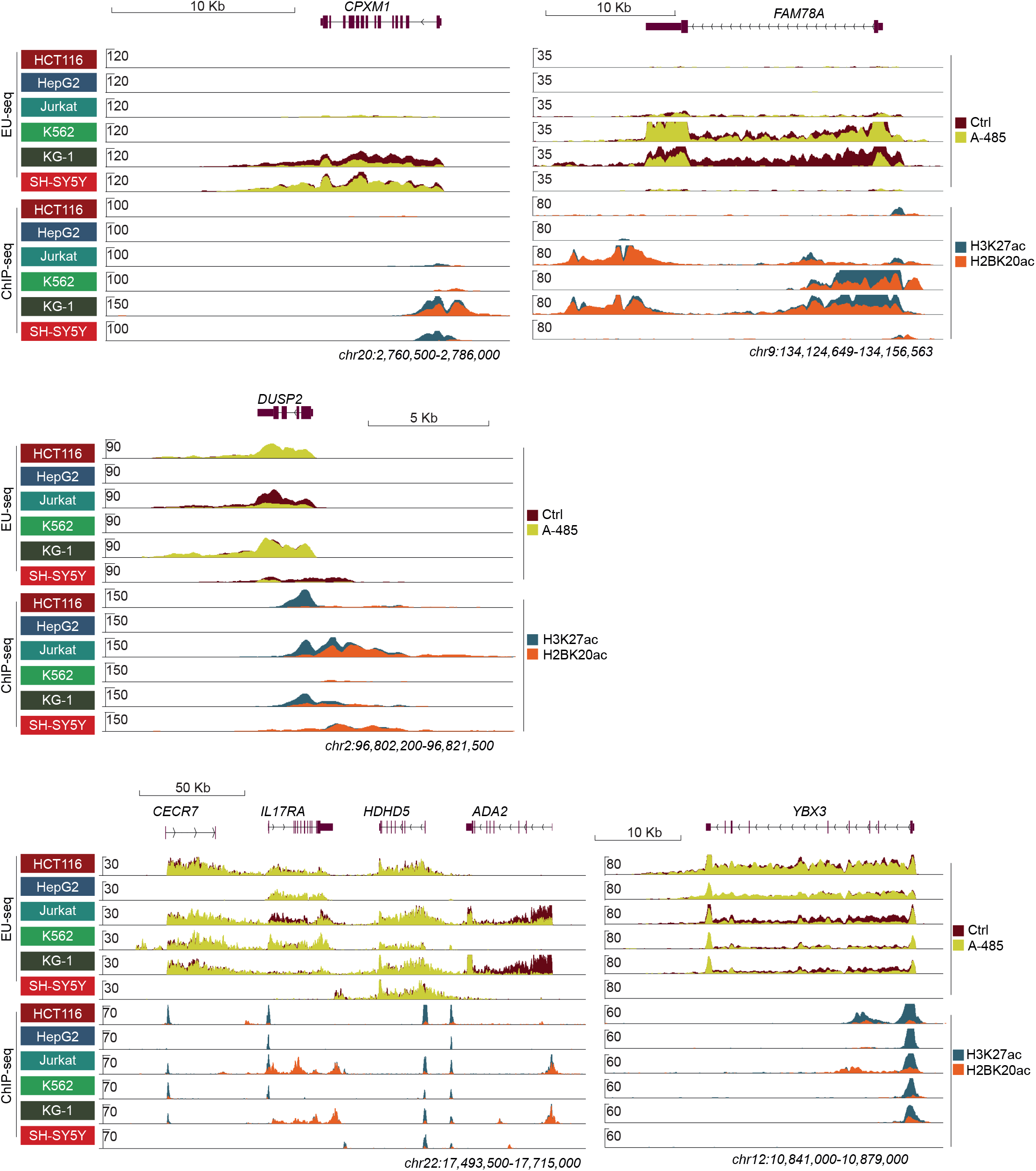
Combined strengths of enhancers and PREs afford quantitative expression variation of cell-type-specific genes. Genome browser view of representative human cell-type-specific genes showing quantitative expression variation through cell-type-specific tuning of the strength of enhancers, PREs, or both. Because of the variable strength of PREs and enhancers, expression of the genes is downregulated to a variable extent by A-485. There is a notable agreement between the occurrence of proximal H2BK20ac marked enhancers and A-485 sensitivity.

**Extended Data Fig. 18.**
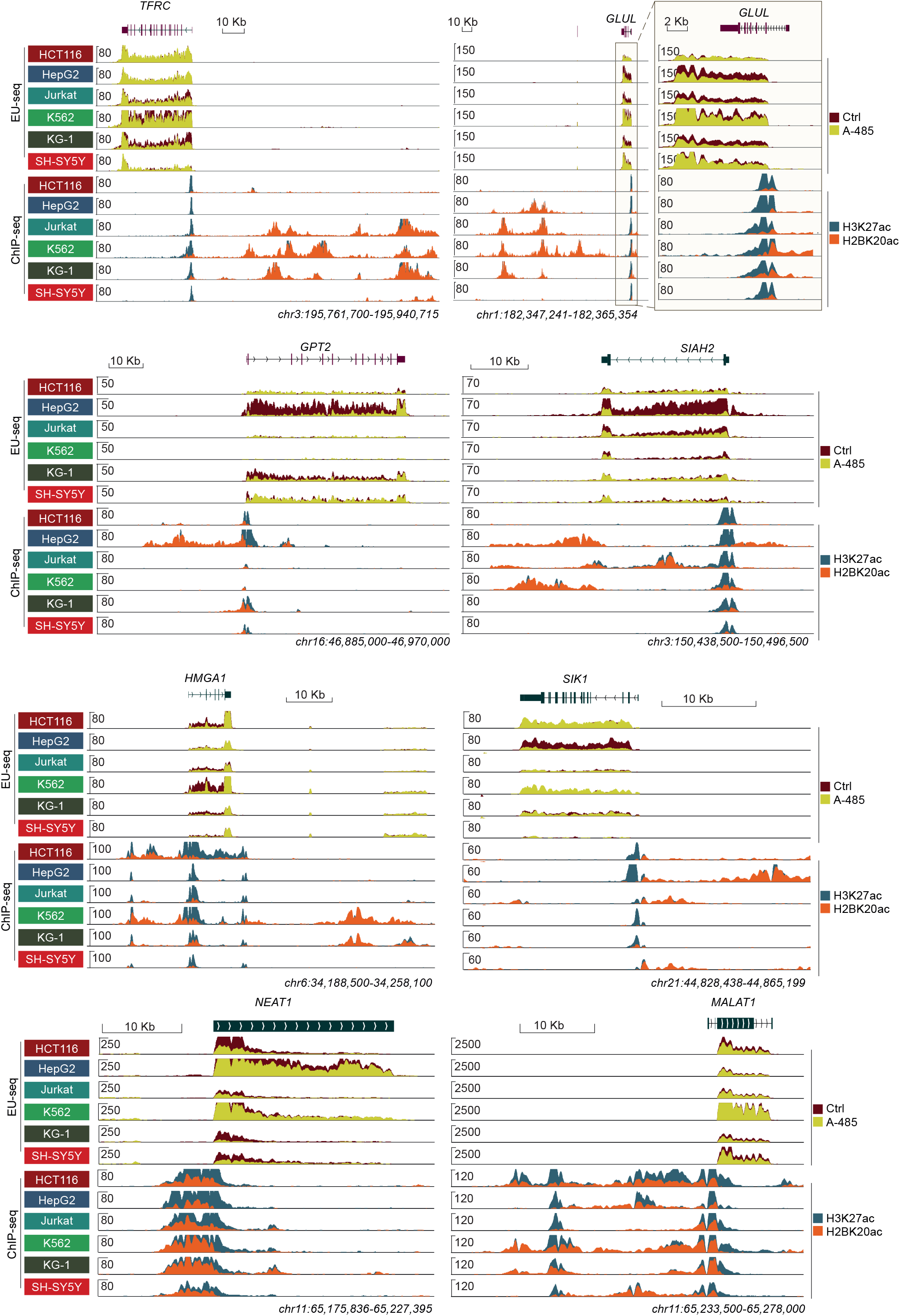
Tunable and integrated strengths of enhancers and PREs afford cell-type-specific quantitative expression variation of widely expressed genes. Genome browser view of representative human genes that are expressed in all the cell lines analyzed, yet show large quantitative expression variation. The expression of these genes is apparently altered through cell-type-specific tuning of the strength of enhancers, PREs, or both. The top and middle panels show genes (*TFRC, GLUL, GPT2, SIHA2, HMGA1, SIK1*) that are expressed at relatively low-to-modest levels. Their cell-type-specific quantitative variation and cell-type-specific regulation by enhancers are apparent in the reduction of nascent transcription in the A-485 treated condition. The bottom panel shows genes (*MALAT1*, and *NEAT1*) that are expressed at a high level. Enhancers also appear to contribute to their expression, but depending on the strength of PREs in individual cell lines, the measurable impact of enhancers is highly variable, and in some cell lines it is barely noticeable. This shows that in genes that harbor very strong PREs, the measurable relative impact of weak and modest enhancers can be extremely small.

**Extended Data Fig. 19.**
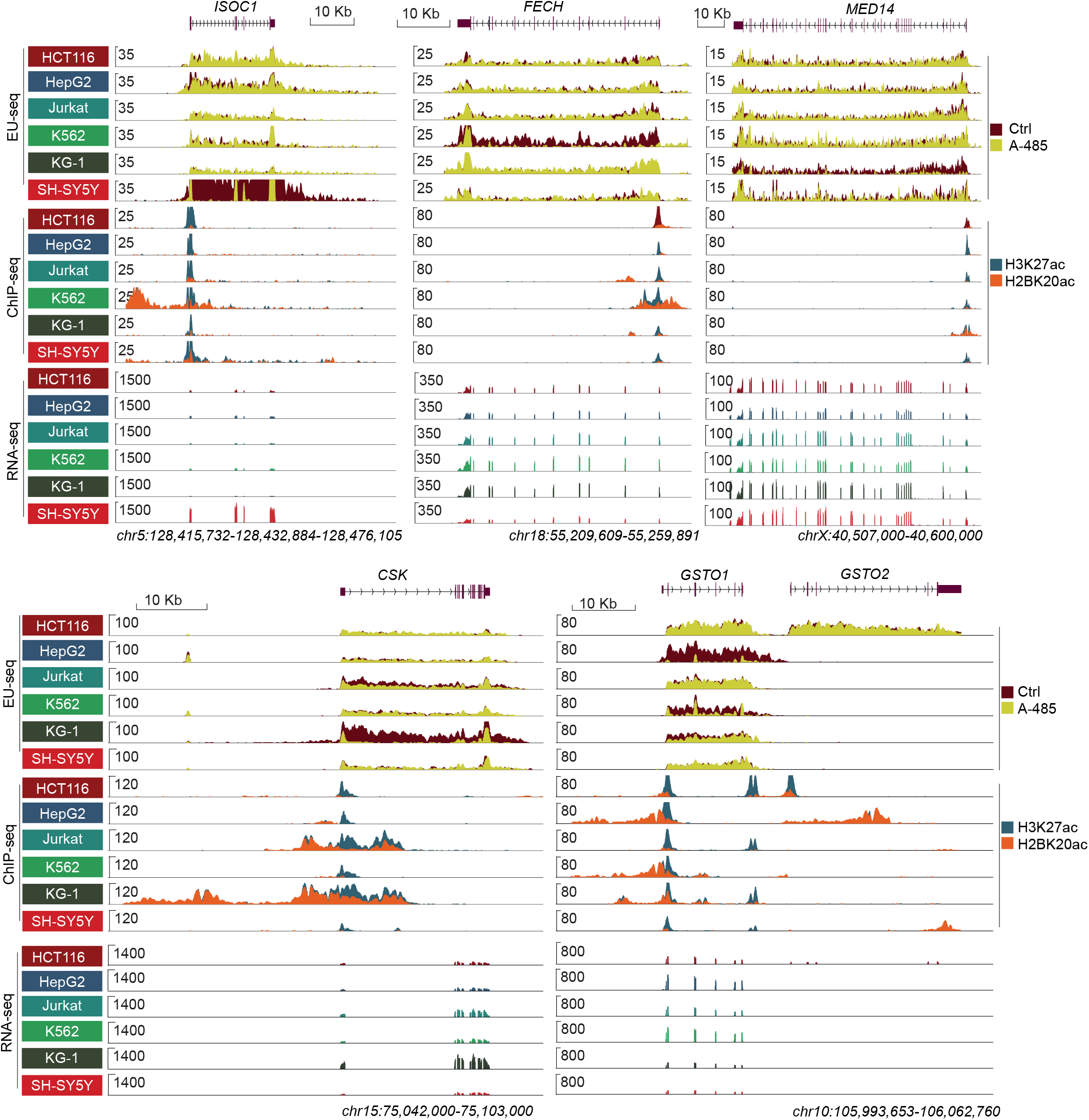
PREs and enhancers can interchangeably activate the same genes in different cell types. Genome browser tracks showing cell-type-specific, promoter-to-enhancer type gene activation switching. A-485 treatment abrogates *ISOC1* expression in SH-SY5Y cells, yet has no appreciable impact on any of the other cell lines. Expression of *FECH* and *MED14* is almost abrogated after CBP/p300 inhibition in K562 and KG-1, respectively, yet in the other cell lines, the genes remain unaffected by the inhibitor. Notably, the expression of *FECH* and *MED14* is similar in all cell lines, but the type of regulation differs among the cell lines. *CSK* is partially downregulated by A-485 in Jurkat, its expression is ablated in KG-1, but the expression is unaffected in the other four cell lines. Similarly, *GSTO1* expression is abrogated by A-485 in HepG2, remains unaffected in HCT116, K562, and SH-SY5Y, and is partially downregulated in KG-1 and Jurkat. Of note, the proximally occurring gene, *GSTO2*, is cell-type-specifically expressed, yet not regulated by A-485. Remarkably, all genes showing PRE-to-enhancer switching are marked with the strongest H2BK20ac signal, suggesting that gene regulation switching occurs as a consequence of the cell-type-specific acquisition of enhancers.

**Extended Data Fig. 20.**
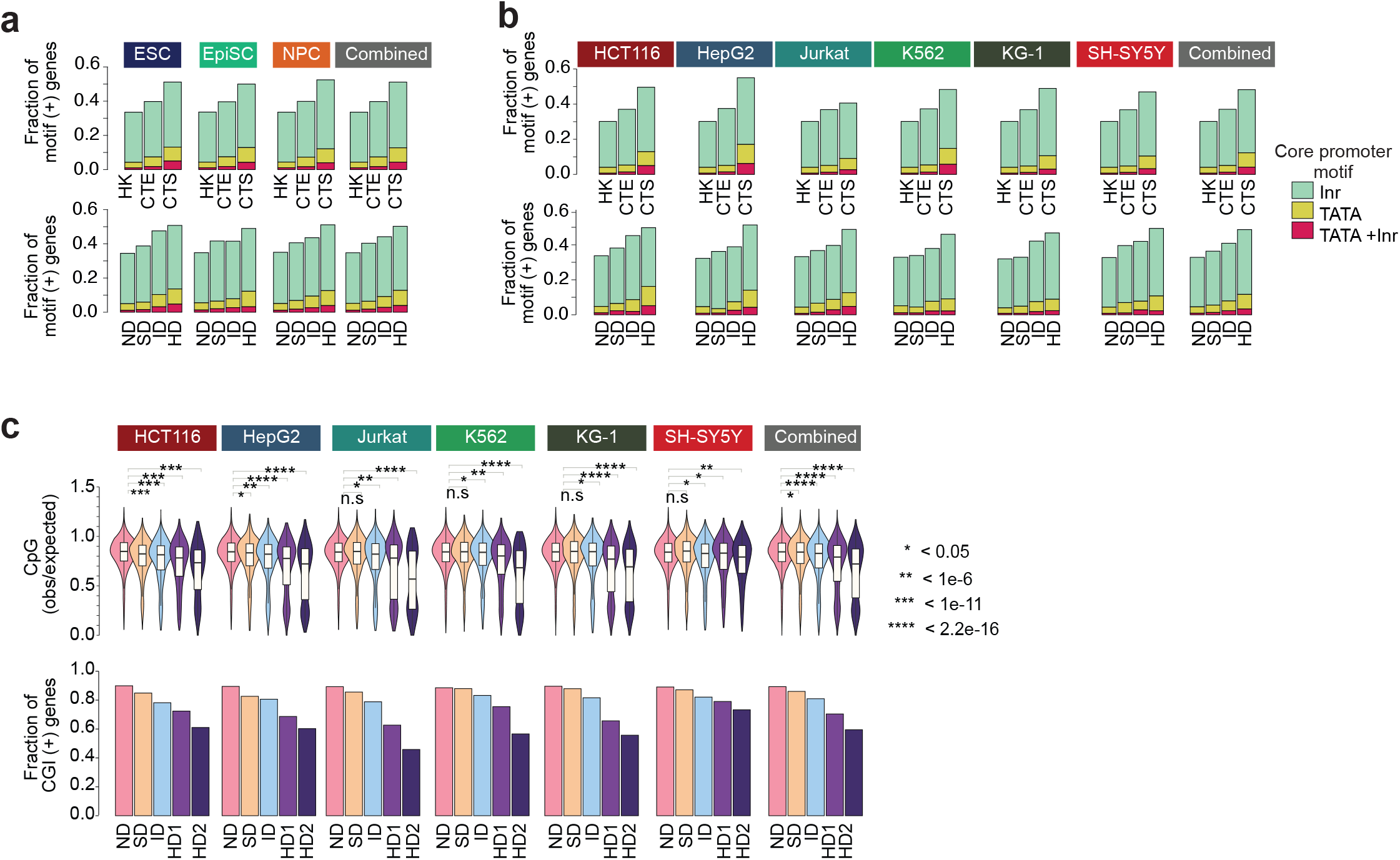
A greater fraction of enhancer-dependent promoters harbor TATA box, and autonomous promoters contain higher CGI density. **a-b**, Fraction of promoters containing the indicated core promoter motifs, in the specified cell lines. Genes were grouped based on cell-type-specificity (a) or A-485 downregulated gene (b) categories, and the fraction of genes harboring the indicated core promoter motifs are shown. Across different cell-type-specificity and A-485 regulated gene categories, the fraction of Inr motif-bearing promoters is relatively similar. The fraction of genes with TATA-box increases with increasing cell-type-specificity (HK<CTE<CTS) and with increasing A-485-induced gene down-regulation (ND<SD<ID<HD). However, overall, in all categories, only a small fraction of genes harbor a TATA box. c, Shown is the observed to expected (O/E) ratio of CpG dinucleotides (top panel) and the fraction of CpG island (CGI) positive promoters (bottom panel) and in the indicated A-485 regulated gene categories and specified cell lines. CpG O/E ratio was calculated using the sequence of ±250bp from TSS. CGI positive genes are defined as GC content > 50 % and CpG O/E ratio 0.6 in the ±250bp regions from TSS. * p < 0.05, ** p < 1e-6, *** p < 1e-11, **** p < 2.2e-16, Mann–Whitney U test.

**Extended Data Fig. 21.**
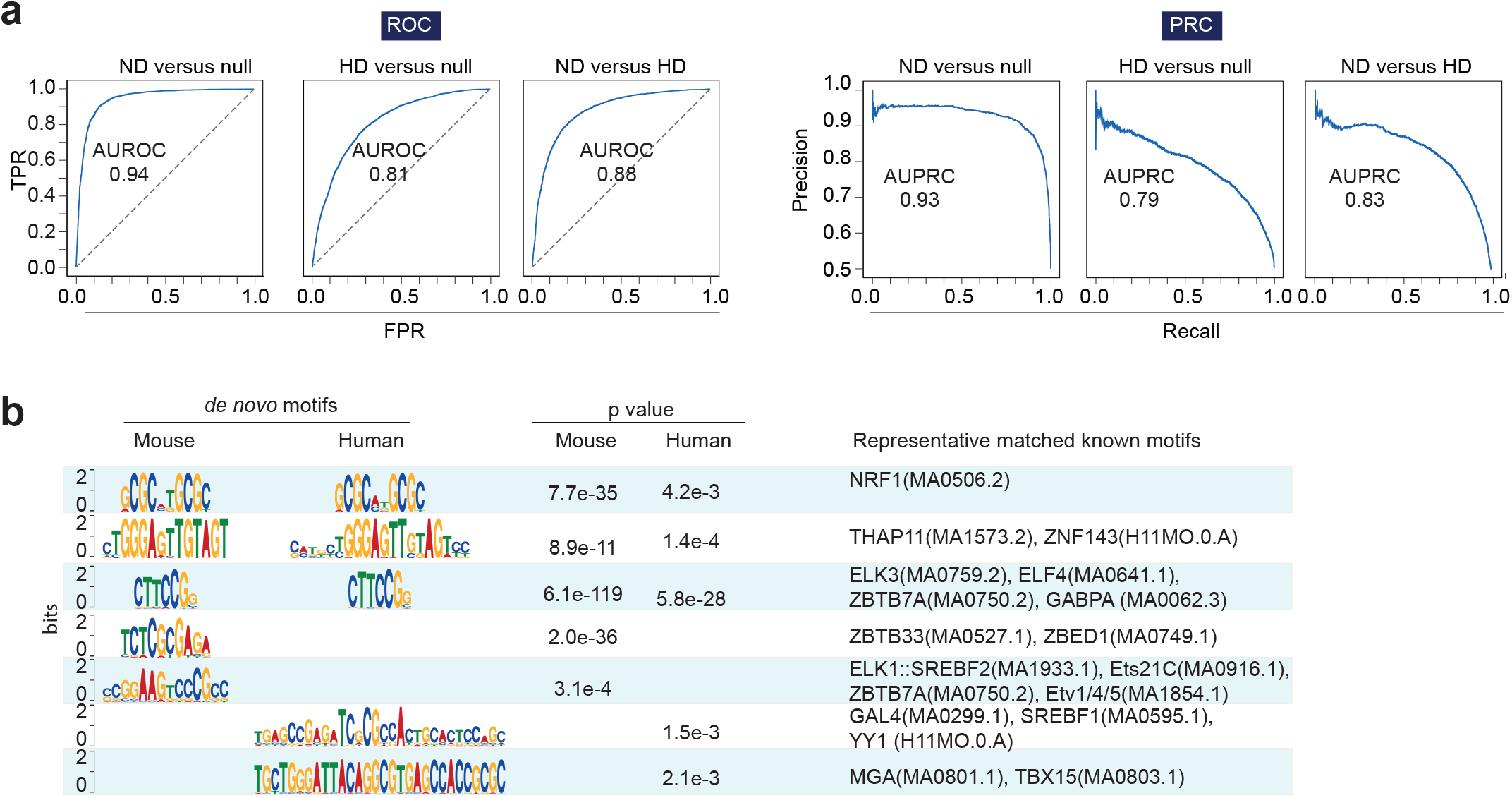
PRE sequences discriminate between autonomous and enhancer-dependent promoters. **a,**Gapped k-mer support vector machine (SVM)-based kernels (gkm-SVM) can discriminate ND and HD promoters from each other, and null DNA sequences. The receiver operating characteristic curve (ROC, top panel) and precision-recall curve (PRC, bottom panel) show the accuracy of the gkm-SVM classifier for discriminating ND and HD promoters from null sequences, as well as for discriminating ND promoters from HD promoters. Notably, ND promoters are discriminated from null sequences with greater accuracy than HD promoters. ND promoters can be discriminated from HD promoters with similar accuracy as for HD promoters to null sequences. gkm-SVM kernels were trained using DNA sequences, upstream 500bp from TSS of ND (n= 3258) and HD (n= 3154) gene promoters from 6 human cell lines, or randomly selected repeat- and GC-matching sequences from the same genome (see methods for details). AUROC and AUPRC are shown. **b,** Shown are the de novo sequence motifs represented in the importance scores calculated by gkm-SVM discriminating ND promoters from HD promoters. Representative transcription factors with matching DNA binding motifs are shown. The enrichment of each motif in ND promoter over HD promoter (upstream 500bp from TSS) is tested using a two-sided fisher test and the calculated p values are shown.

**Extended Data Fig. 22.**
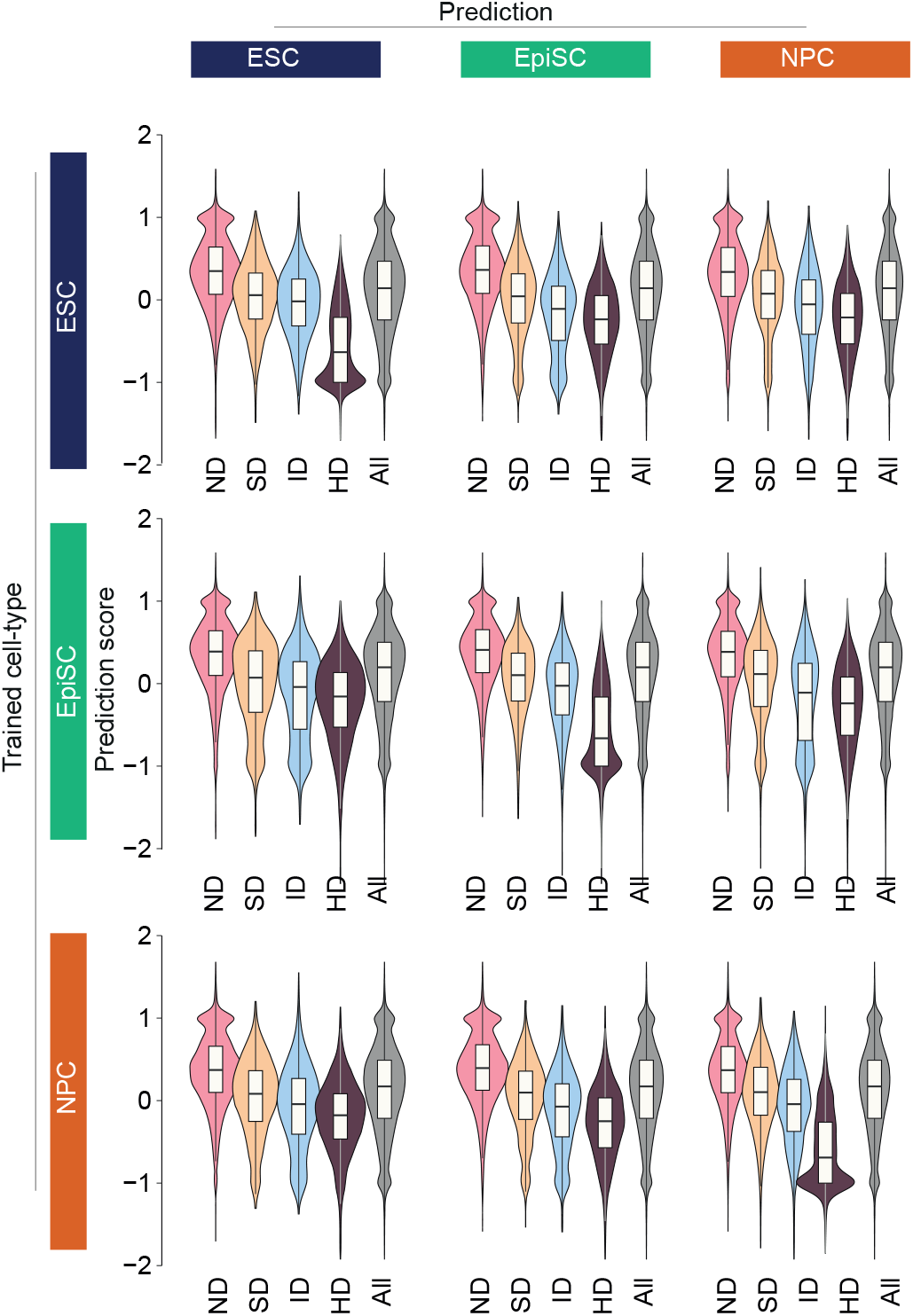
Promoter sequence difference discriminate enhancer-dependence of genes. The classifier was trained using ND and HD genes exclusively downregulated in one of the indicated cell lines and performance was tested on the same cell line, as well as in the other two held-out cell lines.

**Extended Data Fig. 23.**
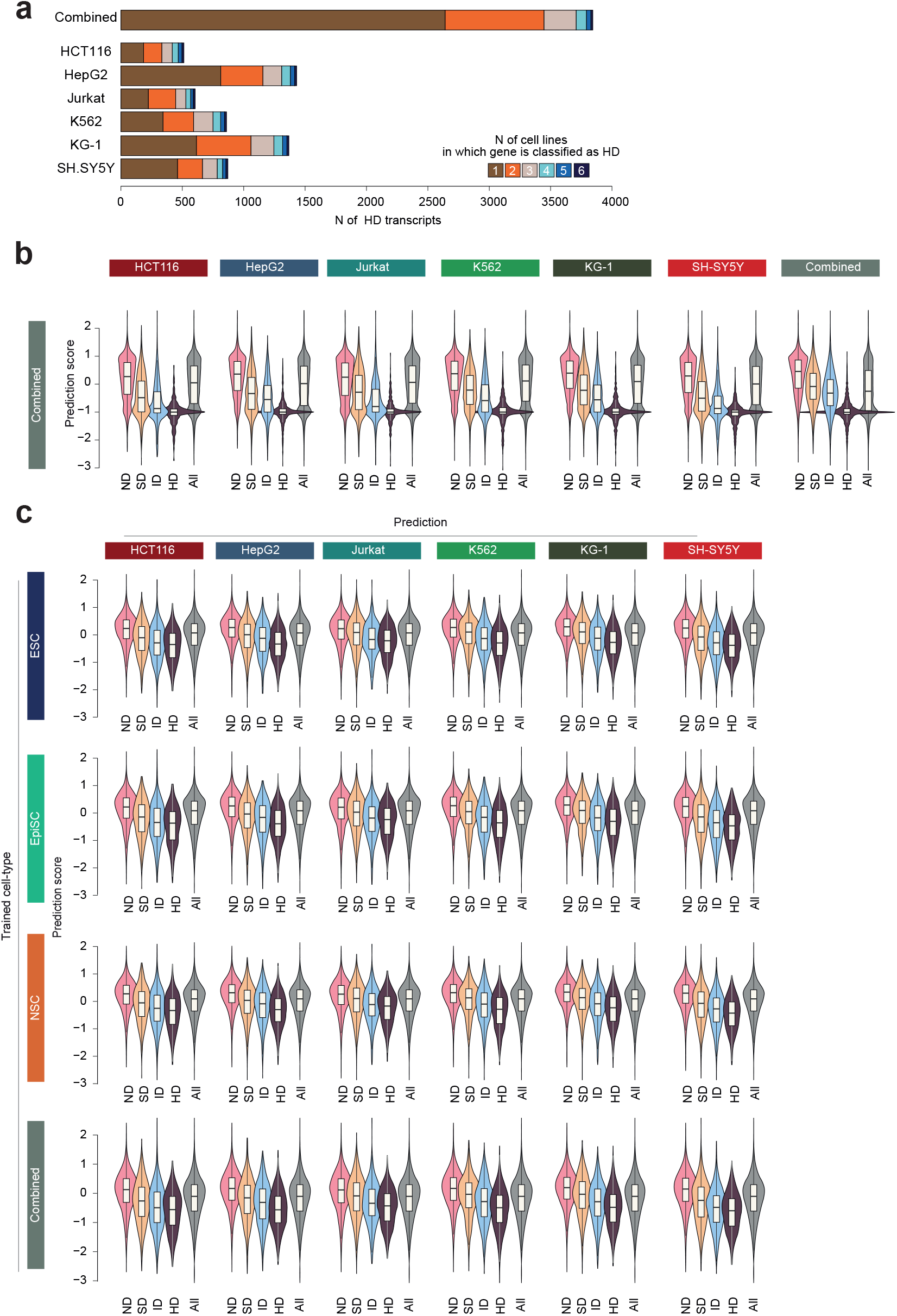
Mouse and human ND and HD promoters share sequence features. **a,** The overlap of human HD genes between different cell-types. The bar chart shows the number of HD transcripts on each human cell lines and the number of overlapping with other cell lines are shown with individual colors. **b-c,** A gkm SVM model was trained using ND and HD genes as positive and negative sets, respectively. For training the model in human cells **(b)**, ND and HD genes combined from all the human cell lines were used. Separately, a model was trained using ND and HD genes from one of the indicated mouse cell lines **(c)**. For both the models, trained in human or mouse cell lines, performance was tested on each of the indicated human cell lines.

